# CMG2 interaction with actin is required for growth factor-induced chemotaxis in endothelial cells

**DOI:** 10.1101/2025.07.21.665964

**Authors:** Fangfang Jiang, Raj Kumar Mongre, Jacob Truman, Raphaela de Oliveira, Ethan Hardy, Elliot Hardy, James Fowler, Chao Wang, Hsien-Jung L. Lin, Zachary J. Rose, Ryan T. Kelly, P. Christine Ackroyd, Michael S. Rogers, Kenneth A. Christensen

**Affiliations:** Department of Chemistry and Biochemistry, Brigham Young University, Provo, Utah, United States of America 84602; Department of Surgery, Vascular Biology Program, Children’s Hospital Boston, Harvard Medical School, Boston, Massachusetts, United States of America 02115

**Keywords:** CMG2, F-actin, chemotaxis, migration, primary endothelial cell, angiogenesis

## Abstract

Capillary morphogenesis gene 2 (CMG2/ANTXR2) is a cell surface receptor that contributes to corneal angiogenesis and is essential for orienting endothelial cell chemotaxis in growth factor gradients. However, the mechanism by which CMG2 transmits signals to orient endothelial cells is unknown. Here, we use affinity proteomics to identify proteins that interact with CMG2. This led us to investigate CMG2 interaction with F-actin in the context of serum and individual growth factors (bFGF/PDGF/VEGF) in both primary and immortalized endothelial cells. We also measured different matrix-protein-mediated changes in chemotaxis. We find that CMG2 function is matrix dependent, and that CMG2-actin interaction is required for serum- and growth-factor-induced chemotaxis. Inhibiting chemotaxis with CMG2 antagonists (collagen 6, penta-galloyl-glucose, or anthrax toxin protective antigen; PA) leads to the release of actin cytoskeleton from CMG2. Furthermore, CMG2 co-localizes with actin at the leading edge during live migration. This colocalization is disrupted by PA, which also disrupts directional movement. Finally, we demonstrate that the conserved actin binding domain of CMG2 directly binds to F-actin *in vitro* where it bundles F-actin, suggesting a mechanism by which it enables directional migration.

## Introduction

Angiogenesis is the formation of new blood vessels from pre-existing vasculature. The process is tightly regulated and involves endothelial cell activation, proliferation, migration, and tube formation [1]. While essential for physiological processes such as wound healing and tissue repair, dysregulated angiogenesis contributes to pathological conditions, including cancer, endometriosis, and corneal neovascularization, a leading cause of blindness [2]. Current anti-angiogenic therapies target vascular endothelial growth factor (VEGF) signaling, a key regulator of blood vessel formation [3]. However, despite their clinical efficacy, anti-VEGF-pathway treatments are associated with significant limitations, including adverse side effects, diminishing therapeutic response over time, and drug resistance [4,3]. These challenges underscore the need to develop novel anti-angiogenic strategies and identify alternative molecular targets.

One promising candidate is Capillary Morphogenesis Gene 2 (CMG2/ANTXR2), an integrin-like transmembrane protein implicated in angiogenesis [5,6]. CMG2 was initially identified as a novel gene upregulated during endothelial tube formation *in vitro* [7]. Recent *in vivo* studies have demonstrated that CMG2 contributes to corneal neovascularization, highlighting its importance in angiogenesis [5,8]. *In vivo,* genetic ablation of ANTXR2 in mice or pharmacological inhibition of CMG2 results in reduced angiogenesis [9]. *In vitro* studies using the immortalized endothelial-like cell line EA.hy926 demonstrates that CMG2 is essential for chemotaxis - a directional migration response pivotal for angiogenesis [5]. While wild-type cells migrate efficiently toward growth factors, CMG2-deficient cells or those treated with CMG2 antagonists lose this chemotactic capability but retain overall motility [5]. These findings collectively position CMG2 as a key regulator of endothelial cell migration, though the precise molecular mechanisms remain unclear.

The actin cytoskeleton is fundamental to angiogenic processes, orchestrating endothelial cell proliferation, migration, and remodeling of the extracellular matrix [5,10]. During sprouting angiogenesis, endothelial tip cells extend actin-rich filopodia at their leading edge, allowing them to sense, detect and respond directionally to gradients of guidance cues such as VEGF [11]. VEGF signaling through its receptors (VEGFRs) drives reorganization of the actin cytoskeleton and modulates focal adhesion turnover, generating the contractile forces required for chemotactic migration [12]. Notably, in the context of zebrafish development [13] and hyaline fibromatosis syndrome (HFS) [14] interactions between CMG2 function and the actin cytoskeleton have been suggested. But, despite these advances, the precise relationship between CMG2 and F-actin during angiogenesis remains poorly understood. In this study, we demonstrate that direct interaction between CMG2 and F-actin is essential for guided endothelial chemotaxis in a manner dependent on both growth factor signaling and extracellular matrix context.

## Results

### Blocking CMG2 reduces both HMVEC and *EA.hy926* endothelial cells migration *in vitro*

Our previous studies of *in vitro* migration using EA.hy926 cells revealed that blockade or knockout of CMG2 significantly impairs chemotaxis, suggesting that CMG2 regulates angiogenesis by controlling endothelial cell migration [5]. However, the role of CMG2 in primary human microvascular endothelial cell (HMVEC) migration remained unexplored. In addition, the CellAsic device that we previously used to measure chemotaxis was discontinued. To address these gaps, we investigated the migration of HMVECs toward serum, both in the presence and absence of CMG2 antagonists, using a new ibidi microfluidics device for migration assays. We found that HMVECs can migrate toward serum as shown in ***Fig. 1a,c&d*,** validating both the migratory capability of HMVECs and the reliability of the ibidi device for chemotaxis studies. We then measured the effects of the CMG2 antagonists PA and PGG on HMVEC migration. PA is a high-affinity CMG2 antagonist (Kd ∼ 0.2 nM) and PGG binds CMG2 and inhibits HMVEC migration with an IC_50_ of 0.8 µM in trans-well assays [5]. In the ibidi assays, we measured two parameters to evaluate cell migration, the distance travelled up the chemotactic gradient (chemotaxis), and forward migration index (FMI) which is the distance travelled up the gradient divided by the total distance travelled and indicates the strength of chemotactic effect. Treatment with either PA (2 nM) or PGG (10 µM) significantly reduced HMVEC chemotaxis and FMI toward serum (***Fig. 1b-d & Fig. S1***). Similarly, targeting CMG2 with PA or PGG in EA.hy926 cells dramatically impaired their serum-induced chemotaxis and FMI (***Fig. 1e,f* *& Fig. S1***).

**Fig 1:**
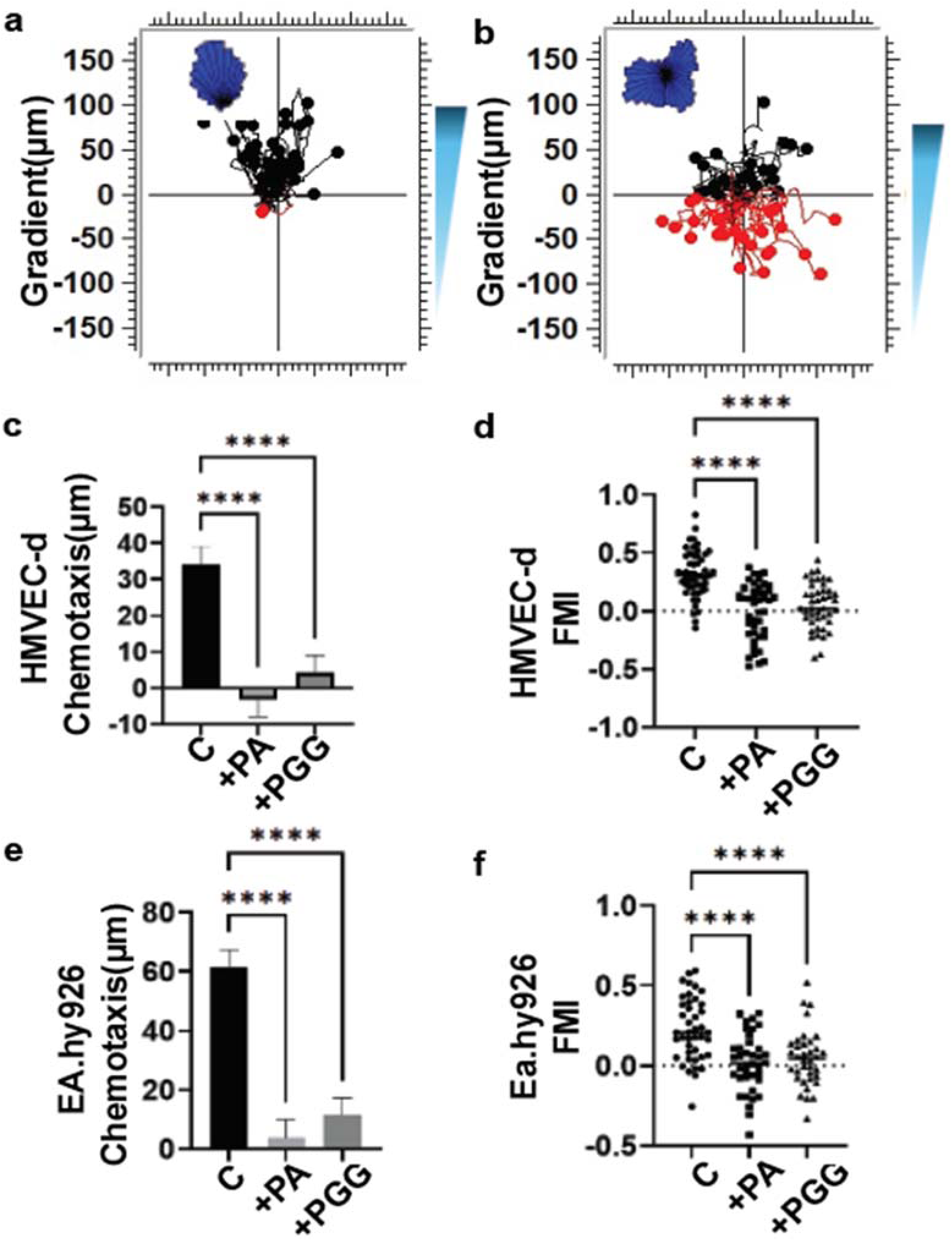
Blocking CMG2 in primary endothelial cells inhibits migration. **a-b:** Representative tracking plot with (**HMVEC-d**) or without (**b, HMVEC-d+PA**) chemotaxis behavior. **c-d:** Chemotaxis directness or FMI was measured by statistical analysis of trajectory plots in “Chemotaxis and Migration Tool V2.0”. **e-f:** Chemotaxis and FMI comparison between WT, PA or PGG treatment in EA.hy926 cells. Error bars are standard error of mean, *p<0.05; **p<0.01; ****p<0.0001 by one-way ANOVA.

### CMG2-actin interaction is required for CMG2-mediated migration toward the serum

To unravel the molecular mechanism(s) underlying how CMG2 regulates chemotaxis, we explored potential interactors involved in CMG2-mediated signaling. To this end, we transduced EA.hy926 cells with a lentiviral expression vector for V5-tagged CMG2. Since CMG2 expression is under a doxycycline-inducible system, we titrated CMG2 expression and found that 250 ng/mL doxycycline induced a 1.5-fold increase in CMG2 expression (***Fig. S2***) and used that doxycycline concentration to perform data-acquisition-independent affinity-pull-down tandem mass spectrometry with the workflow described in ***Fig. 2a***. Using a volcano plot to compare proteins pulled down from cells treated with PA to untreated cells, we were struck by the remarkable decrease in CMG2 interaction caused by PA (***Fig. 2b***). This suggests that PA treatment disconnects CMG2 from a substantial protein complex. To better identify the function of proteins whose interaction with CMG2 was significantly altered by PA treatment were subjected to a stepwise gene ontology (GO) analysis [15–17], focusing on those whose interaction was most strongly changed by PA treatment. This led us to focus on two GO biological processes whose proteins were differentially associated with CMG2 upon PA treatment. In the case of “actin cytoskeleton organization”, 107 of 543 proteins associated with the GO process were differentially associated with CMG2 upon PA treatment, and in the case of “positive regulation of lamellipodium assembly”, 10/28 proteins were.

**Fig 2:**
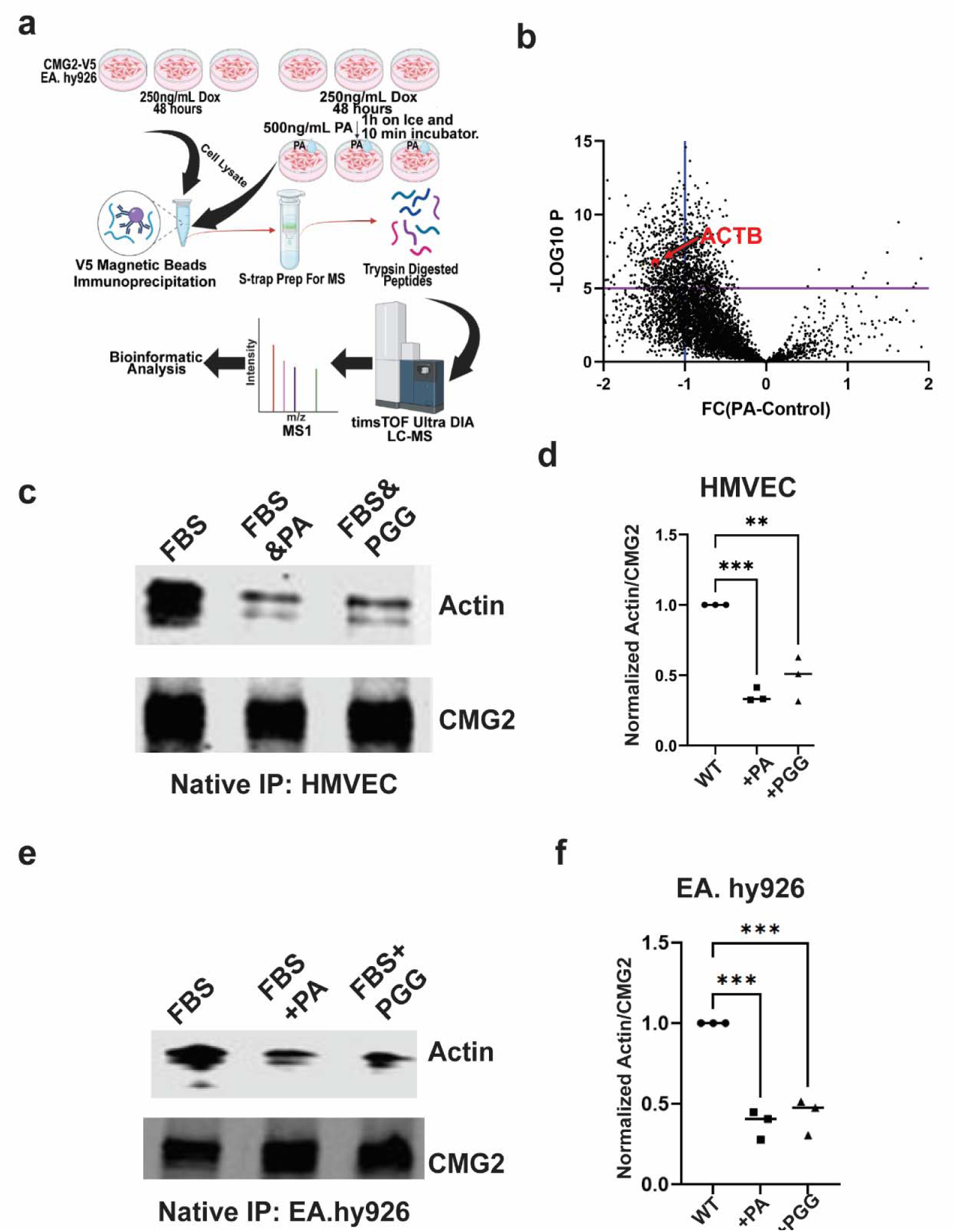
CMG2-actin interactions correlate with CMG2-mediated chemotaxis towards serum. **a:** Schematic workflow for affinity pull-down DIA mass spec. **b:** Volcano plot analysis from AP-MS with or without PA treatment. **c &e:** CMG2 and actin western blots of anti-CMG2 immunoprecipitation from HMVEC-d (**c**) or EA.hy926 (**e**) cells treated with PA or PGG. **d & f:** Quantitative analysis of IP-western blots performed as in c&e, showing the comparison of normalized actin/CMG2 between control, PA and PGG treated cells. Error bars are standard error of mean, *p<0.05; **p<0.01; ****p<0.0001 by one-way ANOVA in GraphPad Prism.

This enrichment of actin cytoskeleton components among putative interactions decreased by PA treatment is of particular interest because previous studies demonstrated relationships between CMG2 and actin in the context of anthrax intoxication [18], zebrafish spindle orientation [19], and the intracellular degradation of matrix proteins in HFS [20]. Given that F-actin is a well-established regulator of chemotaxis and directional cell migration [21,22], we hypothesized that CMG2 interacts with F-actin to mediate chemotaxis in endothelial cells.

To test this, we performed native pull-down assays using HMVECs cultured in complete ECM2. These experiments showed a robust interaction between CMG2 and F-actin. However, treatment with CMG2 antagonists PA or PGG significantly reduced this binding (***Fig. 2c,d***). Similar results were observed in EA.hy926 cells (***Fig. 2e,f***). Together, these findings support the idea that the CMG2-actin interaction is disrupted by CMG2 inhibitors that inhibit chemotaxis, providing a mechanistic link between CMG2 and cytoskeletal dynamics during migration.

### Multiple growth factors in serum facilitating CMG2-actin relationship promote chemotaxis

In the assays described above, we used molecules in serum as both the chemoattractant and migration substrate. But serum contains a complex mix of chemokines, growth factors, proteins, and hormones as well as migration substrates including fibrin and fibronectin [23]. To identify specific growth factors that use CMG2 for chemotactic signaling, we first focused on angiogenic growth factors. Based on previous migration assays with EA.hy926 cells, which demonstrated that bFGF, PDGF, and VEGF induce CMG2-dependent directional endothelial cell movement [5], we selected these growth factors for further investigation in HMVECs. We plated HMVECs on 10% complete DMEM media, which we then replaced with serum-free medium +/- growth factor to establish a growth factor gradient. Under these conditions, cells exhibited directional migration in response to bFGF, PDGF, and VEGF (***Fig. 3a-c* *& Fig. S3)***, confirming that these growth factors direct chemotaxis on a serum-coated substrate. But, the CMG2 antagonist PA abolished chemotaxis, indicating that CMG2 signaling is essential for growth factor-induced chemotaxis on serum-protein substrates (***Fig. 3a-c* *& Fig. S3***). To explore the mechanism underlying CMG2-dependent chemotaxis, we measured the impact of these growth factors on CMG2-actin interactions. Native pull-down assays in EA.hy926 revealed that bFGF, PDGF, or VEGF enhance the binding of CMG2 to F-actin. This increased interaction was significantly reduced upon PA treatment (***Fig. S4***) Furthermore, confocal imaging of fixed, CMG2-V5-expressing EA.hy926 cells revealed that treatment with bFGF, PDGF, or VEGF resulted in significant accumulation of colocalized CMG2 and F-actin at the cell periphery (***Fig. S5***). Strikingly, PA (protective antigen) abolished this growth-factor-induced colocalization (***Fig. S5***). These results indicate that CMG2-mediated migration of HMVEC on serum-protein substrates in response to bFGF, PDGF, and VEGF may depend on CMG2-F-actin interaction.

**Fig 3:**
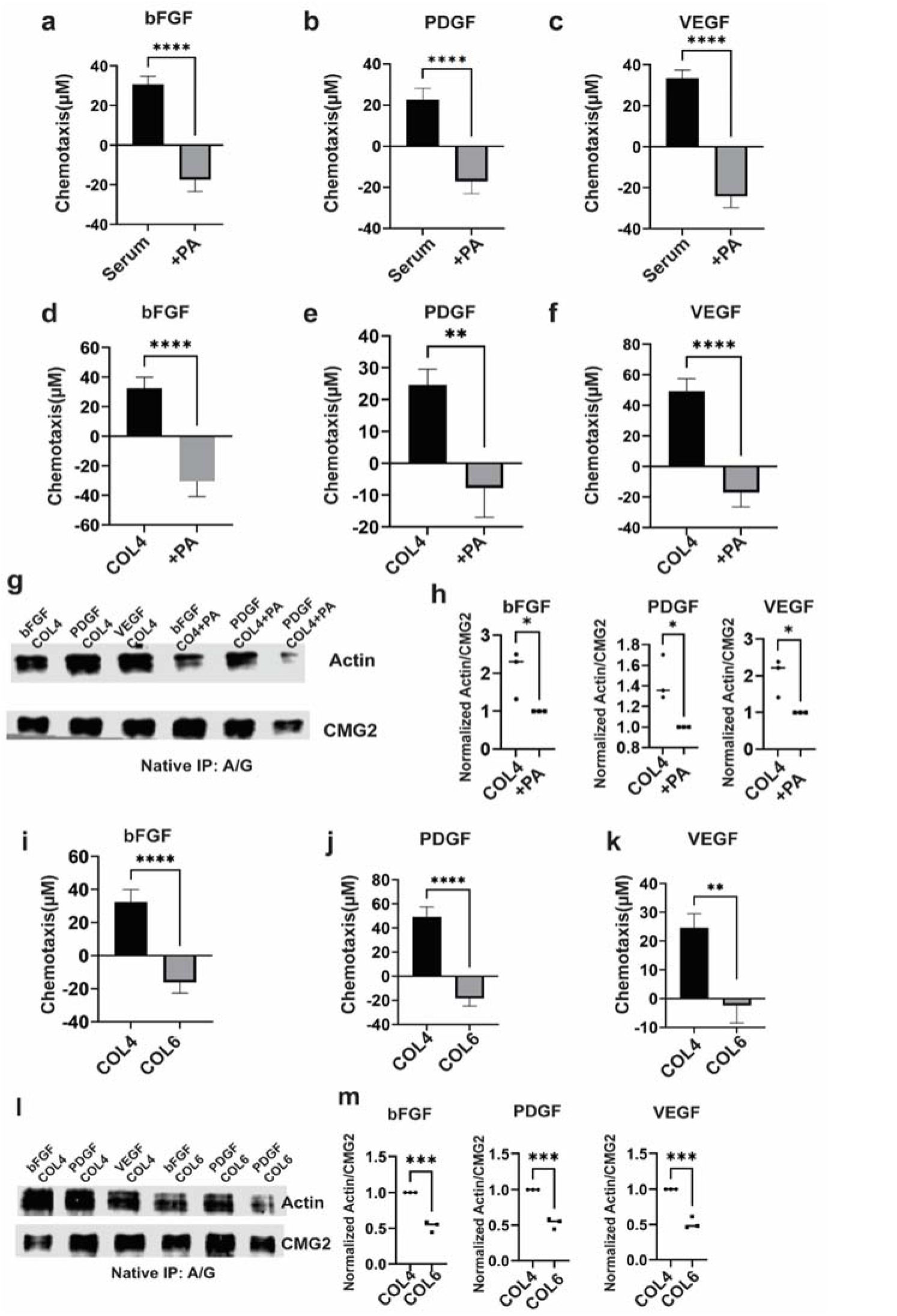
Individual growth factors increase actin interaction with CMG2. **a-f:** Chemotaxis directness toward 50ng/mL of bFGF, 50ng/mL VEGF or 50ng/mL PDGF of HMVEC-d plated on serum (**a-c**) or Col4 (**d-f**) in the absence of presence PA (n=40, 3 independent biological replicates). **g**: After overnight starvation, HMVEC-d cultured in basal media were treated with PA or vehicle control for 2 hours and then incubated with bFGF, VEGF or PDGF for an additional 6 hours. After anti-CMG2 immunoprecipitation, western blot against actin and CMG2 were analyzed and quantified **(h)**. **i-k:** HMVEC-d chemotaxis toward bFGF, VEGF or PDGF when plated on COL4 or COL6. **l:** CMG2-actin interaction as measured by co-IP western in HMVEC plated on COL4 or COL6 and treated with bFGF, VEGF or PDGF for 12 hours. **m:** Quantification of three independent experiments performed as in (l). Error bars are standard error of mean, *p<0.05; **p<0.01; ****p<0.0001 by unpaired t-test.

### Different matrix proteins effect on migration regulated by CMG2-actin interaction

In addition to growth factors, extracellular matrix (ECM) proteins present in fetal bovine serum (FBS) also play a significant role in angiogenesis by interacting with each other, growth factors, and/or cell surface receptors [24]. CMG2 binds with ∼0.5 μM affinity to several ECM proteins, including collagen IV (Col4), collagen VI (Col6), fibronectin, and laminin-111 [20,5]. Since Col4 is a major component of the basement membrane and plays a key role in endothelial proliferation, adhesion, and migration [25], we examined its role in the CMG2-mediated chemotaxis of HMVECs by plated them on Col4. We found that HMVECs chemotax well on Col4, but that blocking CMG2 with PA strongly reduced migration toward bFGF, PDGF, or VEGF on this substrate (***Fig. 3d-f* *& Fig. S3***). Native pull-down assays confirmed that PA treatment decreased CMG2-actin binding in HMVECs cultured on Col4 (***Fig. 3g&h***), supporting the idea that Col4 enhances CMG2-actin interaction in the presence of growth factors.

In contrast, Col6, a native ligand of CMG2, exhibited PA-like behavior. Unlike Col4, Col6 coating abolished HMVEC migration toward bFGF, PDGF, or VEGF (***Fig. 3i-k* *& Fig. S3***). Furthermore, native pull-down assays from HMVECs revealed that Col6 disrupted CMG2-actin binding, when compared to Col4 (***Fig. 3l&m***). Similarly, Col6 dramatically reduced EA.hy926 cells migration to bFGF, PDGF and VEGF and interfered with CMG2-actin binding compared to EA.hy926 plated on Col4 (***Fig. S6***). Together, these data demonstrate that CMG2 can mediate dramatically different responses to endogenous ligands, with Col6-like ligands inhibiting CMG2-mediated chemotaxis and Col4-like ligands supporting chemotaxis, and that interaction with actin correlates with CMG2’s chemotactic activity.

### CMG2 and F-actin colocalizes at the leading edge guiding directional migration

To better understand CMG2-actin dynamics during directional migration, we conducted live imaging of cells chemotaxing *in vivo*. We co-transfected EA.hy926 cells with mNeonGreen-tagged CMG2 and Lifeact, an F-actin marker. Time-lapse imaging of co-transfected EA.hy926 cells migrating in response to serum revealed that CMG2 (green) and F-actin (red) colocalized at the leading edge, driving the directional movement of endothelial cells (***Fig. 4a&c***). Treatment with PA disrupted the colocalization of CMG2 and F-actin at the leading edge (***Fig. 4b&d***), resulting in a loss of directional movement towards the serum (***Supplemental Video***).

**Fig 4:**
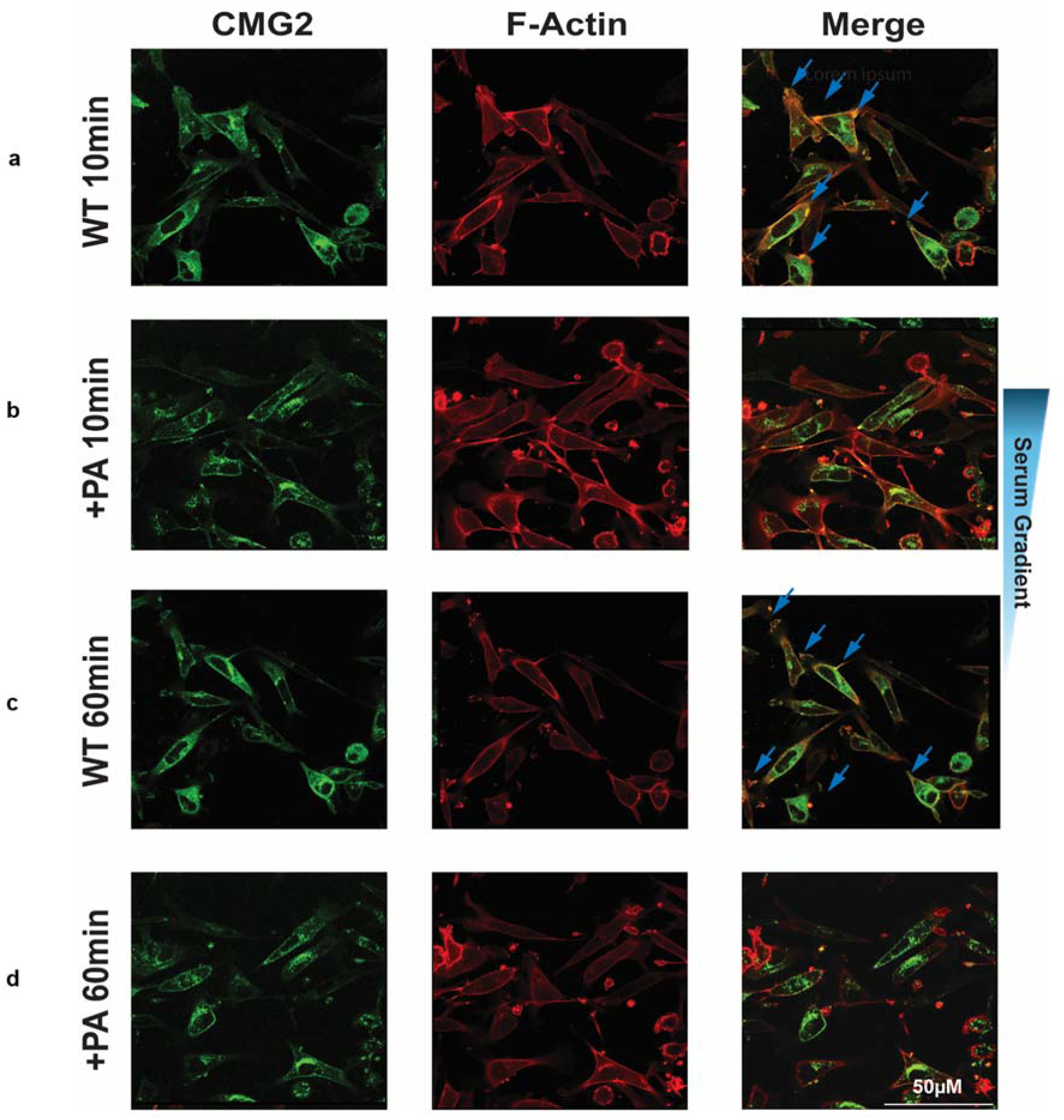
PA treatment disrupts CMG2-actin colocalization at the leading edge. EA.hy926 cells were co-transfected with CMG2-mNeonGreen and Lifeact (F-actin marker, m-Cherry), then plated on ibidi device in a 5% FBS chemoattractant gradient. Live recording timelapse of dynamic using Leica DMI8 between CMG2-MNG and F-actin with or without PA treatment. (Recorded movie is provided in supplemental). Arrow indicates the overlapping of CMG2 with F-actin.

To extend these results to primary cells, we seeded HMVEC-d cells in ibidi chemotaxis chambers in a serum gradient and then performed immunofluorescence staining for F-actin and CMG2. Interestingly both CMG2 and F-actin staining intensity was elevated in control cells as compared to cells treated with the CMG2 inhibitors PA or PGG (***Fig. 5a***). Furthermore, we noted that treatment with PA and PGG resulted in a noticeable reduction in staining for both F-actin and CMG2 at the leading edge (P<0.0001 by ANOVA, ***Fig. 5b,d***). However, CMG2 inhibitor treatment caused no significant reduction in F-actin staining at the trailing edge (***Fig. 5c***). However, we did observe a down regulation in CMG2 at the trailing edge (***Fig. 5e***), consistent with an overall reduction in cellular CMG2 as observed by western blot (***Fig. 5i***).

**Fig 5:**
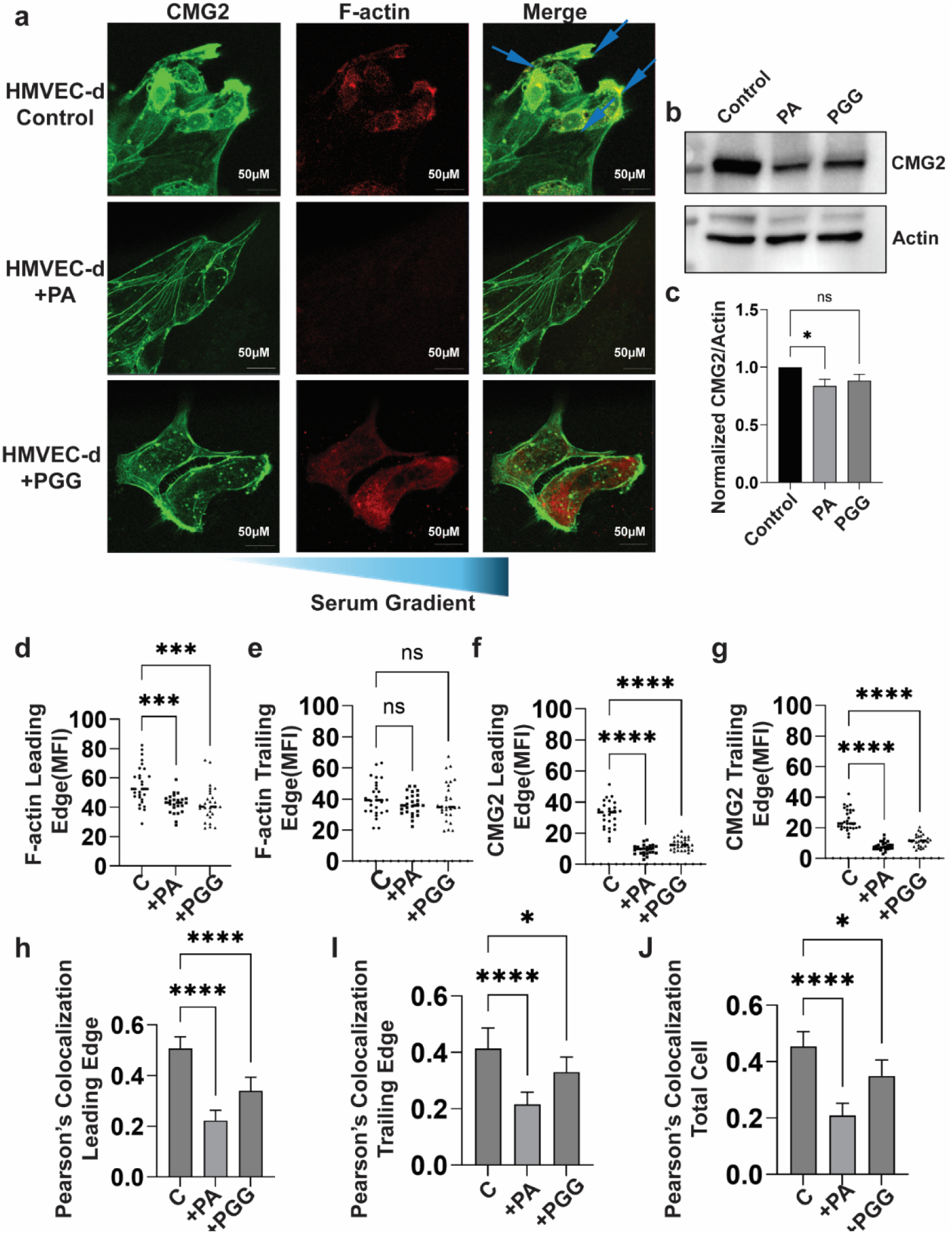
CMG2 and F-actin accumulation and co-localization at the leading edge of HMVEC-d in an FBS gradient is inhibited by CMG2 antagonists. **a:** Immunofluorescence staining showing colocalization of CMG2 (red) and F-actin (green) in chemotaxing HMVEC-d cells. Arrows indicate strong colocalization in the migrating front. Scale bar is 50 μm. **b-c:** western blot analysis of CMG2 protein abundance in control, PA and PGG treated HMVEC-d cell lysates. β-actin was used as a loading control. Data are representative of 7 sets independent experiments. (n = 7, ns P>0.05, *P < 0.05 by ANOVA followed by Dunnett’s post-test for control vs. PA and vs. PGG treated cells). **d-g:** ImageJ quantification of colocalization and staining intensity of F-actin and CMG2 in three independent experiments. Mean Fluorescence intensity (MFI) was measured in each cell from a standardized 264x389 pixel region of interest; MFI for individual cell’s leading and trailing edges was measured similarly. (n = 30, ns P>0.05, **P < 0.01, ***P < 0.001, ****P < 0.0001 by ANOVA followed by Dunnett’s post-test for control versus PA and PGG treated cells). **h-j.** Colocalization of CMG2 and F-actin in chemotaxing cells, as measured by Pearson’s correlation analysis using the Coloc2 function in the Colocalization Plugin in ImageJ, using standardized ROI as with b-e. Comparisons were performed using 2-way ANOVA with multiple Tukey’s posttest (*p<0.05; **p<0.01; ****p<0.0001).

To evaluate whether CMG2 might directly induce F-actin accumulation at the cells leading edge to promote endothelial migration, we measured colocalization of CMG2 with F-actin in chemotaxing HMVEC-d. We observed strong colocalization across the cell, particularly in the migrating front at the base of structures resembling chemotaxis-associated lamellipodia (***Fig. 5a***). As measured by Pearson correlation, PA treatment reduced the colocalization of CMG2 with F-actin across the cell, with particularly strong effects at the leading edge of HMVEC-d (***Fig. 5f-h***). These findings suggest that CMG2 plays a key role in regulating the accumulation of F-actin at the leading front of HMVEC-d, initiating directional migration and thereby enhancing chemotaxis.

### Conserved actin binding domain is essential for CMG2-actin interaction

Next, we investigated the specific region of CMG2 responsible for F-actin binding. Talin and vinculin have been reported to mediate the connection between F-actin and the CMG2 tail in the absence of ligand [20,26], suggesting that the CMG2 tail is critical for F-actin binding. To test this, we deleted the intracellular tail domain (CMG2^ΔITD^) and transduced the resulting construct into EA.hy926 cells. These cells failed to migrate towards serum chemoattractant (***Fig. 6a***). Furthermore, immunoprecipitation demonstrated that the interaction between CMG2^ΔITD^ and actin was completely abolished, confirming that the cytosolic tail of CMG2 is essential for F-actin binding (***Fig. 6b***). Interestingly, TEM8 (tumor endothelial marker 8), a paralog of CMG2 in mammalian cells, has been shown to directly interact with actin through a cytoplasmic sequence that is conserved in CMG2 [5]. To determine whether the homologous sequence has similar activity in CMG2, we deleted the putative actin-binding domain (CMG2^ΔABD^), spanning residues Lys366 to Pro419. We found that serum-induced migration of EA.hy926 endothelial cells was completely abolished in the absence of this sequence (***Fig. 6a***). Additionally, pull-down assays failed to detect any interaction between CMG2^ΔABD^ and F-actin (***Fig. 6b***), indicating that the actin-binding domain within the tail is necessary for CMG2-actin association.

**Fig 6:**
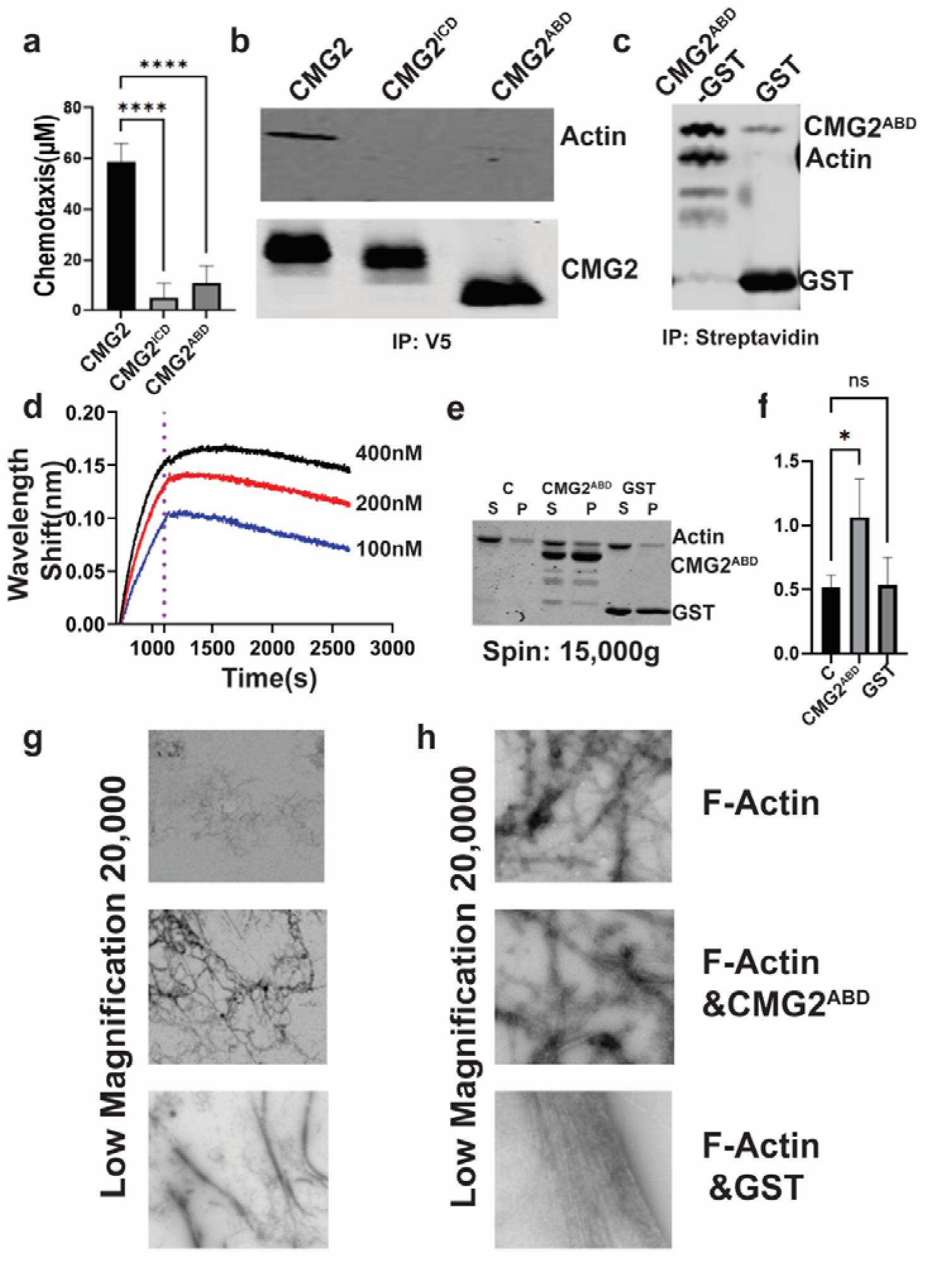
The CMG2 actin binding domain directly interaction with actin and facilitates F-actin bundling. **a:** Quantification of chemotaxis from CMG2 intracellular tail deletion (CMG2^ICD^), actin binding domain deletion (CMG2^ABD^) **b:** Anti-V5 bead pull-down followed by anti-actin and anti-CMG2 western blot. **c:** CMG2^ABD^ binds to F-actin derived from the EA.hy926 cell lysate. Coomassie blue staining of an avidin pull down of EA.hy926 cell lysate mixed with purified Avi-GST-CMG2^ABD^ or Avi-GST. **d:** Measurement of F-actin binding (400nM, 200nM and 100nM) to streptavidin coated biosensor coated with Avi-GST-CMG2^ABD^. **e:** Sedimentation assay to measure actin bundling activity. Coomasie blue staining of supernatant (S) and 15,000g pellet (P) of actin (C) mixed with Avi-GST-CMG2^ABD^ (CMG2^ABD^) or Avi-GST (GST). The relative mobility of the various components is indicted on the right. **f:** The ratio of actin in the pellet to actin in the supernatant in three independent experiments performed as in **(e)**. **g-h:** SEM images of F-actin filaments with F-actin alone, or with added Avi-GST-CMG2ABD. (CMG2ABD) or Avi-GST (GST). Error bars are standard error of mean. ns P>0.05, *P < 0.05, ****P < 0.0001 by ANOVA followed by Tukey’s post-test.

Consistent with these findings, confocal imaging revealed a stark contrast between CMG2-WT and CMG2^ΔABD^. While CMG2-WT showed clear colocalization with F-actin, CMG2^ΔABD^ failed to colocalize with F-actin (***Fig. S7***). Together, these experiments demonstrate that the actin-binding domain (ABD) within the CMG2 tail is indispensable for F-actin interaction and is required for the chemotactic migration of endothelial cells. These results emphasize the critical role of the CMG2 tail in mediating cytoskeletal interactions and directional cell movement.

### Direct CMG2-actin binding enables F-actin bundling *in vitro*

Lastly, we aimed to establish the biochemical function of the CMG2 ABD by determining whether this region of CMG2 binds directly to F-actin as is the case with its paralog, TEM8 [5]. We expressed and purified this domain with Avi and GST tags (Avi-GST-CMG2^ABD^) using a T7 bacterial expression system. We mixed Avi-GST-CMG2^ABD^ with EA.hy926 cell lysates and pulled-down the mixture with streptavidin and found that F-actin co-precipitates with Avi-GST-CMG2^ABD^ (***Fig. 6c***). This result indicates that the CMG2-ABD domain alone can interact with F-actin. To determine if this interaction is direct (i.e., not mediated by other proteins), we performed biolayer interferometry (BLI) to measure the binding affinity between purified Avi-GST-CMG2^ABD^ and purified actin. We found that Avi-GST-CMG2^ABD^ binds directly to F-actin with an estimated binding affinity of 10 nM (***Fig. 6d***).

To investigate whether CMG2-ABD can bundle β-actin into organized filaments like the homologous region of TEM8 [27,28], we conducted an actin sedimentation assay [29]. Centrifugation demonstrated that the presence of Avi-GST-CMG2^ABD^ significantly increased the amount of actin in the pellet (***Fig. 6e&f***), suggesting that CMG2-ABD promotes actin assembly, cross-linking, or bundling.

To distinguish between these mechanisms, we incubated Avi-GST-CMG2^ABD^ with actin and visualized the resulting filaments using negative-staining scanning electron microscopy (SEM). The resulting images revealed that Avi-GST-CMG2-ABD induced the formation of directional, side-by-side F-actin bundles (***Fig. 6g&h***). In contrast, actin alone or in the presence of Avi-GST control exhibited randomly clustered filaments without organized bundling (***Fig. 6g&h***). These findings suggest that CMG2-ABD directly binds to F-actin and facilitates its organization into parallel bundles, a process that likely plays a critical role in regulating cell migration. In summary, our data demonstrate that the CMG2-ABD domain is not only essential for direct actin binding but also promotes the formation of organized actin bundles, providing mechanistic insight into how CMG2 influences cytoskeletal dynamics and directional cell movement.

## Discussion

CMG2 plays a critical role in angiogenesis, but while its physiological role in anthrax toxin intoxication and hyaline fibromatosis syndrome (HFS) has been extensively studied [20], the detailed mechanisms by which CMG2 mediates endothelial cell chemotaxis remain poorly understood. This is true despite strong evidence supporting its indispensability in angiogenesis-mediating chemotaxis. Therefore, we investigated the signaling pathways involved in CMG2-mediated chemotaxis using HMVEC-d and EA.hy926 cells. We find that CMG2 directly interacts with F-actin *in vitro*, and this interaction is essential for directional cell migration in response to serum and growth factors such as bFGF, PDGF, and VEGF. Furthermore, the CMG2-actin interaction is modulated by ligands, including extracellular matrix proteins. Col4 promotes this association, facilitating chemotaxis, while PA, PGG, and Col6 inhibit it, leading to randomized cell migration. Importantly, we identified a conserved actin-binding domain (ABD) within the cytoplasmic tail of CMG2 that mediates direct F-actin binding and bundling. This CMG2-actin interaction localizes to the leading edge of migrating cells where it can drive directional movement. Despite these advances, the *in vivo* relationship between CMG2 and actin remains enigmatic. Investigating this interaction, particularly in a mouse model, could provide critical insights. Both CMG2 and its endogenous ligands, Col6 and Col4, are ubiquitously distributed in tissues [30]. Dominant-negative Col6 mutations are associated with collagen VI-related dystrophy, where patient-derived fibroblasts show CMG2 hyperphosphorylation, disrupting endosomal/lysosomal homeostasis and driving organelle accumulation [31]. Given our evidence that Col6 impacts CMG2-actin interactions, further exploration of the CMG2-actin complex in COL6-RD pathologies—using patient tissues or mouse models—is warranted. This would clarify whether CMG2 alone or its collaboration with actin regulates collagen VI homeostasis in fibrotic contexts.

Additionally, while our data strongly supports the role of growth factors like bFGF, PDGF, and VEGF in enhancing CMG2-actin interactions to facilitate endothelial chemotaxis, the molecular signaling connecting these extracellular signals to the intracellular CMG2-actin complex remains unclear. CMG2 is unlikely to act as a receptor for these diverse growth factors. Intriguingly, a 2011 study found CMG2 present in focal adhesion complexes [32], which are also regulated by the receptor tyrosine kinases from these growth factors, suggesting the possibility of a shared regulatory pathway. In this context, the impact of post-translational modifications, such as tyrosine phosphorylation and ubiquitination, on the CMG2-actin interaction in endothelial cells remains unexplored. PA and Col6, which disrupt CMG2-actin interactions, trigger tyrosine phosphorylation followed by ubiquitination [14]. Investigating how silencing these modifications affects CMG2-actin association could provide deeper insights into the regulatory mechanisms governing CMG2-actin signaling. Understanding these processes will not only enhance our knowledge of CMG2’s role in cell migration but also pave the way for novel therapeutic interventions targeting CMG2-related pathologies.

## Conclusion

Our study highlights the critical role of CMG2-actin signaling in endothelial cell migration and chemotaxis. Importantly, we discovered the actin-binding domain (ABD) of CMG2, which enables direct interaction with actin, facilitating CMG2-actin colocalization at the leading edge and promoting the bundling of actin filaments into organized, directional arrays during migration. This research provides a molecular mechanism explaining how CMG2 mediates endothelial cell migration in response to serum and growth factors (bFGF, PDGF, and VEGF). Interestingly, in the chemotaxis context, the endogenous CMG2 ligand Col6 behaves like the CMG antagonists PA and PGG, suggesting that differenced in ECM patterning may play an important role in guiding angiogenesis *in vivo*.

Looking ahead, future studies should focus on understanding how other protein partners interact with extracellular matrix proteins, growth factor receptors, and CMG2 antagonists like PA to trigger different intracellular signal transduction cascades. These signals likely regulate the activation or inhibition of the CMG2-actin complex, ultimately determining distinct migration behaviors under various conditions. Elucidating the interplay between actin cytoskeleton dynamics and the CMG2 complex is essential for fully understanding CMG2-mediated chemotaxis and its role in angiogenesis. Such insights hold significant promise for advancing antiangiogenic therapies and improving treatments for related diseases.

## Materials and methods

### Plasmid generation

To generate the ***CMG2-V5***, ***CMG2-mNeonGreen***, and ***CMG2^ΔITD^*** lentiviral transfer plasmids, sequences encoding CMG2-V5, CMG2-mNeonGreen, and CMG2^ΔITD^ were obtained from Twist Biosciences (South San Francisco, CA, USA) as entry clones flanked by attL sites in the pTwist ENTR Kozak backbone. These were Gateway cloned into pCW57.1 (a gift from David Root, Addgene plasmid # 41393), a tet-on vector containing compatible attR sites. For the LR recombination reaction, the following components were combined: 1 µL 50 ng entry clone plasmid, 1 µL 100 ng destination vector (pCW57.1), 1 µL LR Clonase II Enzyme Mix (Thermo Fisher, Cat. # 11791020) and 2 µL TE buffer. The reaction was incubated at room temperature for 16 hours, followed by the addition of 1 µL Proteinase K to terminate the reaction. The resulting recombinant plasmids were transformed into Stbl3 chemically competent cells (Thermo Fisher, Cat. # C737303) for selection and plasmid propagation.

To generate the ***CMG2^ΔABD^*** lentiviral transfer plasmid, the CMG2-V5 recombinant plasmid in pCW57.1 was subjected to Q5 site-directed mutagenesis using the following primer pair (Forward Primer: GAAGAGACAGAGGAGCCTATTAGGCCTAGA, Reverse Primer: CGGTTCTTCCTCCTCCTCTTTCGGAGCG) Following PCR amplification, the product was treated with the KLD reaction mix (NEB M0554S) to circularize the mutagenized DNA and digest the parental template. This step successfully removed the actin-binding domain (ABD), generating the CMG2^ΔABD^ expression construct. The final plasmid was transformed into Stbl3 competent *E. coli* (Thermo Fisher, Cat. # C737303), and colonies were screened to confirm the correct deletion.

The ***Avi-GST-CMG2^ABD^*** gene fragment was synthesized pET24 expression backbone. by Twist Bioscience. To construct the ***Avi-GST*** expression plasmid, we performed Q5 site-directed mutagenesis on the Avi-GST-CMG2^ABD^ vector using the following primers:( F: AAGCTTGCGGCCGC; R: CTATTTTGGAGGATGGTCGCCAC). The PCR product was treated with KLD enzyme mix to circularize the DNA and remove the template, thereby removing the CMG2 sequence. The resulting Avi-GST plasmid was transformed into NEB 10-beta competent E. coli (NEB) for selection. Positive clones were verified by colony PCR and sequencing.

### Cell Culture

EA.hy926 cells (ATCC, Manassas, VA, USA, Cat. # CRL-2922) were cultured in DMEM supplemented with 10% FBS and 1% penicillin-streptomycin at 37°C with 5% CO2 in a humidified incubator. Lenti-X 293T cells (Takara Bio, San Jose, CA, USA) were cultured in DMEM supplemented with 10% FBS and 1% penicillin-streptomycin at 37°C with 5% CO2 in a humidified incubator. Human Dermal Microvascular Endothelial Cells (HMVEC-d) were purchased from Lonza Bioscience (Walkersville, MD, USA, Cat No. CC-2543) and cultured in complete EGM™-2 Endothelial Cell Growth Medium-2 BulletKit™ (EBM-2 Basal Medium, Lonza Cat No. CC-3162) at 37 °C and 5% CO_2_. We used HMVEC-d at passage 4 to 7 for all chemotaxis and migration assays.

### Anthrax Protective Antigen Production

Anthrax protective antigen (PA) and the cleavage-resistant protective antigen mutant PA^SSSR^ were produced as described previously [5]. Briefly the proteins were expressed from pET-22b (RRID: Addgene 11079) into the periplasm and purified from the periplasmic lysate via anion exchange chromatography (Q-sepharose, GE Life Sciences Cat: 25236), using 20 mM Tris–HCl pH 8.0 with 20 mM NaCl (Buffer A) and Buffer A + 1 M NaCl. Endotoxin was removed by passing twice through poly-lysine coated cellulose beads (ThermoFisher, Waltham, MA, USA, Cat. # 88,275), followed by an endotoxin test using the gel clot method and appropriate dilutions.

### HMVEC-d and EA.hy926 Chemotaxis Assay

The endothelial cell migration and chemotaxis assay was carried out largely as described previously [5], but using uncoated (ibiTreat) μ-slide ibidi chemotaxis chambers [33] (ibidi USA, Fitchburg, WI, Cat. # 80326). We followed the 2D chemotaxis assay using the µ-Slide Chemotaxis for our migration setting (Application Note 17). Prior to performing HMVEC-d chemotaxis assay we freshly prepared attractant-free and chemoattractant medium. HMVEC-d were harvested by trypsinization at 80-90% confluence, and 3.0 × 10^6^ cells/mL were suspended in attractant-free EBM-2 containing 0% FBS and 0.1% bovine serum albumin (BSA). EBM-2 with 10% FBS and 0.1% BSA were used as a chemoattractant for migration. 7μL of the HMVEC-d suspension was gently seeded in the uncoated ibidi chemotaxis chamber and placed at 37 °C in a 5% CO_2_ incubator for at least 5 hours to allow cells to adhere. Then, left and right gradient chambers were filled with attractant-free EBM-2 medium then 15µL of chemoattractant EBM-2 medium (10% FBS, 0.1% BSA) was added twice to the right side of the chamber to make gradient after which all filling ports were closed with plugs.

For EA.hy926 migration in response to serum, following trypsinization, EA.hy926 cells were resuspended in DMEM supplemented with 10% FBS, loaded into an ibidi chamber, and incubated for 2 hours to allow adhesion. Attached cells were washed three times with serum-free DMEM containing 1% BSA, and the side chamber was filled with the same serum-free medium. To induce chemotaxis, DMEM with 10% FBS was added to the designated port as a chemoattractant. For inhibitor-treated conditions, 2 nM PA or 20 mM PGG was included in all media (cell suspension, wash buffer, and chemoattractant), while other steps remained identical. Migration was then monitored and analyzed.

For growth factor-mediated migration assay on serum coated chambers, trypsinized cells were resuspended in complete DMEM (10% FBS, 1% Pen/Strep) with or without 2 nM PA, and 7 µL of cell suspension was loaded into an ibidi chemotaxis chamber followed by 2-hour incubation (37°C, 5% CO₂) for adhesion. Cells were washed three times with EBM-2 containing 1% BSA, and both side reservoirs were filled with the same medium. Chemoattractants: 50 µM VEGF (ThermoFisher: 100-20-100UG), 50 µM bFGF (ThermoFisher: 100-18B-50UG), or 50 µM PDGF (ThermoFisher: 100-14B-50UG) in EBM-2 containing 1% BSA were introduced by sequentially adding 15 µL to the inlet port while aspirating 15 µL from the opposite port, repeated once to achieve 30 µL in the reservoir. For EA.hy926 cells, EBM-2 was substituted with DMEM.

For growth factor-mediated migration assays within collagen IV or collagen VI coated chambers, the µ-Slide ibidi chamber was pre-coated with 100 µg/mL collagen IV or VI overnight at 4°C, followed by three washes with EBM-2 containing 1% BSA (for HMVEC-d) or DMEM with 1% BSA (for EA.hy926 cells). Cells were then seeded in their respective media (HMVEC-d in EBM-2/1% BSA or EA.hy926 in DMEM/1% BSA) and allowed to adhere for 2 hours at 37°C in a 5% CO₂ incubator. Subsequent steps were performed identically to the migration assay protocol as described in the previous paragraph.

To do live cell imaging we did a time-lapse microscopy of migrating HMVEC-d using an inverted brightfield light microscope, capturing images every 10 minutes to a total of 80 images. Then, using the “Manual Tracking” option in ImageJ (NIH), we performed individual cell tracking of 45-50 cells in each experiment. We used the “Chemotaxis and Migration Tool V2.0” (ibidi GmbH, Gräfelfing, Germany) to analyze cell tracks, generate and quantify chemotaxis and forward migration index (FMI), and perform statistical tests.

### Stable Cell Line Generation via Lentiviral Transduction

Lenti-X 293T cells were grown in a 6 cm plate to 50-70% confluency in 5 mL of media. Lentiviral particles were produced by transfecting Lenti-293T cells with a transfer plasmid containing the gene of interest, psPAX2 (a gift from Didier Trono, Addgene plasmid # 12260), and pCMV-VSV-G (a gift from Bob Weinberg, Addgene plasmid # 8454) using TransIT-Lenti reagent (Muris Bio:6604, Madison, WI, USA). Specifically, 3 μL of TransIT-Lenti reagent was mixed with 200 μL of FBS-free media, combined with 1 μg of the transfer plasmid (pCW57.1), 4 μg of PAX2, and 1 μg of VSV-G in 200 μL of FBS-free media. The mixtures were incubated for 15 minutes at room temperature and then added dropwise to the Lenti-X 293T cells. A second 6 cm plate of Lenti-X293T cells received the same transfection protocol but with 9 μL of TransIT-Lenti reagent. The plates were incubated at 37°C for 48 hours before harvesting and pooling the virus-containing supernatant. The supernatant was mixed from two plates and centrifuged at 500 x g for 5 minutes, filtered through a 0.45 μm filter, and used immediately or stored at -80°C.

For transduction, EA.hy926 cells at 50-70% confluency in a 10 cm plate were treated with 5 mL of virus-containing media supplemented with 5 μL of Transduce-IT reagent and 5 mL of DMEM with 10% FBS (without penicillin-streptomycin). Cells were incubated for 48-72 hours before undergoing puromycin selection to eliminate non-transduced cells. EA.hy926 cells were treated with 10 ng/μL of puromycin for 24 hours, and cell survivability was monitored. Puromycin pressure was maintained for 72 hours with adjustments as needed.

Following selection, cells were treated with doxycycline (250ng/ml) to upregulate CMG2 expression, and sorted via FACS. We used PA conjugated to Alexa-Fluor-546 (ThermoFischer) [34] to measure CMG2 overexpression, comparing transduced cells to wild-type EA.hy926 cells treated with doxycycline and GFP-conjugated PA.

### MS sample preparation

CMG2-V5 was lentivirally-transduced into EA.hy926 cells, which were treated with 250ng/ml doxycycline for 48 hours to induce CMG2 expression. Then, we added 500ng/ml PA and immediately put the plate at 4°C followed by 10 minutes incubation at 37°C. Cells were washed 4x with ice-cold PBS and lysed with IP lysis buffer (Pierce, 87787) supplemented with 1X protease inhibitor cocktail (ThermoFischer, Cat. # 87786). Then, 500µg cell lysate protein was incubated with buffer-equilibrated V5 beads for 1 hour on a wheel at 4°C. After this binding step, V5 beads were washed three times in washing buffer (10 mM Tris/Cl pH 7.5, 150 mM NaCl, 0.05 % NP40 Substitute, 0.5 mM EDTA), separated with magnetic stands, resuspended in 5% SDS buffer dissolved in 100mM TEAB (ThermoFisher, Cat. # 90114), and heated for 5 minutes at 95°C. After cooling, V5 bead supernatants were prepared using S-trap (ProtiFi, Fairport, NY, USA Cat. # C02-mini-80) for mass spec analysis. Samples were reduced with 22mM DTT for 10 minutes at 95°C and alkylated with iodoacetamide (IAA) (Sigma-Aldrich, St. Louis, MO, USA, Cat. # I1149) for 1 hour at room temperature. Samples trapped on the column after washing/binding step were digested using trypsin (enzyme/protein∼1:10) overnight at 37°C, and peptide concentration determined using the Pierce quantitative colorimetric peptide assay kit (ThermoFischer, Cat. # 23275). At the end, 50ng of peptide was aliquoted out for timsTOF LC-MS analysis.

### LC-MS/MS data acquisition

For each technical replicate, three injections of 50 ng each were analyzed using a UHPLC-Nano system (ThermoFisher) interfaced via a home-etched 10 μm emitter with a timsTOF Ultra mass spectrometer (Bruker Daltonics, Billerica, MA, USA) equipped with a CaptiveSpray 2 ESI source. Peptides were pre-concentrated on a trap column (5 cm × 150 μm, C18, 3.5 μm), separated on an analytical column (15 cm × 75 μm, C18, 1.7 μm) and eluted using a linear active gradient from solvent A (0.1% formic acid in water) to solvent B (0.1% formic acid in acetonitrile) over 20 min at a flow rate of 0.3μL/min (total gradient duration of 40 minutes). Spray voltage was kept at 1500 V for peptide ionization and scan mass range set to 100-1700 m/z. Both ramp and accumulation time were set to 150 ms using a 100% duty cycle. Under DIA-PASEF mode, 15 MS/MS windows and 5 ramps were used to fragment precursors across mass and mobility ranges of 400-1000 m/z and 0.64-1.45 V-s cm2, respectively. The collision energy was set to linearly increase from 20 (0.6 V-s cm2) to 63 eV (1.6 V-s cm2). All mass spectrometry data have been deposited to the ProteomeXchange Consortium via the PRIDE partner repository with the data set identifier PXD number.

### MS Data analysis

dia-PASEF raw data was uploaded to DIA-NN v1.8.1 for proteomics data processing. Uniprot human proteome against DIA data is applied to identify peptides and protein. We set the protease specificity to trypsin with a maximum number of two missed cleavages and required a minimum peptide length of 7 amino acids. The mass tolerances for precursor and fragment ions were set to ‘Dynamic’ for both MS1 and MS2 level. False discovery rates were controlled by a target-decoy approach to ≤1% at precursor and protein levels. Processed data was further analyzed following the method described in [35]. T-test and fold change calculated from excel sheet were transferred to GraphPad Prism to make a volcano plot.

### Gene Ontology (GO) Analysis of Proteomics Data

Proteomics resulted in the identification of 5596 proteins for which a ratio between vehicle and PA-treated samples could be determine and 5105 for which a P-value could be calculated. Of these, 1746 were significantly different (P<0.05 after Bonferroni correction), 1718 whose interaction with CMG2 was decreased by PA treatment and 28 whose interaction was increased. These proteins were subjected to gene ontology (GO) analysis [15–17] on 2025-6-27 using release 2025-06-01 and version 10.5281/zenodo.15611813. Proteins whose interaction with CMG2 was increased by PA treatment were not significantly associated with any GO human biological processes, but those decreased were associated with many. Reasoning that those proteins whose interaction was most strongly changed by PA would give the best indication of the biological processes disrupted by PA treatment, we performed GO analysis on the most strongly affected subsets. We found no GO associations up to 100 proteins, but when we used the top 150 proteins, we identified 5 biological processes in the top tier of enrichment. Four of these were likely related to CMG2 production and transport to the cell membrane. But one, “actin cytoskeleton organization”, was interpretable as affecting endothelial cell chemotaxis and thereby angiogenesis. In the whole dataset of interactors significantly affected by PA treatment, 107 different proteins were associated with this GO term, which comprises 543 total. When we further extended the search dataset to the top 200 proteins, an additional 9 GO terms were identified, the most strongly enriched of which was “positive regulation of lamellipodium assembly”. In the whole dataset, 10 differentially interacting proteins were associated with this GO term, which comprises 28 total.

### Protein A/G and V5 magnetic beads Immunoprecipitation

To perform native actin co-immunoprecipitations (IP) from HMVEC-d or EA.hy926 cells, cells were washed three times in cold 1x PBS on ice and lysed in IP Buffer (ThermoFisher, Cat. # 87787) for 5 min on ice with periodic mixing. Lysates were then centrifuged at 4°C at 5,000 rpm (Eppendorf 5424R) for 10 minutes to pellet debris and the supernatant was then pre-cleared with protein A/G beads (ThermoFisher, Cat. # 80105G) for 1 h on a wheel at 4°C. Meanwhile, 3 μl of goat anti-human CMG2 (R&D Systems, Minneapolis, MN, USA, Cat. # AF2940) and 10 μl of protein A/G beads (ThermoFisher, Cat. # 80105G) were incubated on a rotating wheel at 4°C for one hour. Finally, the pre-cleared supernatant was incubated with protein A/G beads pre-incubated with goat anti-human CMG2 for 2 h on a rotating wheel at 4°C. Beads were collected with magnetic stands and washed three times in a wash buffer (PBS, containing 0.05% Tween-20). Lastly, the beads were boiled in Laemmli buffer (Bio-Rad) for 5 min before being subjected to western blotting.

For the anti-V5-bead immunoprecipitation, cell lysis was the same as with the protein A/G beads. Prior to protein binding, anti-V5-beads (ChromoTek, Planegg, Germany, Cat. # v5tma) were equilibrated in the dilution buffer (10 mM Tris/Cl pH 7.5, 150 mM NaCl, 0.5 mM EDTA) and collected with a magnet. Then, cell lysates were incubated with equilibrated beads for 1 hour on a wheel at 4°C, V5 beads were separated with magnetic stands and resuspended in wash buffer (10 mM Tris/Cl pH 7.5, 150 mM NaCl, 0.05 % NP40 Substitute, 0.5 mM EDTA) three times. Finally, anti-V5-beads were resuspended and boiled in the Laemmli buffer for 5 minutes at 95°C and supernatants were used for SDS-PAGE/western blotting analysis.

### Live-confocal imaging of migrating EA.hy926 cells with expression of fluorescently tagged protein

EA.hy926 cells transfected with the F-actin marker Lifeact-mCherry (a gift from Klaus Hahn, Addgene plasmid # 193300) and CMG2-mNeonGreen were generated. To actively track the migration, co-transfected EA.hy926 cells were trypsinized, plated on ibidi chemotaxis chambers (ibidi USA, Fitchburg, WI, Cat. # 80326), and incubated at 37°C, 5% CO2 for 2 hours to allow cell adherence. Adherent cells in the observation area were washed with FluoroBrite DMEM (ThermoFischer, Cat. # A1896701) with 1% BSA 4 times and then, 70uL of FluoroBrite DMEM alone were loaded into the left channel and FluoroBrite DMEM containing final 2.5% FBS was added to the right side of the chamber to create a chemo-attractant gradient. Migration was recorded using a Leica DMI8 live cell microscope using a 60x objective (Leica Microsystems, Buffalo Grove, IL, USA). Specifically, the ibidi device with a drop of immersion oil was placed on the plate heater at the setting of 37°C and supplied with 5% CO2 within a closed perfusion chamber. To minimize stage drift, the ibidi device was equilibrated for 30 minutes before beginning a long capture. Images were captured every 10 minutes with a 10-hour time-lapse duration.

### Immunocytochemistry, Confocal Microscopy, and Colocalization

At the end of a chemotaxis experiment, chemotaxing HMVEC-d were washed with 1X cold PBS then fixed with 4% paraformaldehyde (PFA in 1X PBS) for 12 minutes at room temperature (RT). Following three washes with 1X cold PBS, cells were permeabilized using 0.1% Triton X-100 with 1% BSA in PBS for 3 minutes, then cells were gently washed again three times with PBS. Non-specific binding was blocked with 5% BSA for 60 minutes at RT and HMVEC-d were stained with 1:1000 rabbit anti-CMG2 polyclonal antibody (Invitrogen, Catalog # PA5-118953) in 1X cold PBS containing 3% BSA and incubated for overnight at 4°C, then washed three times with 1X cold PBS. Secondary antibody / counterstaining cocktail was prepared by diluting DyLight 594 goat anti-rabbit IgG (Vector Lab, DI-1594-1.5) to stain CMG2) and phalloidin-AF488 (Life Technologies, Carlsbad, CA, USA, Cat. # A12379, to label F-actin) in 1:1000 ratio in 3% BSA containing PBS. Cells were co-stained with counter stain cocktail by incubating for 4 hours at 4°C in the dark. After three gentle washings with 1x cold PBS, cells were mounted with an anti-fade mounting media Fluoromount-G® (SouthernBiotech, Birmingham, AL, USA, Cat. #: 0100-01) and then wells were sealed with plugs to prevent drying. Then cells were imaged using a Zeiss LSM 880 Confocal Laser Scanning Microscope (Zeiss, Jena, Germany). We used a high magnification set up with a Plan-Apochromat 63×/1.4 NA oil immersion objective for imaging. To ensure the highest possible image quality, a drop of Zeiss immersion oil (refractive index ∼1.518) was applied to the lens before placing the slide. To quantify colocalization, we used ImageJ and performed Pearson correlation analysis with the Coloc2 function under the colocalization plugin (NIH) in Fiji (ImageJ).

### Western blot

HMVEC-d were cultured in EGM-2 complete medium overnight then treated with 4nM PA or 1ug PGG for 30 min on ice then incubated in a CO2 incubator for 3 hours and 24 hours respectively. Cells were washed twice with cold 1X PBS and lysed in RIPA lysis buffer containing protease inhibitors (PI, Roche cOmplete™ ULTRA Tablets, Sigma-Aldrich, Cat. # 5892970001) then incubated on ice for 30 minutes. Then, protein samples were centrifuged using an Eppendorf 5417R Centrifuge, Rotor FA-45-30-11 at 10,000 rpm for 10 minutes at 4°C and supernatant were collected and boiled in 1x Laemmli buffer (Laemmli SDS-Sample Buffer (6X, Reducing), Boston BioProducts, Milford, MA, USA, Cat. # BP-111R) for 5 min at 95°C. Total protein (20μg) was loaded on SDS-PAGE using 4-20% tris-glycine gradient gels (BioRad, 4–20% Mini-PROTEAN^®^ TGX™ Precast Protein Gels, Cat#4561094), then transferred onto a PVDF membrane (Thermo Scientific, 0.45 μm, Cat. # PI88518). Prior to antibody binding, the membrane was pre-blocked with 5% BSA in 1X TBS (Boston BioProducts, Cat. # BM-301). Then, the membrane was incubated overnight with 1:1000 anti-CMG2 Rabbit Polyclonal antibody (Invitrogen, Catalog # PA5-118953) in blocking buffer. The next day, membrane was washed with 1x TBS (four times, each 10 mins) and then further incubated in 1:4000 HRP-coupled anti-rabbit IgG secondary antibody (Cell Signaling Technology, #7074) in blocking buffer. Lastly, the immunoblot was washed four times for 10 minutes in 1x TBS and developed using the ECL (ThermoFisher, Cat. # 32106) according to the manufacturer’s protocol, imaged on a BioRad GelDoc system (BioRad, Hercules, CA, USA), and analysis was performed using ImageJ.

### GST Fusion Protein Expression and Purification

For *vitro* assays, we constructed a pET21-derived vector expressing CMG2-ABD fused to an N-terminal Avi tag and C-terminal GST tag (Avi-GST-CMG2^ABD^). To express Avi-GST-CMG2^ABD^ and Avi-GST (negative control), the plasmid was transformed into BL21(DE3) (ThermoFisher). Cells were grown in LB broth medium containing 100µg/mL of kanamycin at 37°C with 250rpm shaking. Then, D-biotin (GoldBio, St. Louis, MO, USA) at a final concentration of 50µg/mL was added to the cultures at the OD_600_ 0.3 with continuous shaking at 250rpm. When the OD_600_ reached 0.6, Avi-GST-CMG2^ABD^ expression was initiated by adding 0.5mM IPTG and cultures were incubated at room temperature (∼25°C) for 6 hours with 250rpm shaking. Bacteria were harvested by centrifugation at 8000rpm in a Thermo, F9-6Χ1000 LEX rotor (ThermoFisher).

Cells were suspended in 20mM Tris-HCL, 1mM PMSF, 300µg/mL lysozyme, 10µg/mL DNAase, passed through an 18-gauge needle followed by a 24-gauge needle and then lysed at 18,000 psi in 2 passes through a NanoDeBEE 45-2 homogenizer (Pion Inc., Billerica, MA, USA). Lysed cells were centrifuged at 40,000rpm (ThermoFisher, A23 6X100) and 4°C for 30 minutes and supernatants were collected flowed across a GST trap FF column (Cytiva, Marlborough, MA, USA) to purify Avi-GST-CMG2^ABD^ or Av-GST protein. Protein was eluted with reduced glutathione (50mM Tri-HCl, pH8.0 with 10mM reduced glutathione) and then desalted in a Zeba spinning column equilibrated with 20mM Tri-HCl, 150mM NaCl and 1mM PMSF. We then added glycerol to 50% and stored our final purified protein at -20°C.

### BLI (Biolayer Interferometry) Assay

Biolayer interferometry was used to measure the binding affinity between CMG2 and F-actin, as reported by Wallner et [36]. The assay buffer was composed of 50 mM HEPES, 150 mM NaCl at pH 7.2, 0.1% Tween-20, 1 mg/mL BSA, and 0.02% NaN3. First, we incubated streptavidin biosensors (ForteBio, Menlo Park, CA, USA) in assay buffer for 30 minutes at room temperature; four sensors were loaded and run in parallel (3 sensors for actin binding, one for reference controls). Binding assays were performed by using an Octet RED96 biolayer interferometer (ForteBio) and the Octet 8.2 Data Acquisition software (ForteBio). Assays were performed at 30 °C and 1000 rpm shaking. The streptavidin sensors were equilibrated in assay buffer (120s) followed by loading of Avi-GST-CMG2^ABD^ (200s), baseline washing (500s), an association step with a serial dilution (500nM, 250nM and 125nM) of the actin (600s) and dissociation in assay buffer (1800 s). Processed data was fit to a 1:1 binding model to calculate kinetic and thermodynamic parameters. Residuals were examined to assess the quality of fit, and no systematic deviation was observed.

### F-actin high speed centrifuge sedimentation Assay

We incubated 100µL of 2µM of polymerized F-actin (Cytoskeleton Inc., Denver, CO, USA) with 10µM of Avi-GST-CMG2^ABD^ or 10µM of Avi-GST in KMEI (50mM KCl, 1mM MgCl_2_, 1mM EGTA, 10mM imidazole and 1mM ATP. pH 7) buffer for 30 minutes at room temperature. Then, samples were centrifuged at 14.000g for 20 minutes at 4°C. After centrifugation, supernatants were gently transferred to a new tube and pellets from each sample were resuspended in the equal amount of 1× Laemmli buffer (BioRad, Cat. # 1610747). Denatured supernatants and pellets were analyzed with SDS-PAGE in a 4–20% gel (BioRad, Cat. # 4561094) followed by Coomassie staining (BioRad, Cat. # 1610436) to visualize proteins. At the end, a Li-Cor Odyssey CLX image scanner (Li-Cor Inc., Lincoln, NE, USA) was used to capture the stained gel image and measure protein band intensities.

### Scanning electron microscopy (SEM)

To prepare the sample for SEM, 1µM F-actin (Cytoskeleton Inc.) was pre-incubated with 7.5µM CMG2^ABD^-GST or 7.5µM Avi-GST in KMEI buffer for 30 minutes at room temperature. F-actin in-contrast imaging were performed as described by Schu et al [37]. In brief, a small drop of sample solution (∼5-10 microliters) was placed on a carbon stabilized, formvar coated, 200 mesh Cu TEM grid (FisherScientfic, Cat. # NC1721304) and incubated for approximately 1 minute. Sample solution was then wicked off of the grid using a filter paper wedge and a similar drop of 2% uranium acetate (aq) stain (Fisher Scientific, ThermoFischer, Cat. # NC1085517) was applied. After approximately 1 minute, the stain solution was wicked off of the grid with a filter paper wedge. The sample was then imaged in a Helios Nanolab 600 (ThermoFisher) using STEM mode.

### Statistical analysis

Statistical analysis employed a two-tailed, unpaired Student’s t-test or one-way ANOVA in GraphPad Prism 10.4.2, as described in figure legends. All data are presented as mean±SD. P<0.05 was considered to be statistically significant.

## Supporting information

Plasmid Maps

## Author Contribution

FJ, PCA, KAC, and MSR conceptualized the manuscript. FJ, RKM, JT, RO, EH, JF, EH, and ZJR performed investigation and FJ, RKM, JT, RO, EH, JF, EH, MSR, and KAC performed formal data analysis. MSR, PCA, and KAC were responsible for funding acquisition and MSR, KAC were responsible for project administration. The original draft was written by FJ, RKM, JT, EH, MSR, and KAC. All authors reviewed and/or edited the manuscript.

## Acknowledgements

We gratefully acknowledge the technical assistance and expertise provided by the Electron Microscopy Facility at Brigham Young University (BYU), with special thanks to Michael Standing for his invaluable support. We also extend our sincere appreciation for the generous funding provided by the BYU Simmons Center Fellowship.

## Data availability

The data that support the findings of this study are available from the corresponding author upon reasonable request.

---

**Fig S1:**
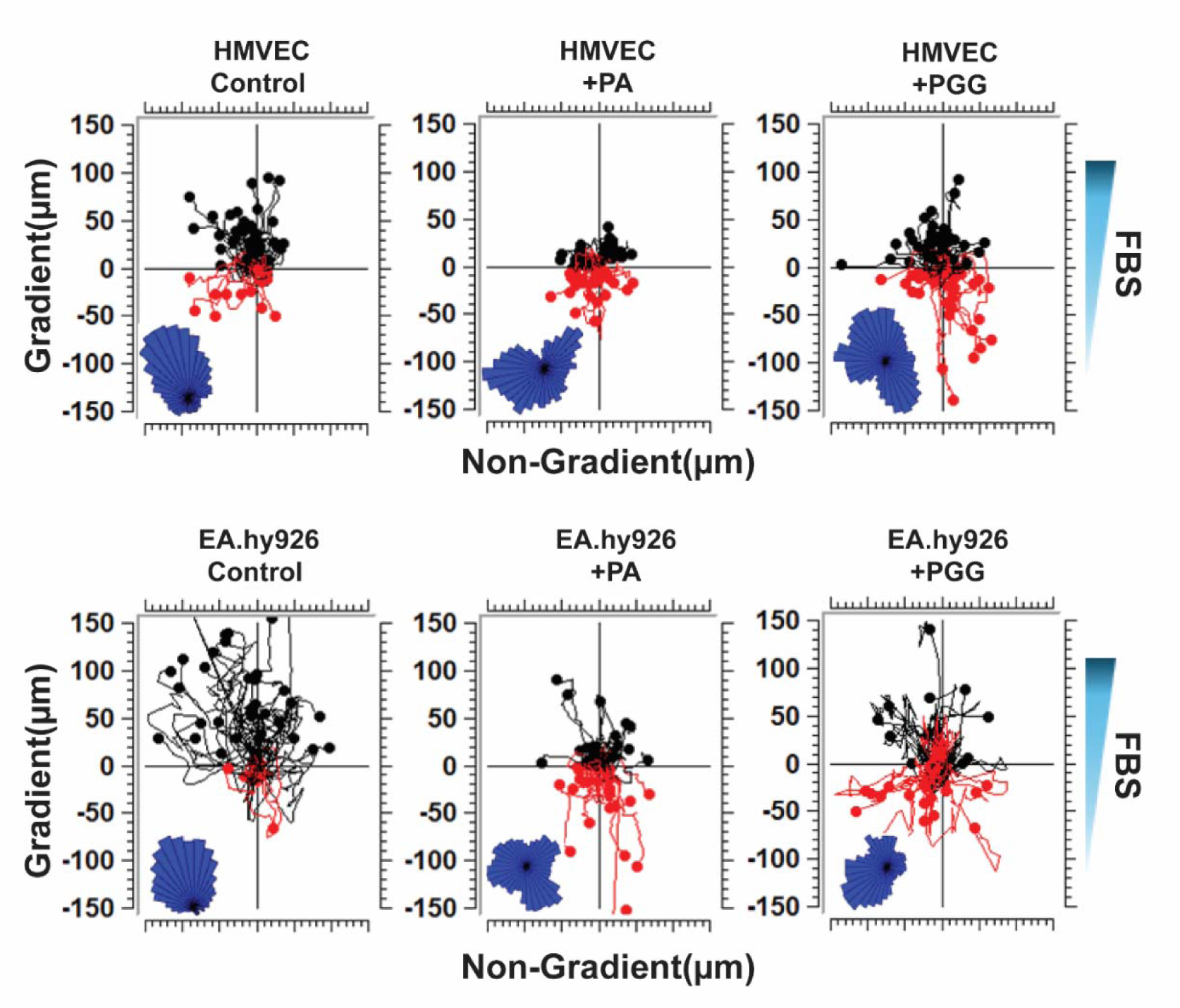
Sample chemotaxis tracks of HMVEC-d and EA.hy926 in the ibidi device. Each track plot represents the migration of HMVEC-d or EA.hy926 cells over the course of 8 hours toward serum in the ibidi microfluidic chip. The inset rose diagram depicts the sum of the vectored movement of the cells. Cells were seeded in the chip on serum, and treated with vehicle, PA, or PGG.

**Fig S2:**
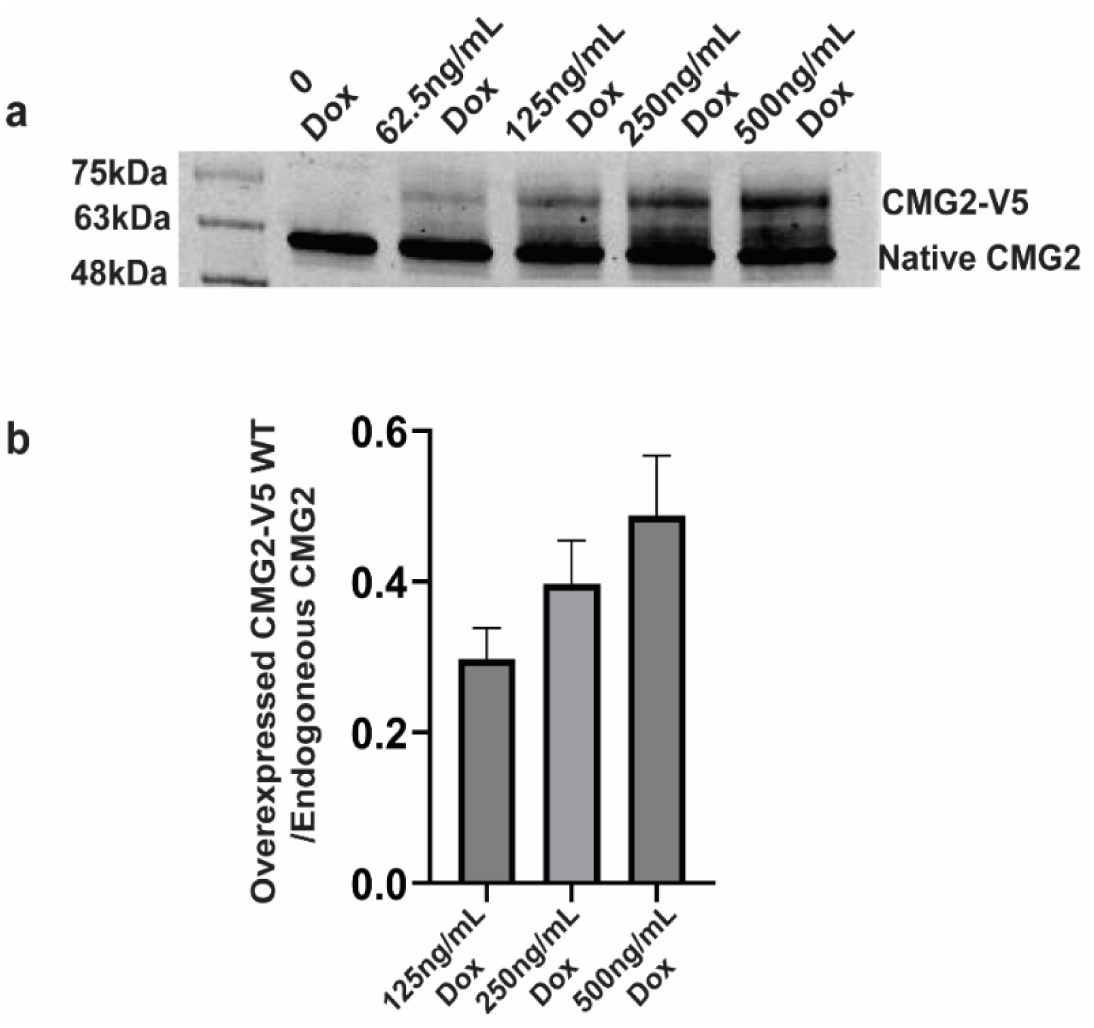
Validation of V5 tagged CMG2 lenti-transfection in EA. hy926 cells. **a:** Analysis of doxycycline-titration-controlled CMG2-V5 expression by western blot in EA.hy926 cells. **b:** The ratio of overexpressed CMG2-V5 to endogenous CMG2 quantified following treatment with 125ng/ml, 250ng/mL or 500ng/mL doxycycline (n=3 biological replicates).

**Fig S3:**
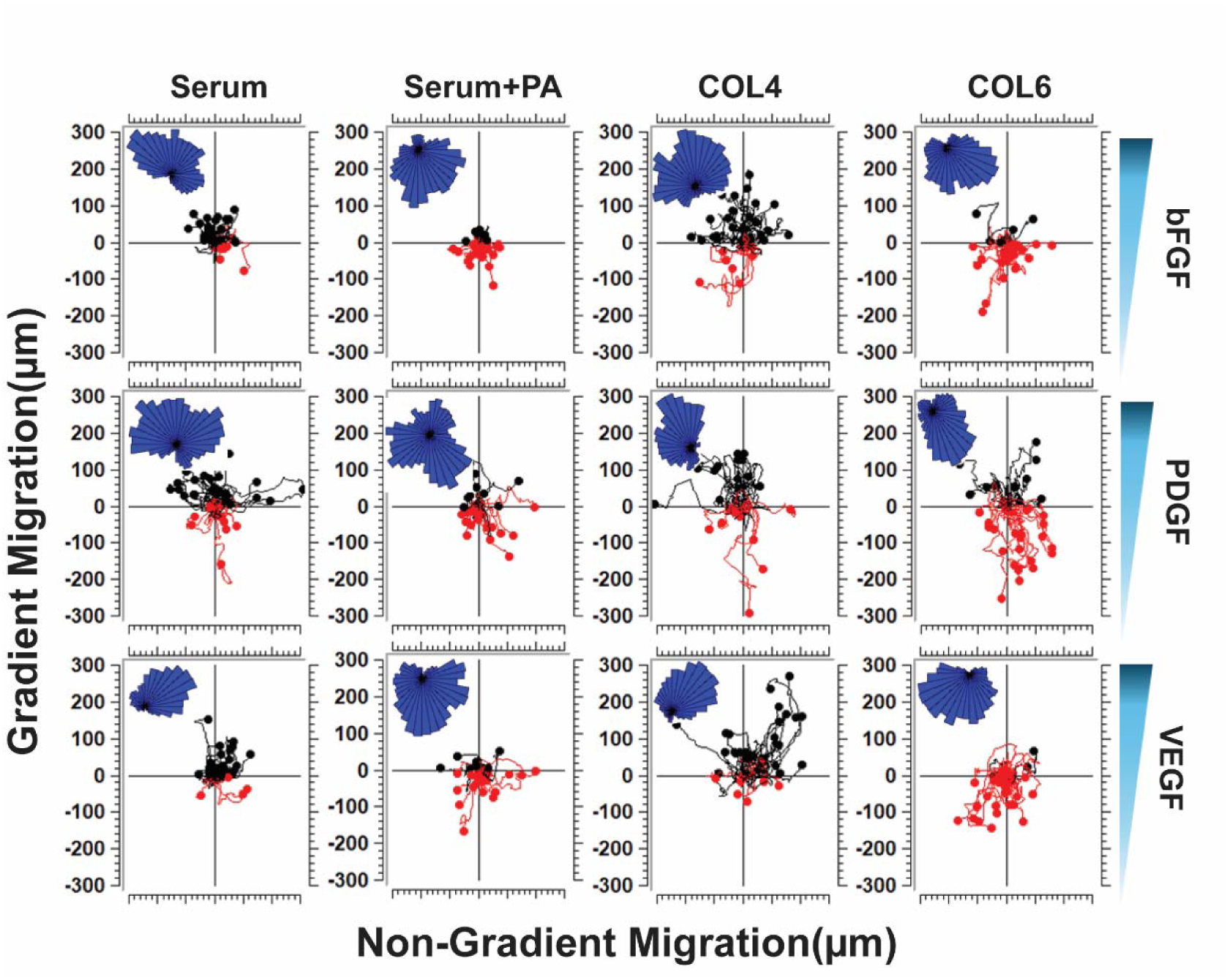
Sample chemotaxis tracks of HMVEC-d and EA.hy926 migrating in different chemotactic gradients on different matrices in the ibidi device. Each track plot represents the migration of endothelial cells (HMVEC-d) over the course of 8 hours towards its respective growth factor (bFGF, PDGF, and VEGF) in the ibidi microfluidic chip. The inset rose diagram depicts the sum of the vectored movement of the cells. Cells were seeded in the chip on either serum, Collagen IV, or Collagen VI.

**Fig S4:**
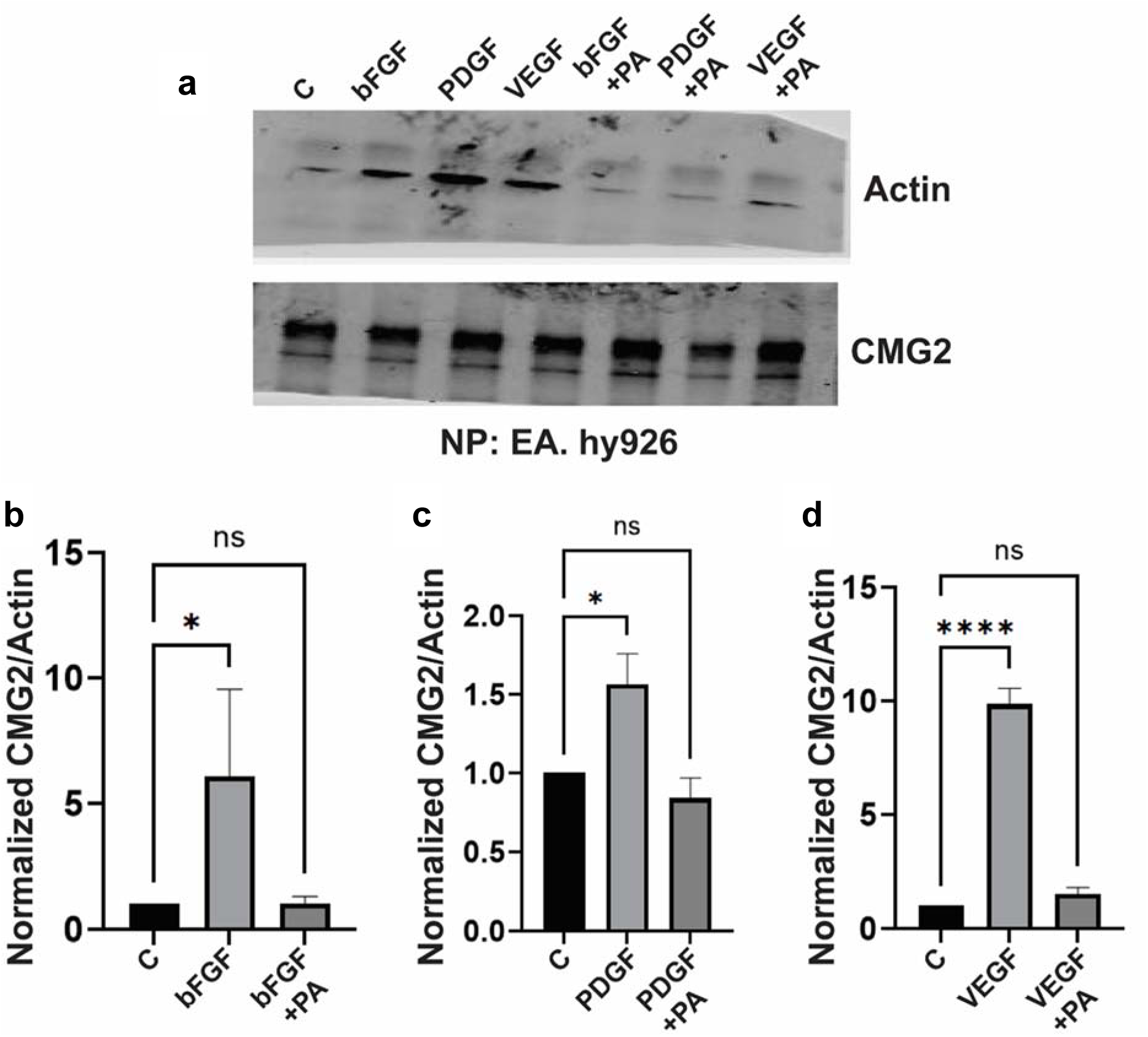
In EA.hy926 cells plated on serum, PA results in reduced growth-factor-induced actin-CMG2 interaction. **a:** Protein A/G beads bound to anti-CMG2 antibody were immunoprecipitated and analyzed by western blot using antibodies against actin and CMG2. **b-d:** Quantitative analysis of the CMG2/actin ratio among control, bFGF, bFGF+PA, PDGF, PDGF+PA, VEGF, VEGF+PA (3 biological replicates). Error bars are standard error of mean, *p<0.05; **p<0.01; ****p<0.0001 by one-way ANOVA in GraphPad Prism.

**Fig S5:**
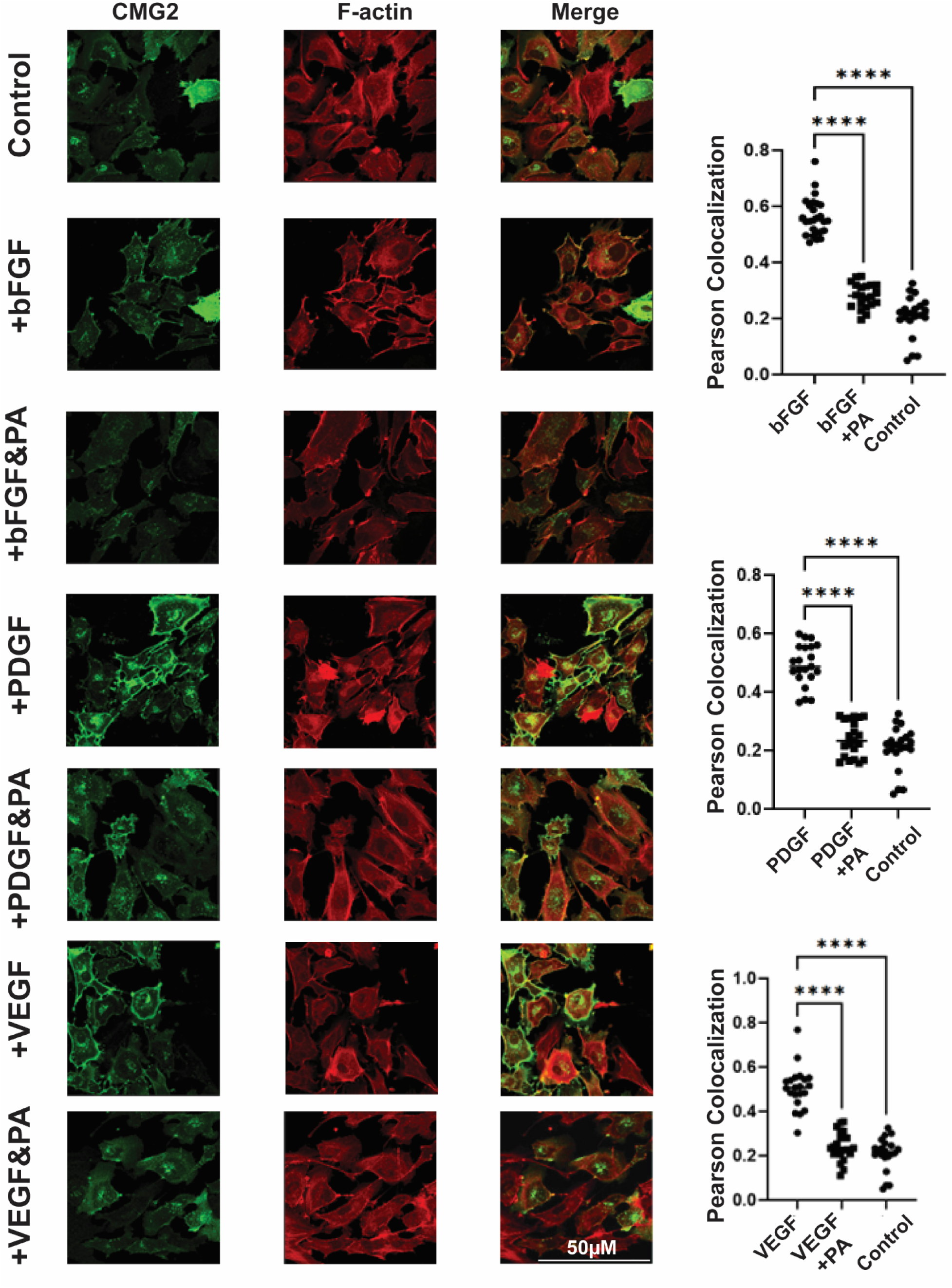
PA treatment decreases growth-factor-induced F-actin-CMG2 colocalization in the cell periphery. Immunofluorescence staining for CMG2-V5 and F-actin in CMG2-V5 transfected EA. hy926 cells. Medium alone is control. The addition of 50ng/mL bFGF, 50ng/mL PDGF or 50 ng/mL VEGF with or without 2nM PA was used for the different experiment settings. Peason’s colocalization between CMG2 and F-actin using Leica LAX-S software was calculated and loaded into the GraphPad Prism for ANOVA. (*p<0.05; **p<0.01; ****p<0.0001)

**Fig S6:**
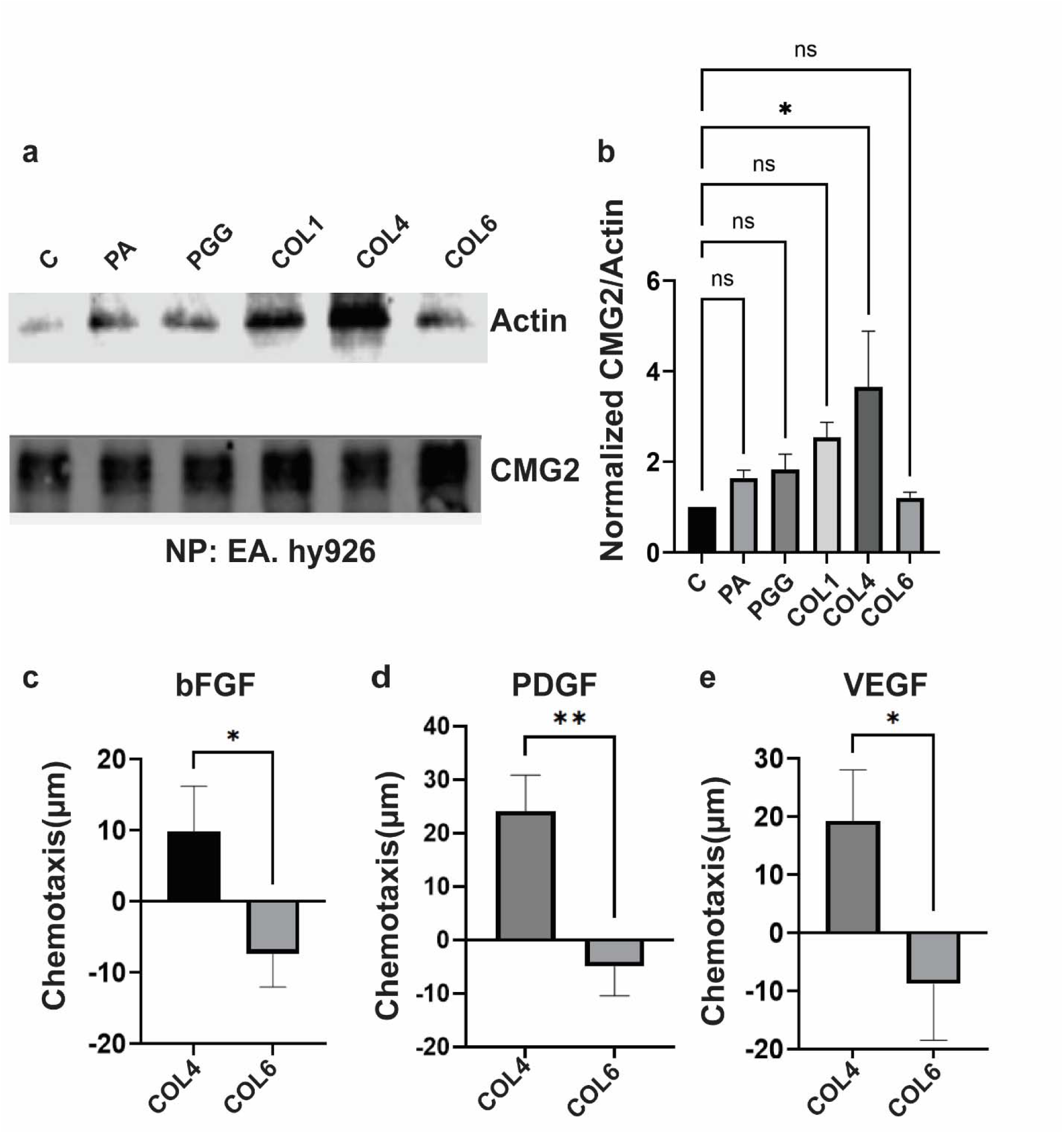
Matrix proteins affect actin interaction with CMG2 in EA. hy926 cells. **a:** Western blot after immunoprecipitation against actin and CMG2. **b:** Blot quantitation of normalized CMG2/actin from (a). **c-d**: EA. hy926 migration toward bFGF, PDGF or VEGF after plating on COL4 or COL6. Differences were assessed by one-way ANOVA in GraphPad Prism (*p<0.05; **p<0.01; ****p<0.0001).

**Fig S7:**
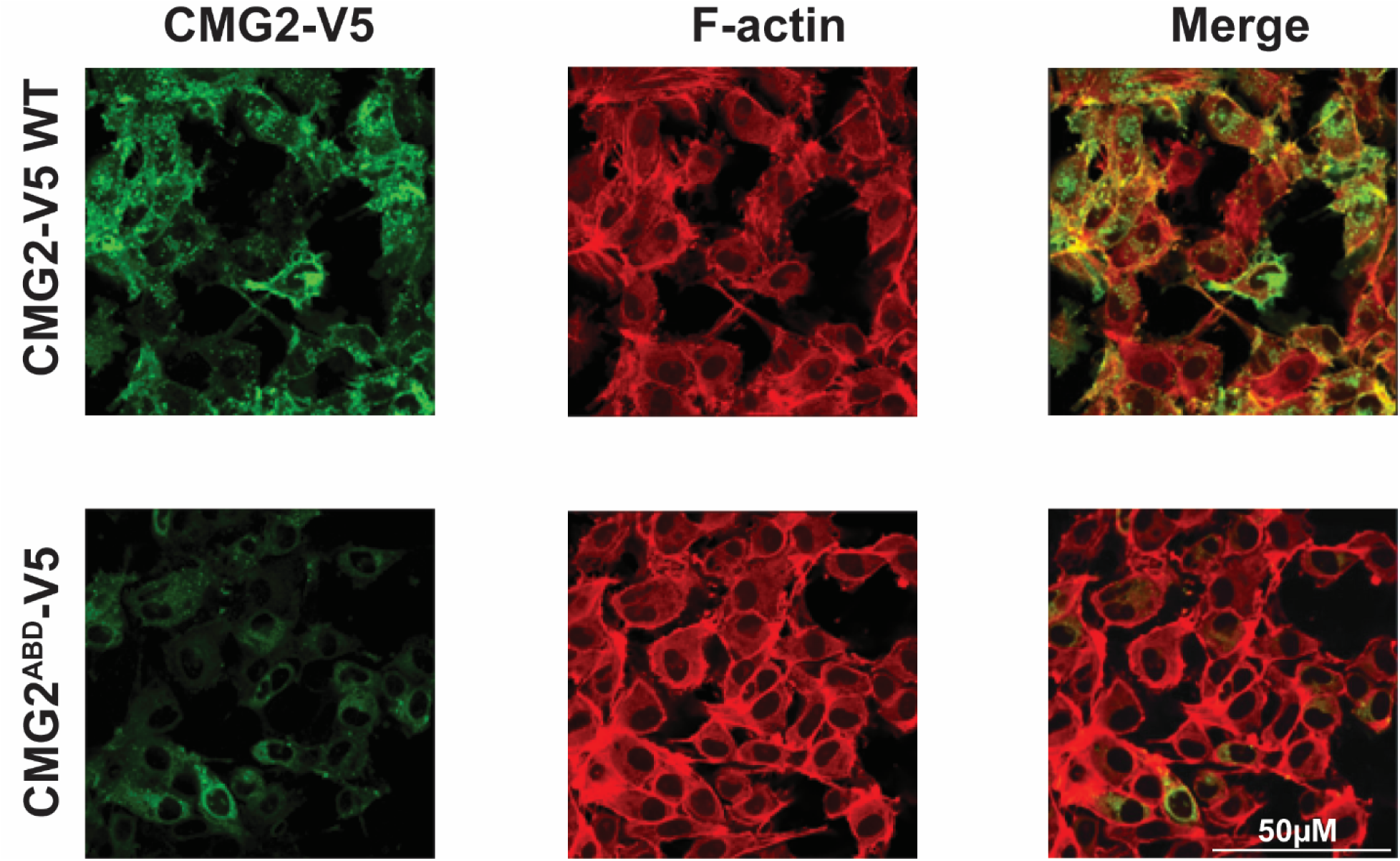
Reduced CMG2 and F-actin colocation in CMG2^ΔABD^. EA.hy926 cells were transduced with lentivirus expressing CMG2-WT and CMG2^ΔABD^, fixed, and immuno-stained with antibodies against V5 and F-actin. Colocalization was observed using 40X magnification in a Leica DMI8 confocal microscope.

