## Supplementary material for "CMG2 interaction with actin is required for growth factor-induced chemotaxis in endothelial cells": Plasmid Maps

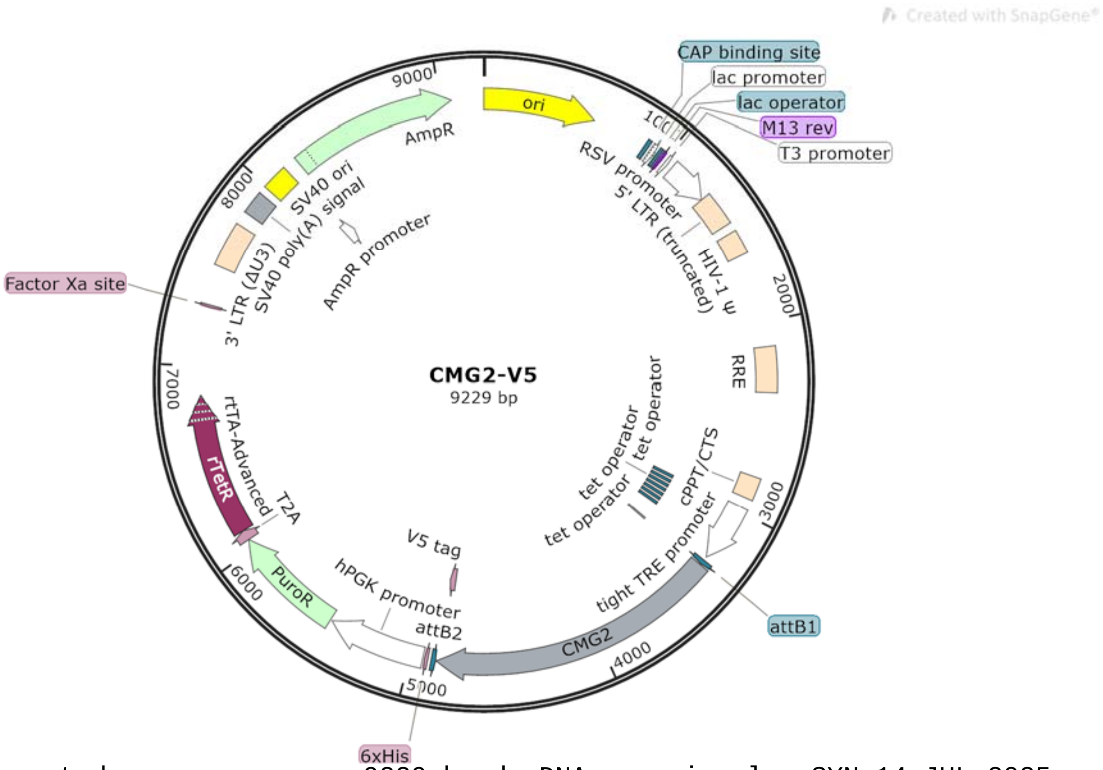


```
LOCUS       Exported                9229 bp ds-DNA     circular SYN 14-JUL-2025
DEFINITION  synthetic circular DNA
ACCESSION   .
VERSION     .
KEYWORDS    .
SOURCE      synthetic DNA construct
  ORGANISM  synthetic DNA construct
REFERENCE   1  (bases 1 to 9229)
  AUTHORS   Vascular Biology Program
  TITLE     Direct Submission
  JOURNAL   Exported Jul 14, 2025 from SnapGene 4.2.11
            http://www.snapgene.com
FEATURES             Location/Qualifiers
     source          1..9229
                     /organism="synthetic DNA construct"
                     /mol_type="other DNA"
     rep_origin      1..589
                     /direction=RIGHT
                     /label=ori
                     /note="high-copy-number ColE1/pMB1/pBR322/pUC origin of
                     replication"
     protein_bind    877..898
                     /label=CAP binding site
                     /bound_moiety="E. coli catabolite activator protein"
                     /note="CAP binding activates transcription in the presence
                     of cAMP."
     promoter        913..943
                     /label=lac promoter
                     /note="promoter for the E. coli lac operon"
     protein_bind    951..967
                     /label=lac operator
                     /bound_moiety="lac repressor encoded by lacI"
                     /note="The lac repressor binds to the lac operator to
                     inhibit transcription in E. coli. This inhibition can be
                     relieved by adding lactose or
                     isopropyl-beta-D-thiogalactopyranoside (IPTG)."
     primer_bind     975..991
                     /label=M13 rev
```

```
                /note="common sequencing primer, one of multiple similar
                variants"
promoter        1012..1030
                /label=T3 promoter
                /note="promoter for bacteriophage T3 RNA polymerase"
promoter        1058..1284
                /label=RSV promoter
                /note="Rous sarcoma virus enhancer/promoter"
LTR             1285..1465
                /label=5' LTR (truncated)
                /note="truncated 5' long terminal repeat (LTR) from HIV-1"
misc_feature    1512..1637
                /label=HIV-1 Psi
                /note="packaging signal of human immunodeficiency virus
                type 1"
misc_feature    2130..2363
                /label=RRE
                /note="The Rev response element (RRE) of HIV-1 allows for
                Rev-dependent mRNA export from the nucleus to the
                cytoplasm."
misc_feature    2805..2922
                /label=cPPT/CTS
                /note="central polypurine tract and central termination
                sequence of HIV-1"
promoter        2976..3290
                /label=tight TRE promoter
                /note="Tet-responsive promoter PTight, consisting of seven
                tet operator sequences followed by the minimal CMV
                promoter"
protein_bind    2984..3002
                /gene="tetO"
                /label=tet operator
                /bound_moiety="tetracycline repressor TetR"
                /note="bacterial operator O2 for the tetR and tetA genes"
protein_bind    3020..3038
                /gene="tetO"
                /label=tet operator
                /bound_moiety="tetracycline repressor TetR"
                /note="bacterial operator O2 for the tetR and tetA genes"
protein_bind    3055..3073
                /gene="tetO"
                /label=tet operator
                /bound_moiety="tetracycline repressor TetR"
                /note="bacterial operator O2 for the tetR and tetA genes"
protein_bind    3091..3109
                /gene="tetO"
                /label=tet operator
                /bound_moiety="tetracycline repressor TetR"
                /note="bacterial operator O2 for the tetR and tetA genes"
protein_bind    3127..3145
                /gene="tetO"
                /label=tet operator
                /bound_moiety="tetracycline repressor TetR"
                /note="bacterial operator O2 for the tetR and tetA genes"
protein_bind    3162..3180
                /gene="tetO"
                /label=tet operator
                /bound_moiety="tetracycline repressor TetR"
                /note="bacterial operator O2 for the tetR and tetA genes"
protein_bind    3198..3216
                /gene="tetO"
                /label=tet operator
                /bound_moiety="tetracycline repressor TetR"
```

```
                     /note="bacterial operator O2 for the tetR and tetA genes"
     protein_bind    3311..3335
                     /gene="mutant version of attB"
                     /label=attB1
                     /bound_moiety="BP Clonase(TM)"
                     /note="recombination site for the Gateway(R) BP reaction"
     regulatory      3339..3348
                     /regulatory_class="other"
                     /note="vertebrate consensus sequence for strong initiation
                     of translation (Kozak, 1987)"
     CDS             3341..4864
                     /codon_start=1
                     /label=CMG2
                     /translation="MVAERSPARSPGSWLFPGLWLLVLSGPGGLLRAQEQPSCRRAFDL
                     YFVLDKSGSVANNWIEIYNFVQQLAERFVSPEMRLSFIVFSSQATIILPLTGDRGKISK
                     GLEDLKRVSPVGETYIHEGLKLANEQIQKAGGLKTSSIIIALTDGKLDGLVPSYAEKEA
                     KISRSLGASVYCVGVLDFEQAQLERIADSKEQVFPVKGGFQALKGIINSILAQSCTEIL
                     ELQPSSVCVGEEFQIVLSGRGFMLGSRNGSVLCTYTVNETYTTSVKPVSVQLNSMLCPA
                     PILNKAGETLDVSVSFNGGKSVISGSLIVTATECSNGIAAIIVILVLLLLLGIGLMWWF
                     WPLCCKVVIKDPPPPPAPAPKEEEEEPLPTKKWPTVDASYYGGRGVGGIKRMEVRWGDK
                     GSTEEGARLEKAKNAVVKIPEETEEPIRPRPPRPKPTHQPPQTKWYTPIKGRLDALWAL
                     LRRQYDRVSLMRPQEGDEGRCINFSRVPSQGGGGSGKPIPNPLLGLDST"
     CDS             4820..4861
                     /codon_start=1
                     /product="epitope tag from simian virus 5"
                     /label=V5 tag
                     /translation="GKPIPNPLLGLDST"
     protein_bind    complement(4866..4890)
                     /gene="mutant version of attB"
                     /label=attB2
                     /bound_moiety="BP Clonase(TM)"
                     /note="recombination site for the Gateway(R) BP reaction"
     CDS             4908..4925
                     /codon_start=1
                     /product="6xHis affinity tag"
                     /label=6xHis
                     /translation="HHHHHH"
     promoter        4935..5445
                     /label=hPGK promoter
                     /note="human phosphoglycerate kinase 1 promoter"
     CDS             5455..6051
                     /codon_start=1
                     /gene="pac from Streptomyces alboniger"
                     /product="puromycin N-acetyltransferase"
                     /label=PuroR
                     /note="confers resistance to puromycin"
                     /translation="MTEYKPTVRLATRDDVPRAVRTLAAAFADYPATRHTVDPDRHIER
                     VTELQELFLTRVGLDIGKVWVADDGAAVAVWTTPESVEAGAVFAEIGPRMAELSGSRLA
                     AQQQMEGLLAPHRPKEPAWFLATVGVSPDHQGKGLGSAVVLPGVEAAERAGVPAFLETS
                     APRNLPFYERLGFTVTADVEVPEGPRTWCMTRKPGA"
     CDS             6052..6105
                     /codon_start=1
                     /product="2A peptide from Thosea asigna virus capsid
                     protein"
                     /label=T2A
                     /note="Eukaryotic ribosomes fail to insert a peptide bond
                     between the Gly and Pro residues, yielding separate
                     polypeptides."
                     /translation="EGRGSLLTCGDVEENPGP"
     CDS             6106..6852
                     /codon_start=1
                     /product="improved tetracycline-controlled transactivator"
                     /label=rtTA-Advanced
```

```
                     /note="In the Tet-On(R) system, rtTA-Advanced binds to the
                     Tet-responsive element and stimulates transcription only in
                     the presence of tetracycline or doxycycline."
                     /translation="MSRLDKSKVINGALELLNGVGIEGLTTRKLAQKLGVEQPTLYWHV
                     KNKRALLDALPIEMLDRHHTHFCPLEGESWQDFLRNNAKSYRCALLSHRDGAKVHLGTR
                     PTEKQYETLENQLAFLCQQGFSLENALYALSAVGHFTLGCVLEEQEHQVAKEERETPTT
                     DSMPPLLRQAIELFDRQGAEPAFLFGLELIICGLEKQLKCESGGPTDALDDFDLDMLPA
                     DALDDFDLDMLPADALDDFDLDMLPG"
     CDS             complement(7310..7321)
                     /codon_start=1
                     /product="Factor Xa recognition and cleavage site"
                     /label=Factor Xa site
                     /translation="IEGR"
     LTR             7526..7759
                     /label=3' LTR (Delta-U3)
                     /note="self-inactivating 3' long terminal repeat (LTR) from
                     HIV-1"
     polyA_signal    7831..7952
                     /label=SV40 poly(A) signal
                     /note="SV40 polyadenylation signal"
     rep_origin      7992..8127
                     /label=SV40 ori
                     /note="SV40 origin of replication"
     promoter        8127..8198
                     /gene="bla"
                     /label=AmpR promoter
     CDS             8199..9059
                     /codon_start=1
                     /gene="bla"
                     /product="beta-lactamase"
                     /label=AmpR
                     /note="confers resistance to ampicillin, carbenicillin, and
                     related antibiotics"
                     /translation="MSIQHFRVALIPFFAAFCLPVFAHPETLVKVKDAEDQLGARVGYI
                     ELDLNSGKILESFRPEERFPMMSTFKVLLCGAVLSRIDAGQEQLGRRIHYSQNDLVEYS
                     PVTEKHLTDGMTVRELCSAAITMSDNTAANLLLTTIGGPKELTAFLHNMGDHVTRLDRW
                     EPELNEAIPNDERDTTMPVAMATTLRKLLTGELLTLASRQQLIDWMEADKVAGPLLRSA
                     LPAGWFIADKSGAGERGSRGIIAALGPDGKPSRIVVIYTTGSQATMDERNRQIAEIGAS
                     LIKHW"
ORIGIN
        1 ttgagatcct ttttttctgc gcgtaatctg ctgcttgcaa acaaaaaaac caccgctacc
       61 agcggtggtt tgtttgccgg atcaagagct accaactctt tttccgaagg taactggctt
      121 cagcagagcg cagataccaa atactgttct tctagtgtag ccgtagttag gccaccactt
      181 caagaactct gtagcaccgc ctacatacct cgctctgcta atcctgttac cagtggctgc
      241 tgccagtggc gataagtcgt gtcttaccgg gttggactca agacgatagt taccggataa
      301 ggcgcagcgg tcgggctgaa cggggggttc gtgcacacag cccagcttgg agcgaacgac
      361 ctacaccgaa ctgagatacc tacagcgtga gctatgagaa agcgccacgc ttcccgaagg
      421 gagaaaggcg gacaggtatc cggtaagcgg cagggtcgga acaggagagc gcacgaggga
      481 gcttccaggg ggaaacgcct ggtatcttta tagtcctgtc gggtttcgcc acctctgact
      541 tgagcgtcga tttttgtgat gctcgtcagg ggggcggagc ctatggaaaa acgccagcaa
      601 cgcggccttt ttacggttcc tggccttttg ctggcctttt gctcacatgt tctttcctgc
      661 gttatcccct gattctgtgg ataaccgtat taccgccttt gagtgagctg ataccgctcg
      721 ccgcagccga acgaccgagc gcagcgagtc agtgagcgag gaagcggaag agcgcccaat
      781 acgcaaaccg cctctccccg cgcgttggcc gattcattaa tgcagctggc acgacaggtt
      841 tcccgactgg aaagcgggca gtgagcgcaa cgcaattaat gtgagttagc tcactcatta
      901 ggcaccccag gctttacact ttatgcttcc ggctcgtatg ttgtgtggaa ttgtgagcgg
      961 ataacaattt cacacaggaa acagctatga ccatgattac gccaagcgcg caattaaccc
     1021 tcactaaagg gaacaaaagc tggagctgca agcttaatgt agtcttatgc aatactcttg
     1081 tagtcttgca acatggtaac gatgagttag caacatgcct tacaaggaga gaaaaagcac
     1141 cgtgcatgcc gattggtgga agtaaggtgg tacgatcgtg ccttattagg aaggcaacag
     1201 acgggtctga catggattgg acgaaccact gaattgccgc attgcagaga tattgtattt
     1261 aagtgcctag ctcgatacat aaacgggtct ctctggttag accagatctg agcctgggag
     1321 ctctctggct aactagggaa cccactgctt aagcctcaat aaagcttgcc ttgagtgctt
```

1381 caagtagtgt gtgcccgtct gttgtgtgac tctggtaact agagatccct cagacccttt
1441 tagtcagtgt ggaaaatctc tagcagtggc gcccgaacag ggacttgaaa gcgaaaggga
1501 aaccagagga gctctctcga cgcaggactc ggcttgctga agcgcgcacg gcaagaggcg
1561 aggggcggcg actggtgagt acgccaaaaa ttttgactag cggaggctag aaggagagag
1621 atgggtgcga gagcgtcagt attaagcggg ggagaattag atcgcgatgg gaaaaaattc
1681 ggttaaggcc agggggaaag aaaaaatata aattaaaaca tatagtatgg gcaagcaggg
1741 agctagaacg attcgcagtt aatcctggcc tgttagaaac atcagaaggc tgtagacaaa
1801 tactgggaca gctacaacca tcccttcaga caggatcaga agaacttaga tcattatata
1861 atacagtagc aaccctctat tgtgtgcatc aaaggataga gataaaagac accaaggaag
1921 ctttagacaa gatagaggaa gagcaaaaca aaagtaagac caccgcacag caagcggccg
1981 ctgatcttca gacctggagg aggagatatg agggacaatt ggagaagtga attatataaa
2041 tataaagtag taaaaattga accattagga gtagcaccca ccaaggcaaa gagaagagtg
2101 gtgcagagag aaaaaagagc agtgggaata ggagctttgt tccttgggtt cttgggagca
2161 gcaggaagca ctatgggcgc agcgtcaatg acgctgacgg tacaggccag acaattattg
2221 tctggtatag tgcagcagca gaacaatttg ctgagggcta ttgaggcgca acagcatctg
2281 ttgcaactca cagtctgggg catcaagcag ctccaggcaa gaatcctggc tgtggaaaga
2341 tacctaaagg atcaacagct cctggggatt tggggttgct ctggaaaact catttgcacc
2401 actgctgtgc cttggaatgc tagttggagt aataaatctc tggaacagat ttggaatcac
2461 acgacctgga tggagtggga cagagaaatt aacaattaca caagcttaat acactcctta
2521 attgaagaat cgcaaaacca gcaagaaaag aatgaacaag aattattgga attagataaa
2581 tgggcaagtt tgtggaattg gtttaacata acaaattggc tgtggtatat aaaattattc
2641 ataatgatag taggaggctt ggtaggttta agaatagttt ttgctgtact ttctatagtg
2701 aatagagtta ggcagggata ttcaccatta tcgtttcaga cccacctccc aaccccgagg
2761 ggacaattct cgacctcgag acaaatggca gtattcatcc acaattttaa aagaaaaggg
2821 gggattgggg ggtacagtgc aggggaaaga atagtagaca taatagcaac agacatacaa
2881 actaaagaat tacaaaaaca aattacaaaa attcaaaatt ttcgggttta ttacagggac
2941 agcagagatc cactttggcc gcgaatcgat atgtcgagtt tactccctat cagtgataga
3001 gaacgtatgt cgagtttact ccctatcagt gatagagaac gatgtcgagt ttactcccta
3061 tcagtgatag agaacgtatg tcgagtttac tccctatcag tgatagagaa cgtatgtcga
3121 gtttactccc tatcagtgat agagaacgta tgtcgagttt atccctatca gtgatagaga
3181 acgtatgtcg agtttactcc ctatcagtga tagagaacgt atgtcgaggt aggcgtgtac
3241 ggtgggaggc ctatataagc agagctcgtt tagtgaaccg tcagatcgcc tggagaattg
3301 gctagcatca acaagtttgt acaaaaaagc aggctgcacc atggtggcgg aacggtcccc
3361 cgctcgcagt cccggtagct ggctttttcc cgggttatgg cttttggtgt tgtctgggcc
3421 ggggggactc ctccgggctc aggaacagcc ctcttgtaga cgagcatttg atctgtattt
3481 cgtgctggac aaatctggct ccgtcgctaa taattggatt gagatatata actttgtgca
3541 gcagctggct gaacgcttcg tttcaccgga aatgagactg tccttcatcg tgtttagcag
3601 ccaggctact attatcctcc ctctcactgg agaccgggga aagatctcca agggtctcga
3661 agacctcaag cgggtgtccc cggtgggaga aacatacatc catgagggcc tcaaattggc
3721 caacgaacag atacagaaag cgggggggcct gaaaacctcc tccataatca ttgccctcac
3781 agatggaaaa ctggatgggc tcgttccctc ctatgccgag aaagaggcaa aatcagtag
3841 atccctcggc gcctccgtct attgcgtggg ggtgctggac ttcgaacaag cgcagctgga
3901 gcgcattgcc gacagcaagg aacaagtgtt ccctgtgaaa ggaggatttc aagctctcaa
3961 agggatcata aattccatct tggcgcagtc ttgtactgaa atccttgaat tacaaccatc
4021 ttccgtttgt gtcggtgagg agttccagat cgtgttgagt gggaggggct tcatgctggg
4081 gagccgaaat ggcagcgttc tctgcaccta taccgtgaac gaaacatata ctactagtgt
4141 aaaaccagtg agcgtgcagc tgaacagcat gctgtgtcct gcccccatcc tgaataaagc
4201 gggggagact ctcgatgtta gcgtcagttt taatgggggg aagagcgtga ttagtggaag
4261 tctcattgtt accgctacag aatgtagcaa cggtatagcc gctatcattg ttatcctggt
4321 tctcctgttg cttctgggga taggactcat gtggtggttc tggccgctgt gttgtaaggt
4381 ggtaattaag gacccacctc caccacccgc acccgctccg aaagaggagg aggaagaacc
4441 gctgcctact aagaaatggc ccaccgtgga cgcttcatac tatggcgggc gtggcgtggg
4501 cggcattaaa aggatggagg taaggtgggg agataaggga tcaactgaag agggtgcacg
4561 actggagaag gctaagaatg ctgtcgtaaa gattcctgaa gagacagagg agcctattag
4621 gcctagaccc cccagaccaa agcccacaca tcaaccgccc caaacaaagt ggtatacccc
4681 aattaagggt agactggatg cattatgggc cctgctgcgc cgccagtacg acagagtgtc
4741 tttaatgcgg ccacaggagg gcgatgaggg gagatgtata aatttttcac gagtcccatc
4801 tcagggtgga ggtggctccg gcaagccaat ccctaaccct ctgttgggac tggatagcac
4861 ataggaccca gctttcttgt acaaagtggt ttagtaatga accggtccac caccaccacc
4921 accactaagg atccggggtt ggggttgcgc cttttccaag gcagccctgg gtttgcgcag
4981 ggacgcggct gctctgggcg tggttccggg aaacgcagcg gcgccgaccc tgggtctcgc
5041 acattcttca cgtccgttcg cagcgtcacc cggatcttcg ccgctaccct tgtgggcccc
5101 ccggcgacgc ttcctgctcc gcccctaagt cgggaaggtt ccttgcggtt cgcggcgtgc

5161 cggacgtgac aaacggaagc cgcacgtctc actagtaccc tcgcagacgg acagcgccag
5221 ggagcaatgg cagcgcgccg accgcgatgg gctgtggcca atagcggctg ctcagcaggg
5281 cgcgccgaga gcagcggccg ggaaggggcg gtgcgggagg cggggtgtgg ggcggtagtg
5341 tgggccctgt tcctgcccgc gcggtgttcc gcattctgca agcctccgga gcgcacgtcg
5401 gcagtcggct ccctcgttga ccgaatcacc gacctctctc cccagcaatt caccatgacc
5461 gagtacaagc ccacggtgcg cctcgccacc cgcgacgacg tccccagggc cgtacgcacc
5521 ctcgccgccg cgttcgccga ctaccccgcc acgcgccaca ccgtcgatcc ggaccgccac
5581 atcgagcggg tcaccgagct gcaagaactc ttcctcacgc gcgtcgggct cgacatcggc
5641 aaggtgtggg tcgcggacga cggcgccgcg gtggcggtct ggaccacgcc ggagagcgtc
5701 gaagcggggg cggtgttcgc cgagatcggc ccgcgcatgg ccgagttgag cggttcccgg
5761 ctggccgcgc agcaacagat ggaaggcctc ctggcgccgc accggcccaa ggagcccgcg
5821 tggttcctgg ccaccgtcgg cgtctcgccc gaccaccagg gcaagggtct gggcagcgcc
5881 gtcgtgctcc ccggagtgga ggcggccgag cgcgccgggg tgcccgcctt cctggagacc
5941 tccgcgcccc gcaacctccc cttctacgag cggctcggct tcaccgtcac cgccgacgtc
6001 gaggtgcccg aaggaccgcg cacctggtgc atgacccgca agcccggtgc cgaaggtaga
6061 ggttctctcc tcacttgtgg tgatgttgaa gaaaaccctg gtccaatgtc tagactggac
6121 aagagcaaag tcataaacgg agctctggaa ttactcaatg gtgtcggtat cgaaggcctg
6181 acgacaagga aactcgctca aaagctggga gttgagcagc ctaccctgta ctggcacgtg
6241 aagaacaagc gggccctgct cgatgccctg ccaatcgaga tgctggacag gcatcatacc
6301 cacttctgcc ccctggaagg cgagtcatgg caagactttc tgcggaacaa cgccaagtca
6361 taccgctgtg ctctcctctc acatcgcgac ggggctaaag tgcatctcgg cacccgccca
6421 acagagaaac agtacgaaac cctggaaaat cagctcgcgt tcctgtgtca gcaaggcttc
6481 tccctggaga acgcactgta cgctctgtcc gccgtgggcc actttacact gggctgcgta
6541 ttggaggaac aggagcatca agtagcaaaa gaggaaagag agacacctac caccgattct
6601 atgcccccac ttctgagaca agcaattgag ctgttcgacc ggcagggagc cgaacctgcc
6661 ttccttttcg gcctggaact aatcatatgt ggcctggaga aacagctaaa gtgcgaaagc
6721 ggcgggccga ccgacgccct tgacgatttt gacttagaca tgctcccagc cgatgccctt
6781 gacgactttg accttgatat gctgcctgct gacgctcttg acgattttga ccttgacatg
6841 ctccccgggt aactaagtaa ggatcgatcc aagatatcgt attcttaact atgttgctcc
6901 ttttacgcta tgtggatacg ctgctttaat gcctttgtat catgctattg cttcccgtat
6961 ggctttcatt ttctcctcct tgtataaatc ctggttgctg tctctttatg aggagttgtg
7021 gcccgttgtc aggcaacgtg gcgtggtgtg cactgtgttt gctgacgcaa cccccactgg
7081 ttggggcatt gccaccacct gtcagctcct ttccgggact ttcgctttcc ccctccctat
7141 tgccacggcg gaactcatcg ccgcctgcct tgcccgctgc tggacagggg ctcggctgtt
7201 gggcactgac aattccgtgg tgttgtcggg gaagctgacg tcctttccat ggctgctcgc
7261 ctgtgttgcc acctggattc tgcgcgggac gtccttctgc tacgtccctt cggccctcaa
7321 tccagcggac cttccttccc gcggcctgct gccggctctg cggcctcttc cgcgtcttcg
7381 ccttcgccct cagacgagtc ggatctccct ttgggccgcc tccccgcctg tttcgcctcg
7441 gcgtccggac tagaggtacc tttaagacca atgacttaca aggcagctgt agatcttagc
7501 cactttttaa aagaaaaggg gggactggaa gggctaattc actcccaacg aagacaagat
7561 ctgctttttg cttgtactgg gtctctctgg ttagaccaga tctgagcctg ggagctctct
7621 ggctaactag ggaacccact gcttaagcct caataaagct tgccttgagt gcttcaagta
7681 gtgtgtgccc gtctgttgtg tgactctggt aactagagat ccctcagacc cttttagtca
7741 gtgtggaaaa tctctagcag tagtagttca tgtcatctta ttattcagta tttataactt
7801 gcaaagaaat gaatatcaga gagtgagagg aacttgttta ttgcagctta taatggttac
7861 aaataaagca atagcatcac aaatttcaca aataaagcat ttttttcact gcattctagt
7921 tgtggtttgt ccaaactcat caatgtatct tatcatgtct ggctctagct atcccgcccc
7981 taactccgcc catcccgccc ctaactccgc ccagttccgc ccattctccg ccccatggct
8041 gactaatttt ttttatttat gcagaggccg aggccgcctc ggcctctgag ctattccaga
8101 agtagtgagg aggctttttt ggaggccttt caaatatgta tccgctcatg agacaataac
8161 cctgataaat gcttcaataa tattgaaaaa ggaagagtat gagtattcaa catttccgtg
8221 tcgcccttat tccctttttt gcggcatttt gccttcctgt ttttgctcac ccagaaacgc
8281 tggtgaaagt aaaagatgct gaagatcagt tgggtgcacg agtgggttac atcgaactgg
8341 atctcaacag cggtaagatc cttgagagtt ttcgccccga agaacgtttt ccaatgatga
8401 gcacttttaa agttctgcta tgtggcgcgg tattatcccg tattgacgcc gggcaagagc
8461 aactcggtcg ccgcatacac tattctcaga atgacttggt tgagtactca ccagtcacag
8521 aaaagcatct tacggatggc atgacagtaa gagaattatg cagtgctgcc ataaccatga
8581 gtgataacac tgcggccaac ttacttctga caacgatcgg aggaccgaag gagctaaccg
8641 cttttttgca caacatgggg gatcatgtaa ctcgccttga tcgttgggaa ccggagctga
8701 atgaagccat accaaacgac gagcgtgaca ccacgatgcc tgtagcaatg gcaacaacgt
8761 tgcgcaaact attaactggc gaactactta ctctagcttc ccggcaacaa ttaatagact
8821 ggatggaggc ggataaagtt gcaggaccac ttctgcgctc ggcccttccg gctggctggt
8881 ttattgctga taaatctgga gccggtgagc gtgggtctcg cggtatcatt gcagcactgg

```
8941 ggccagatgg taagccctcc cgtatcgtag ttatctacac gacggggagt caggcaacta
9001 tggatgaacg aaatagacag atcgctgaga taggtgcctc actgattaag cattggtaac
9061 tgtcagacca agtttactca tatatacttt agattgattt aaaacttcat ttttaattta
9121 aaaggatcta ggtgaagatc ctttttgata atctcatgac caaaatccct taacgtgagt
9181 tttcgttcca ctgagcgtca gaccccgtag aaaagatcaa aggatcttc
//
```

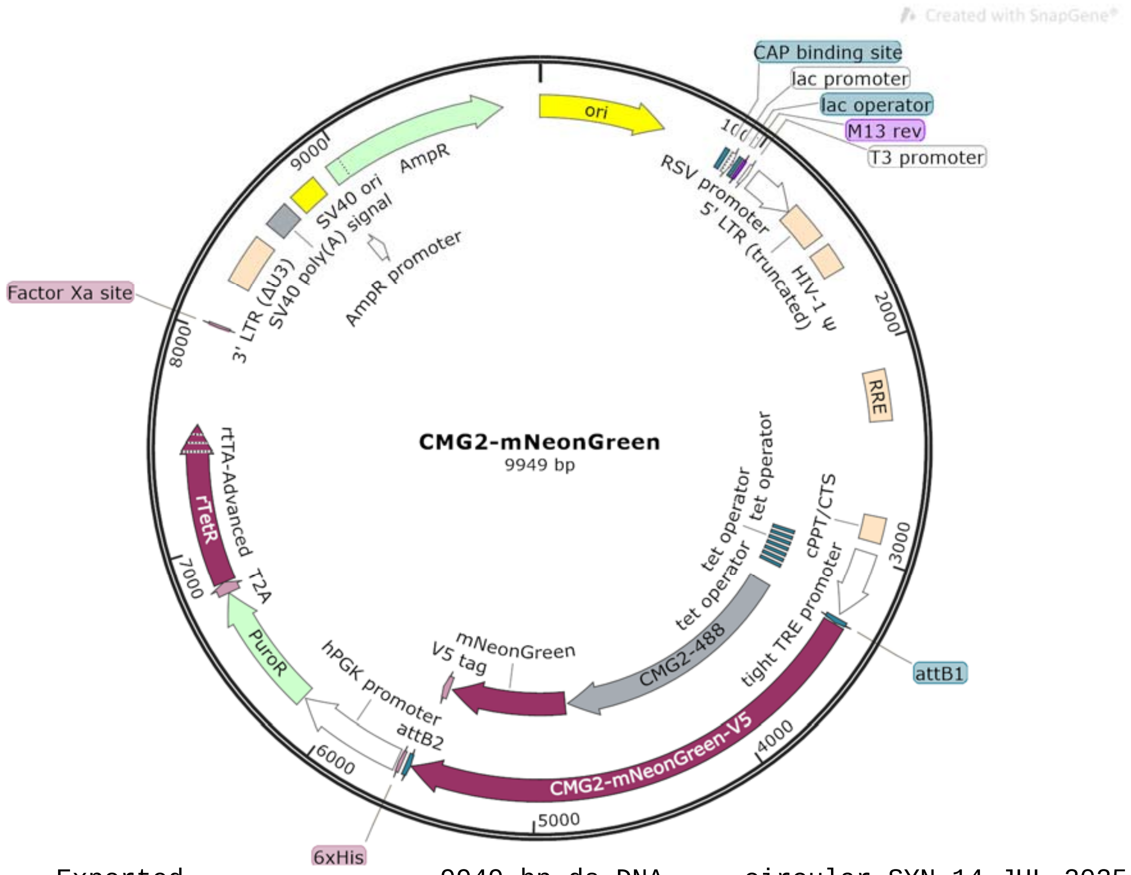


```
LOCUS       Exported                9949 bp ds-DNA     circular SYN 14-JUL-2025
DEFINITION  synthetic circular DNA
ACCESSION   .
VERSION     .
KEYWORDS    .
SOURCE      synthetic DNA construct
  ORGANISM  synthetic DNA construct
REFERENCE   1  (bases 1 to 9949)
  AUTHORS   Vascular Biology Program
  TITLE     Direct Submission
  JOURNAL   Exported Jul 14, 2025 from SnapGene 4.2.11
            http://www.snapgene.com
FEATURES             Location/Qualifiers
     source          1..9949
                     /organism="synthetic DNA construct"
                     /mol_type="other DNA"
     rep_origin      1..589
                     /direction=RIGHT
                     /label=ori
                     /note="high-copy-number ColE1/pMB1/pBR322/pUC origin of
                     replication"
     protein_bind    877..898
                     /label=CAP binding site
                     /bound_moiety="E. coli catabolite activator protein"
                     /note="CAP binding activates transcription in the presence
                     of cAMP."
     promoter        913..943
                     /label=lac promoter
                     /note="promoter for the E. coli lac operon"
     protein_bind    951..967
                     /label=lac operator
                     /bound_moiety="lac repressor encoded by lacI"
                     /note="The lac repressor binds to the lac operator to
                     inhibit transcription in E. coli. This inhibition can be
```

```
                relieved by adding lactose or
                isopropyl-beta-D-thiogalactopyranoside (IPTG)."
primer_bind     975..991
                /label=M13 rev
                /note="common sequencing primer, one of multiple similar
                variants"
promoter        1012..1030
                /label=T3 promoter
                /note="promoter for bacteriophage T3 RNA polymerase"
promoter        1058..1284
                /label=RSV promoter
                /note="Rous sarcoma virus enhancer/promoter"
LTR             1285..1465
                /label=5' LTR (truncated)
                /note="truncated 5' long terminal repeat (LTR) from HIV-1"
misc_feature    1512..1637
                /label=HIV-1 Psi
                /note="packaging signal of human immunodeficiency virus
                type 1"
misc_feature    2130..2363
                /label=RRE
                /note="The Rev response element (RRE) of HIV-1 allows for
                Rev-dependent mRNA export from the nucleus to the
                cytoplasm."
misc_feature    2805..2922
                /label=cPPT/CTS
                /note="central polypurine tract and central termination
                sequence of HIV-1"
promoter        2976..3290
                /label=tight TRE promoter
                /note="Tet-responsive promoter PTight, consisting of seven
                tet operator sequences followed by the minimal CMV
                promoter"
protein_bind    2984..3002
                /gene="tetO"
                /label=tet operator
                /bound_moiety="tetracycline repressor TetR"
                /note="bacterial operator O2 for the tetR and tetA genes"
protein_bind    3020..3038
                /gene="tetO"
                /label=tet operator
                /bound_moiety="tetracycline repressor TetR"
                /note="bacterial operator O2 for the tetR and tetA genes"
protein_bind    3055..3073
                /gene="tetO"
                /label=tet operator
                /bound_moiety="tetracycline repressor TetR"
                /note="bacterial operator O2 for the tetR and tetA genes"
protein_bind    3091..3109
                /gene="tetO"
                /label=tet operator
                /bound_moiety="tetracycline repressor TetR"
                /note="bacterial operator O2 for the tetR and tetA genes"
protein_bind    3127..3145
                /gene="tetO"
                /label=tet operator
                /bound_moiety="tetracycline repressor TetR"
                /note="bacterial operator O2 for the tetR and tetA genes"
protein_bind    3162..3180
                /gene="tetO"
                /label=tet operator
                /bound_moiety="tetracycline repressor TetR"
                /note="bacterial operator O2 for the tetR and tetA genes"
```

```
protein_bind    3198..3216
                /gene="tetO"
                /label=tet operator
                /bound_moiety="tetracycline repressor TetR"
                /note="bacterial operator O2 for the tetR and tetA genes"
protein_bind    3311..3335
                /gene="mutant version of attB"
                /label=attB1
                /bound_moiety="BP Clonase(TM)"
                /note="recombination site for the Gateway(R) BP reaction"
regulatory      3339..3348
                /regulatory_class="other"
                /note="vertebrate consensus sequence for strong initiation
                of translation (Kozak, 1987)"
CDS             3341..5584
                /codon_start=1
                /label=CMG2-mNeonGreen-V5
                /translation="MVAERSPARSPGSWLFPGLWLLVLSGPGGLLRAQEQPSCRRAFDL
                YFVLDKSGSVANNWIEIYNFVQQLAERFVSPEMRLSFIVFSSQATIILPLTGDRGKISK
                GLEDLKRVSPVGETYIHEGLKLANEQIQKAGGLKTSSIIIALTDGKLDGLVPSYAEKEA
                KISRSLGASVYCVGVLDFEQAQLERIADSKEQVFPVKGGFQALKGIINSILAQSCTEIL
                ELQPSSVCVGEEFQIVLSGRGFMLGSRNGSVLCTYTVNETYTTSVKPVSVQLNSMLCPA
                PILNKAGETLDVSVSFNGGKSVISGSLIVTATECSNGIAAIIVILVLLLLLGIGLMWWF
                WPLCCKVVIKDPPPPPAPAPKEEEEEPLPTKKWPTVDASYYGGRGVGGIKRMEVRWGDK
                GSTEEGARLEKAKNAVVKIPEETEEPIRPRPPRPKPTHQPPQTKWYTPIKGRLDALWAL
                LRRQYDRVSLMRPQEGDEGRCINFSRVPSQGGGGSVSKGEEDNMASLPATHELHIFGSI
                NGVDFDMVGQGTGNPNDGYEELNLKSTKGDLQFSPWILVPHIGYGFHQYLPYPDGMSPF
                QAAMVDGSGYQVHRTMQFEDGASLTVNYRYTYEGSHIKGEAQVKGTGFPADGPVMTNSL
                TAADWCRSKKTYPNDKTIISTFKWSYTTGNGKRYRSTARTTYTFAKPMAANYLKNQPMY
                VFRKTELKHSKTELNFKEWQKAFTDVMGMDELYKGGGGSGKPIPNPLLGLDST"
misc_feature    3341..4804
                /label=CMG2-488
CDS             4820..5524
                /codon_start=1
                /label=mNeonGreen
                /translation="VSKGEEDNMASLPATHELHIFGSINGVDFDMVGQGTGNPNDGYEE
                LNLKSTKGDLQFSPWILVPHIGYGFHQYLPYPDGMSPFQAAMVDGSGYQVHRTMQFEDG
                ASLTVNYRYTYEGSHIKGEAQVKGTGFPADGPVMTNSLTAADWCRSKKTYPNDKTIIST
                FKWSYTTGNGKRYRSTARTTYTFAKPMAANYLKNQPMYVFRKTELKHSKTELNFKEWQK
                AFTDVMGMDELYK"
regulatory      5072..5081
                /regulatory_class="other"
                /note="vertebrate consensus sequence for strong initiation
                of translation (Kozak, 1987)"
CDS             5540..5581
                /codon_start=1
                /product="epitope tag from simian virus 5"
                /label=V5 tag
                /translation="GKPIPNPLLGLDST"
protein_bind    complement(5586..5610)
                /gene="mutant version of attB"
                /label=attB2
                /bound_moiety="BP Clonase(TM)"
                /note="recombination site for the Gateway(R) BP reaction"
CDS             5628..5645
                /codon_start=1
                /product="6xHis affinity tag"
                /label=6xHis
                /translation="HHHHHH"
promoter        5655..6165
                /label=hPGK promoter
                /note="human phosphoglycerate kinase 1 promoter"
CDS             6175..6771
```

```
                     /codon_start=1
                     /gene="pac from Streptomyces alboniger"
                     /product="puromycin N-acetyltransferase"
                     /label=PuroR
                     /note="confers resistance to puromycin"
                     /translation="MTEYKPTVRLATRDDVPRAVRTLAAAFADYPATRHTVDPDRHIER
                     VTELQELFLTRVGLDIGKVWVADDGAAVAVWTTPESVEAGAVFAEIGPRMAELSGSRLA
                     AQQQMEGLLAPHRPKEPAWFLATVGVSPDHQGKGLGSAVVLPGVEAAERAGVPAFLETS
                     APRNLPFYERLGFTVTADVEVPEGPRTWCMTRKPGA"
     CDS             6772..6825
                     /codon_start=1
                     /product="2A peptide from Thosea asigna virus capsid
                     protein"
                     /label=T2A
                     /note="Eukaryotic ribosomes fail to insert a peptide bond
                     between the Gly and Pro residues, yielding separate
                     polypeptides."
                     /translation="EGRGSLLTCGDVEENPGP"
     CDS             6826..7572
                     /codon_start=1
                     /product="improved tetracycline-controlled transactivator"
                     /label=rtTA-Advanced
                     /note="In the Tet-On(R) system, rtTA-Advanced binds to the
                     Tet-responsive element and stimulates transcription only in
                     the presence of tetracycline or doxycycline."
                     /translation="MSRLDKSKVINGALELLNGVGIEGLTTRKLAQKLGVEQPTLYWHV
                     KNKRALLDALPIEMLDRHHTHFCPLEGESWQDFLRNNAKSYRCALLSHRDGAKVHLGTR
                     PTEKQYETLENQLAFLCQQGFSLENALYALSAVGHFTLGCVLEEQEHQVAKEERETPTT
                     DSMPPLLRQAIELFDRQGAEPAFLFGLELIICGLEKQLKCESGGPTDALDDFDLDMLPA
                     DALDDFDLDMLPADALDDFDLDMLPG"
     CDS             complement(8030..8041)
                     /codon_start=1
                     /product="Factor Xa recognition and cleavage site"
                     /label=Factor Xa site
                     /translation="IEGR"
     LTR             8246..8479
                     /label=3' LTR (Delta-U3)
                     /note="self-inactivating 3' long terminal repeat (LTR) from
                     HIV-1"
     polyA_signal    8551..8672
                     /label=SV40 poly(A) signal
                     /note="SV40 polyadenylation signal"
     rep_origin      8712..8847
                     /label=SV40 ori
                     /note="SV40 origin of replication"
     promoter        8847..8918
                     /gene="bla"
                     /label=AmpR promoter
     CDS             8919..9779
                     /codon_start=1
                     /gene="bla"
                     /product="beta-lactamase"
                     /label=AmpR
                     /note="confers resistance to ampicillin, carbenicillin, and
                     related antibiotics"
                     /translation="MSIQHFRVALIPFFAAFCLPVFAHPETLVKVKDAEDQLGARVGYI
                     ELDLNSGKILESFRPEERFPMMSTFKVLLCGAVLSRIDAGQEQLGRRIHYSQNDLVEYS
                     PVTEKHLTDGMTVRELCSAAITMSDNTAANLLLTTIGGPKELTAFLHNMGDHVTRLDRW
                     EPELNEAIPNDERDTTMPVAMATTLRKLLTGELLTLASRQQLIDWMEADKVAGPLLRSA
                     LPAGWFIADKSGAGERGSRGIIAALGPDGKPSRIVVIYTTGSQATMDERNRQIAEIGAS
                     LIKHW"
ORIGIN
        1 ttgagatcct ttttttctgc gcgtaatctg ctgcttgcaa acaaaaaaac caccgctacc
```

61 agcggtggtt tgtttgccgg atcaagagct accaactctt tttccgaagg taactggctt
121 cagcagagcg cagataccaa atactgttct tctagtgtag ccgtagttag gccaccactt
181 caagaactct gtagcaccgc ctacatacct cgctctgcta atcctgttac cagtggctgc
241 tgccagtggc gataagtcgt gtcttaccgg gttggactca agacgatagt taccggataa
301 ggcgcagcgg tcgggctgaa cggggggttc gtgcacacag cccagcttgg agcgaacgac
361 ctacaccgaa ctgagatacc tacagcgtga gctatgagaa agcgccacgc ttcccgaagg
421 gagaaaggcg gacaggtatc cggtaagcgg cagggtcgga acaggagagc gcacgaggga
481 gcttccaggg ggaaacgcct ggtatcttta tagtcctgtc gggtttcgcc acctctgact
541 tgagcgtcga tttttgtgat gctcgtcagg ggggcggagc ctatggaaaa acgccagcaa
601 cgcggccttt ttacggttcc tggccttttg ctggcctttt gctcacatgt tctttcctgc
661 gttatcccct gattctgtgg ataaccgtat taccgccttt gagtgagctg ataccgctcg
721 ccgcagccga acgaccgagc gcagcgagtc agtgagcgag gaagcggaag agcgcccaat
781 acgcaaaccg cctctccccg cgcgttggcc gattcattaa tgcagctggc acgacaggtt
841 tcccgactgg aaagcgggca gtgagcgcaa cgcaattaat gtgagttagc tcactcatta
901 ggcaccccag gctttacact ttatgcttcc ggctcgtatg ttgtgtggaa ttgtgagcgg
961 ataacaattt cacacaggaa acagctatga ccatgattac gccaagcgcg caattaaccc
1021 tcactaaagg gaacaaaagc tggagctgca agcttaatgt agtcttatgc aatactcttg
1081 tagtcttgca acatggtaac gatgagttag caacatgcct tacaaggaga gaaaaagcac
1141 cgtgcatgcc gattggtgga agtaaggtgg tacgatcgtg ccttattagg aaggcaacag
1201 acgggtctga catggattgg acgaaccact gaattgccgc attgcagaga tattgtattt
1261 aagtgcctag ctcgatacat aaacgggtct ctctggttag accagatctg agcctgggag
1321 ctctctggct aactagggaa cccactgctt aagcctcaat aaagcttgcc ttgagtgctt
1381 caagtagtgt gtgcccgtct gttgtgtgac tctggtaact agagatccct cagacccttt
1441 tagtcagtgt ggaaaatctc tagcagtggc gcccgaacag ggacttgaaa gcgaaaggga
1501 aaccagagga gctctctcga cgcaggactc ggcttgctga agcgcgcacg gcaagaggcg
1561 aggggcggcg actggtgagt acgccaaaaa ttttgactag cggaggctag aaggagagag
1621 atgggtgcga gagcgtcagt attaagcggg ggagaattag atcgcgatgg gaaaaaattc
1681 ggttaaggcc agggggaaag aaaaaatata aattaaaaca tatagtatgg gcaagcaggg
1741 agctagaacg attcgcagtt aatcctggcc tgttagaaac atcagaaggc tgtagacaaa
1801 tactgggaca gctacaacca tcccttcaga caggatcaga agaacttaga tcattatata
1861 atacagtagc aaccctctat tgtgtgcatc aaaggataga gataaaagac accaaggaag
1921 ctttagacaa gatagaggaa gagcaaaaca aaagtaagac caccgcacag caagcggccg
1981 ctgatcttca gacctggagg aggagatatg agggacaatt ggagaagtga attatataaa
2041 tataaagtag taaaaattga accattagga gtagcaccca ccaaggcaaa gagaagagtg
2101 gtgcagagag aaaaaagagc agtgggaata ggagctttgt tccttgggtt cttgggagca
2161 gcaggaagca ctatgggcgc agcgtcaatg acgctgacgg tacaggccag acaattattg
2221 tctggtatag tgcagcagca gaacaatttg ctgagggcta ttgaggcgca acagcatctg
2281 ttgcaactca cagtctgggg catcaagcag ctccaggcaa gaatcctggc tgtggaaaga
2341 tacctaaagg atcaacagct cctggggatt tggggttgct ctggaaaact catttgcacc
2401 actgctgtgc cttggaatgc tagttggagt aataaatctc tggaacagat ttggaatcac
2461 acgacctgga tggagtggga cagagaaatt aacaattaca caagcttaat acactcctta
2521 attgaagaat cgcaaaacca gcaagaaaag aatgaacaag aattattgga attagataaa
2581 tgggcaagtt tgtggaattg gtttaacata acaaattggc tgtggtatat aaaattattc
2641 ataatgatag taggaggctt ggtaggttta agaatagttt ttgctgtact ttctatagtg
2701 aatagagtta ggcagggata ttcaccatta tcgtttcaga cccacctccc aaccccgagg
2761 ggacaattct cgacctcgag acaaatggca gtattcatcc acaattttaa aagaaaaggg
2821 gggattgggg ggtacagtgc aggggaaaga atagtagaca taatagcaac agacatacaa
2881 actaaagaat tacaaaaaca aattacaaaa attcaaaatt ttcgggttta ttacagggac
2941 agcagagatc cactttggcc gcgaatcgat atgtcgagtt tactccctat cagtgataga
3001 gaacgtatgt cgagtttact ccctatcagt gatagagaac gatgtcgagt ttactcccta
3061 tcagtgatag agaacgtatg tcgagtttac tccctatcag tgatagagaa cgtatgtcga
3121 gtttactccc tatcagtgat agagaacgta tgtcgagttt atccctatca gtgatagaga
3181 acgtatgtcg agtttactcc ctatcagtga tagagaacgt atgtcgaggt aggcgtgtac
3241 ggtgggaggc ctatataagc agagctcgtt tagtgaaccg tcagatcgcc tggagaattg
3301 gctagcatca acaagtttgt acaaaaaagc aggctgcacc atggtggcgg aacggtcccc
3361 cgctcgcagt cccggtagct ggctttttcc cgggttatgg cttttggtgt tgtctgggcc
3421 ggggggactc ctccgggctc aggaacagcc ctcttgtaga cgagcatttg atctgtattt
3481 cgtgctggac aaatctggct ccgtcgctaa taattggatt gagatatata actttgtgca
3541 gcagctggct gaacgcttcg tttcaccgga aatgagactg tccttcatcg tgtttagcag
3601 ccaggctact attatcctcc ctctcactgg agaccgggga aagatctcca agggtctcga
3661 agacctcaag cgggtgtccc cggtgggaga aacatacatc catgagggcc tcaaattggc
3721 caacgaacag atacagaaag cgggggggcct gaaaacctcc tccataatca ttgccctcac
3781 agatggaaaa ctggatgggc tcgttccctc ctatgccgag aaagaggcaa aatcagtag

3841 atccctcggc gcctccgtct attgcgtggg ggtgctggac ttcgaacaag cgcagctgga
3901 gcgcattgcc gacagcaagg aacaagtgtt ccctgtgaaa ggaggatttc aagctctcaa
3961 agggatcata aattccatct tggcgcagtc ttgtactgaa atccttgaat tacaaccatc
4021 ttccgtttgt gtcggtgagg agttccagat cgtgttgagt gggaggggct tcatgctggg
4081 gagccgaaat ggcagcgttc tctgcaccta taccgtgaac gaaacatata ctactagtgt
4141 aaaaccagtg agcgtgcagc tgaacagcat gctgtgtcct gcccccatcc tgaataaagc
4201 gggggagact ctcgatgtta gcgtcagttt taatgggggg aagagcgtga ttagtggaag
4261 tctcattgtt accgctacag aatgtagcaa cggtatagcc gctatcattg ttatcctggt
4321 tctcctgttg cttctgggga taggactcat gtggtggttc tggccgctgt gttgtaaggt
4381 ggtaattaag gacccacctc caccacccgc acccgctccg aaagaggagg aggaagaacc
4441 gctgcctact aagaaatggc ccaccgtgga cgcttcatac tatggcgggc gtggcgtggg
4501 cggcattaaa aggatggagg taaggtgggg agataaggga tcaactgaag agggtgcacg
4561 actggagaag gctaagaatg ctgtcgtaaa gattcctgaa gagacagagg agcctattag
4621 gcctagaccc cccagaccaa agcccacaca tcaaccgccc caaacaaagt ggtatacccc
4681 aattaagggt agactggatg cattatgggc cctgctgcgc cgccagtacg acagagtgtc
4741 tttaatgcgg ccacaggagg gcgatgaggg gagatgtata aatttttcac gagtcccatc
4801 tcagggtgga ggtggctccg tgtccaaggg cgaggaagat aacatggcca gcctcccagc
4861 tacacacgaa cttcacatat tcgggtcaat aaacggcgtc gacttcgaca tggtgggtca
4921 aggcaccgga aatccaaatg atgggtacga ggagctcaac cttaagagca ccaaggggga
4981 cctccagttc tctccatgga tcttagtccc tcacatagga tacggctttc atcagtacct
5041 gccctacccg gacggtatga gccctttcca ggccgccatg gtggacgggt cagggtacca
5101 ggtacacagg actatgcaat tcgaggacgg cgcatcactg accgtaaact accgatatac
5161 gtacgagggg agtcacataa agggcgaagc acaggtcaaa ggaaccgggt tccctgccga
5221 cgggcctgtc atgactaaca gcttaactgc agccgactgg tgtagatcca aaaagacata
5281 ccccaacgac aagacaatca ttagcacatt taagtggagc tacaccactg gaaatgggaa
5341 aagataccgc agtaccgcca gaaccactta taccttcgcc aagccaatgg cggccaatta
5401 tctgaagaac caacctatgt acgtgtttcg caagactgag ctgaaacatt ctaaaacaga
5461 attaaacttc aaggagtggc aaaaagcctt tactgacgtc atgggtatgg atgaattata
5521 taagggtgga ggtggctccg gcaagccaat ccctaaccct ctgttgggac tggatagcac
5581 ataggaccca gctttcttgt acaaagtggt ttagtaatga accggtccac caccaccacc
5641 accactaagg atccggggtt ggggttgcgc cttttccaag gcagccctgg gtttgcgcag
5701 ggacgcggct gctctgggcg tggttccggg aaacgcagcg gcgccgaccc tgggtctcgc
5761 acattcttca cgtccgttcg cagcgtcacc cggatcttcg ccgctaccct tgtgggcccc
5821 ccggcgacgc ttcctgctcc gcccctaagt cgggaaggtt ccttgcggtt cgcggcgtgc
5881 cggacgtgac aaacggaagc cgcacgtctc actagtaccc tcgcagacgg acagcgccag
5941 ggagcaatgg cagcgcgccg accgcgatgg gctgtggcca atagcggctg ctcagcaggg
6001 cgcgccgaga gcagcggccg ggaaggggcg gtgcgggagg cggggtgtgg ggcggtagtg
6061 tgggccctgt tcctgcccgc gcggtgttcc gcattctgca agcctccgga gcgcacgtcg
6121 gcagtcggct ccctcgttga ccgaatcacc gacctctctc cccagcaatt caccatgacc
6181 gagtacaagc ccacggtgcg cctcgccacc cgcgacgacg tccccagggc cgtacgcacc
6241 ctcgccgccg cgttcgccga ctaccccgcc acgcgccaca ccgtcgatcc ggaccgccac
6301 atcgagcggg tcaccgagct gcaagaactc ttcctcacgc gcgtcgggct cgacatcggc
6361 aaggtgtggg tcgcggacga cggcgccgcg gtggcggtct ggaccacgcc ggagagcgtc
6421 gaagcggggg cggtgttcgc cgagatcggc ccgcgcatgg ccgagttgag cggttcccgg
6481 ctggccgcgc agcaacagat ggaaggcctc ctggcgccgc accggcccaa ggagcccgcg
6541 tggttcctgg ccaccgtcgg cgtctcgccc gaccaccagg gcaagggtct gggcagcgcc
6601 gtcgtgctcc ccggagtgga ggcggccgag cgcgccgggg tgcccgcctt cctggagacc
6661 tccgcgcccc gcaacctccc cttctacgag cggctcggct tcaccgtcac cgccgacgtc
6721 gaggtgcccg aaggaccgcg cacctggtgc atgacccgca agcccggtgc cgaaggtaga
6781 ggttctctcc tcacttgtgg tgatgttgaa gaaaaccctg gtccaatgtc tagactggac
6841 aagagcaaag tcataaacgg agctctggaa ttactcaatg gtgtcggtat cgaaggcctg
6901 acgacaagga aactcgctca aaagctggga gttgagcagc ctaccctgta ctggcacgtg
6961 aagaacaagc gggccctgct cgatgccctg ccaatcgaga tgctggacag gcatcatacc
7021 cacttctgcc ccctggaagg cgagtcatgg caagactttc tgcggaacaa cgccaagtca
7081 taccgctgtg ctctcctctc acatcgcgac ggggctaaag tgcatctcgg cacccgccca
7141 acagagaaac agtacgaaac cctggaaaat cagctcgcgt tcctgtgtca gcaaggcttc
7201 tccctggaga acgcactgta cgctctgtcc gccgtgggcc actttacact gggctgcgta
7261 ttggaggaac aggagcatca agtagcaaaa gaggaaagag agacacctac caccgattct
7321 atgcccccac ttctgagaca agcaattgag ctgttcgacc ggcagggagc cgaacctgcc
7381 ttccttttcg gcctggaact aatcatatgt ggcctggaga aacagctaaa gtgcgaaagc
7441 ggcgggccga ccgacgccct tgacgatttt gacttagaca tgctcccagc cgatgccctt
7501 gacgactttg accttgatat gctgcctgct gacgctcttg acgattttga ccttgacatg
7561 ctccccgggt aactaagtaa ggatcgatcc aagatatcgt attcttaact atgttgctcc

```
7621 ttttacgcta tgtggatacg ctgctttaat gcctttgtat catgctattg cttcccgtat
7681 ggctttcatt ttctcctcct tgtataaatc ctggttgctg tctctttatg aggagttgtg
7741 gcccgttgtc aggcaacgtg gcgtggtgtg cactgtgttt gctgacgcaa cccccactgg
7801 ttggggcatt gccaccacct gtcagctcct ttccgggact ttcgctttcc ccctccctat
7861 tgccacggcg gaactcatcg ccgcctgcct tgcccgctgc tggacagggg ctcggctgtt
7921 gggcactgac aattccgtgg tgttgtcggg gaagctgacg tcctttccat ggctgctcgc
7981 ctgtgttgcc acctggattc tgcgcgggac gtccttctgc tacgtccctt cggccctcaa
8041 tccagcggac cttccttccc gcggcctgct gccggctctg cggcctcttc cgcgtcttcg
8101 ccttcgccct cagacgagtc ggatctccct ttgggccgcc tccccgcctg tttcgcctcg
8161 gcgtccggac tagaggtacc tttaagacca atgacttaca aggcagctgt agatcttagc
8221 cactttttaa aagaaaaggg gggactggaa gggctaattc actcccaacg aagacaagat
8281 ctgctttttg cttgtactgg gtctctctgg ttagaccaga tctgagcctg ggagctctct
8341 ggctaactag ggaacccact gcttaagcct caataaagct tgccttgagt gcttcaagta
8401 gtgtgtgccc gtctgttgtg tgactctggt aactagagat ccctcagacc cttttagtca
8461 gtgtggaaaa tctctagcag tagtagttca tgtcatctta ttattcagta tttataactt
8521 gcaaagaaat gaatatcaga gagtgagagg aacttgttta ttgcagctta taatggttac
8581 aaataaagca atagcatcac aaatttcaca aataaagcat ttttttcact gcattctagt
8641 tgtggtttgt ccaaactcat caatgtatct tatcatgtct ggctctagct atcccgcccc
8701 taactccgcc catcccgccc ctaactccgc ccagttccgc ccattctccg ccccatggct
8761 gactaatttt ttttatttat gcagaggccg aggccgcctc ggcctctgag ctattccaga
8821 agtagtgagg aggctttttt ggaggccttt caaatatgta tccgctcatg agacaataac
8881 cctgataaat gcttcaataa tattgaaaaa ggaagagtat gagtattcaa catttccgtg
8941 tcgcccttat tccctttttt gcggcatttt gccttcctgt ttttgctcac ccagaaacgc
9001 tggtgaaagt aaaagatgct gaagatcagt tgggtgcacg agtgggttac atcgaactgg
9061 atctcaacag cggtaagatc cttgagagtt ttcgccccga agaacgtttt ccaatgatga
9121 gcacttttaa agttctgcta tgtggcgcgg tattatcccg tattgacgcc gggcaagagc
9181 aactcggtcg ccgcatacac tattctcaga atgacttggt tgagtactca ccagtcacag
9241 aaaagcatct tacggatggc atgacagtaa gagaattatg cagtgctgcc ataaccatga
9301 gtgataacac tgcggccaac ttacttctga caacgatcgg aggaccgaag gagctaaccg
9361 cttttttgca caacatgggg gatcatgtaa ctcgccttga tcgttgggaa ccggagctga
9421 atgaagccat accaaacgac gagcgtgaca ccacgatgcc tgtagcaatg gcaacaacgt
9481 tgcgcaaact attaactggc gaactactta ctctagcttc ccggcaacaa ttaatagact
9541 ggatggaggc ggataaagtt gcaggaccac ttctgcgctc ggcccttccg gctggctggt
9601 ttattgctga taaatctgga gccggtgagc gtgggtctcg cggtatcatt gcagcactgg
9661 ggccagatgg taagccctcc cgtatcgtag ttatctacac gacggggagt caggcaacta
9721 tggatgaacg aaatagacag atcgctgaga taggtgcctc actgattaag cattggtaac
9781 tgtcagacca agtttactca tatatacttt agattgattt aaaacttcat ttttaattta
9841 aaaggatcta ggtgaagatc ctttttgata atctcatgac caaaatccct taacgtgagt
9901 tttcgttcca ctgagcgtca gaccccgtag aaaagatcaa aggatcttc
```
//

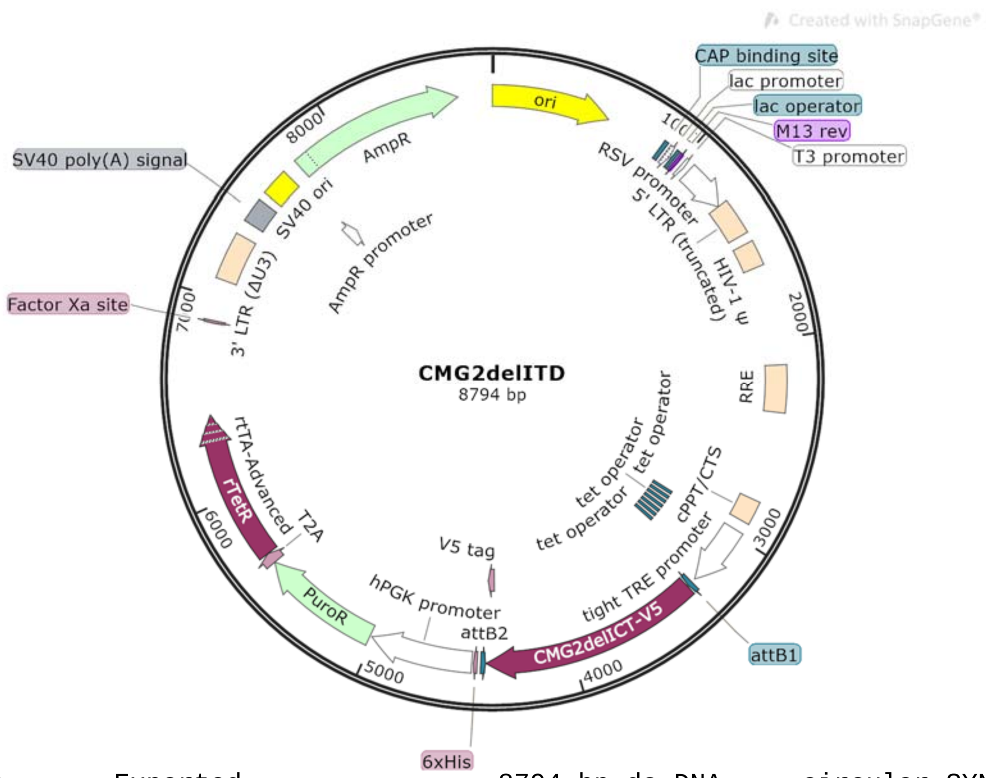


```
LOCUS       Exported                8794 bp ds-DNA     circular SYN 14-JUL-2025
DEFINITION  synthetic circular DNA
ACCESSION   .
VERSION     .
KEYWORDS    .
SOURCE      synthetic DNA construct
  ORGANISM  synthetic DNA construct
REFERENCE   1  (bases 1 to 8794)
  AUTHORS   Vascular Biology Program
  TITLE     Direct Submission
  JOURNAL   Exported Jul 14, 2025 from SnapGene 4.2.11
            http://www.snapgene.com
FEATURES             Location/Qualifiers
     source          1..8794
                     /organism="synthetic DNA construct"
                     /mol_type="other DNA"
     rep_origin      1..589
                     /direction=RIGHT
                     /label=ori
                     /note="high-copy-number ColE1/pMB1/pBR322/pUC origin of
                     replication"
     protein_bind    877..898
                     /label=CAP binding site
                     /bound_moiety="E. coli catabolite activator protein"
                     /note="CAP binding activates transcription in the presence
                     of cAMP."
     promoter        913..943
                     /label=lac promoter
                     /note="promoter for the E. coli lac operon"
     protein_bind    951..967
                     /label=lac operator
                     /bound_moiety="lac repressor encoded by lacI"
                     /note="The lac repressor binds to the lac operator to
                     inhibit transcription in E. coli. This inhibition can be
                     relieved by adding lactose or
                     isopropyl-beta-D-thiogalactopyranoside (IPTG)."
     primer_bind     975..991
                     /label=M13 rev
```

```
                     /note="common sequencing primer, one of multiple similar
                     variants"
     promoter        1012..1030
                     /label=T3 promoter
                     /note="promoter for bacteriophage T3 RNA polymerase"
     promoter        1058..1284
                     /label=RSV promoter
                     /note="Rous sarcoma virus enhancer/promoter"
     LTR             1285..1465
                     /label=5' LTR (truncated)
                     /note="truncated 5' long terminal repeat (LTR) from HIV-1"
     misc_feature    1512..1637
                     /label=HIV-1 Psi
                     /note="packaging signal of human immunodeficiency virus
                     type 1"
     misc_feature    2130..2363
                     /label=RRE
                     /note="The Rev response element (RRE) of HIV-1 allows for
                     Rev-dependent mRNA export from the nucleus to the
                     cytoplasm."
     misc_feature    2805..2922
                     /label=cPPT/CTS
                     /note="central polypurine tract and central termination
                     sequence of HIV-1"
     promoter        2976..3290
                     /label=tight TRE promoter
                     /note="Tet-responsive promoter PTight, consisting of seven
                     tet operator sequences followed by the minimal CMV
                     promoter"
     protein_bind    2984..3002
                     /gene="tetO"
                     /label=tet operator
                     /bound_moiety="tetracycline repressor TetR"
                     /note="bacterial operator O2 for the tetR and tetA genes"
     protein_bind    3020..3038
                     /gene="tetO"
                     /label=tet operator
                     /bound_moiety="tetracycline repressor TetR"
                     /note="bacterial operator O2 for the tetR and tetA genes"
     protein_bind    3055..3073
                     /gene="tetO"
                     /label=tet operator
                     /bound_moiety="tetracycline repressor TetR"
                     /note="bacterial operator O2 for the tetR and tetA genes"
     protein_bind    3091..3109
                     /gene="tetO"
                     /label=tet operator
                     /bound_moiety="tetracycline repressor TetR"
                     /note="bacterial operator O2 for the tetR and tetA genes"
     protein_bind    3127..3145
                     /gene="tetO"
                     /label=tet operator
                     /bound_moiety="tetracycline repressor TetR"
                     /note="bacterial operator O2 for the tetR and tetA genes"
     protein_bind    3162..3180
                     /gene="tetO"
                     /label=tet operator
                     /bound_moiety="tetracycline repressor TetR"
                     /note="bacterial operator O2 for the tetR and tetA genes"
     protein_bind    3198..3216
                     /gene="tetO"
                     /label=tet operator
                     /bound_moiety="tetracycline repressor TetR"
```

```
                     /note="bacterial operator O2 for the tetR and tetA genes"
     protein_bind    3311..3335
                     /gene="mutant version of attB"
                     /label=attB1
                     /bound_moiety="BP Clonase(TM)"
                     /note="recombination site for the Gateway(R) BP reaction"
     CDS             3341..4429
                     /codon_start=1
                     /label=CMG2delICT-V5
                     /translation="MVAERSPARSPGSWLFPGLWLLVLSGPGGLLRAQEQPSCRRAFDL
                     YFVLDKSGSVANNWIEIYNFVQQLAERFVSPEMRLSFIVFSSQATIILPLTGDRGKISK
                     GLEDLKRVSPVGETYIHEGLKLANEQIQKAGGLKTSSIIIALTDGKLDGLVPSYAEKEA
                     KISRSLGASVYCVGVLDFEQAQLERIADSKEQVFPVKGGFQALKGIINSILAQSCTEIL
                     ELQPSSVCVGEEFQIVLSGRGFMLGSRNGSVLCTYTVNETYTTSVKPVSVQLNSMLCPA
                     PILNKAGETLDVSVSFNGGKSVISGSLIVTATECSNGIAAIIVILVLLLLLGIGLMWWF
                     WPLGGGGSGKPIPNPLLGLDST"
     CDS             4385..4426
                     /codon_start=1
                     /product="epitope tag from simian virus 5"
                     /label=V5 tag
                     /translation="GKPIPNPLLGLDST"
     protein_bind    complement(4431..4455)
                     /gene="mutant version of attB"
                     /label=attB2
                     /bound_moiety="BP Clonase(TM)"
                     /note="recombination site for the Gateway(R) BP reaction"
     CDS             4473..4490
                     /codon_start=1
                     /product="6xHis affinity tag"
                     /label=6xHis
                     /translation="HHHHHH"
     promoter        4500..5010
                     /label=hPGK promoter
                     /note="human phosphoglycerate kinase 1 promoter"
     CDS             5020..5616
                     /codon_start=1
                     /gene="pac from Streptomyces alboniger"
                     /product="puromycin N-acetyltransferase"
                     /label=PuroR
                     /note="confers resistance to puromycin"
                     /translation="MTEYKPTVRLATRDDVPRAVRTLAAAFADYPATRHTVDPDRHIER
                     VTELQELFLTRVGLDIGKVWVADDGAAVAVWTTPESVEAGAVFAEIGPRMAELSGSRLA
                     AQQQMEGLLAPHRPKEPAWFLATVGVSPDHQGKGLGSAVVLPGVEAAERAGVPAFLETS
                     APRNLPFYERLGFTVTADVEVPEGPRTWCMTRKPGA"
     CDS             5617..5670
                     /codon_start=1
                     /product="2A peptide from Thosea asigna virus capsid
                     protein"
                     /label=T2A
                     /note="Eukaryotic ribosomes fail to insert a peptide bond
                     between the Gly and Pro residues, yielding separate
                     polypeptides."
                     /translation="EGRGSLLTCGDVEENPGP"
     CDS             5671..6417
                     /codon_start=1
                     /product="improved tetracycline-controlled transactivator"
                     /label=rtTA-Advanced
                     /note="In the Tet-On(R) system, rtTA-Advanced binds to the
                     Tet-responsive element and stimulates transcription only in
                     the presence of tetracycline or doxycycline."
                     /translation="MSRLDKSKVINGALELLNGVGIEGLTTRKLAQKLGVEQPTLYWHV
                     KNKRALLDALPIEMLDRHHTHFCPLEGESWQDFLRNNAKSYRCALLSHRDGAKVHLGTR
                     PTEKQYETLENQLAFLCQQGFSLENALYALSAVGHFTLGCVLEEQEHQVAKEERETPTT
```

```
                     DSMPPLLRQAIELFDRQGAEPAFLFGLELIICGLEKQLKCESGGPTDALDDFDLDMLPA
                     DALDDFDLDMLPADALDDFDLDMLPG"
     CDS             complement(6875..6886)
                     /codon_start=1
                     /product="Factor Xa recognition and cleavage site"
                     /label=Factor Xa site
                     /translation="IEGR"
     LTR             7091..7324
                     /label=3' LTR (Delta-U3)
                     /note="self-inactivating 3' long terminal repeat (LTR) from
                     HIV-1"
     polyA_signal    7396..7517
                     /label=SV40 poly(A) signal
                     /note="SV40 polyadenylation signal"
     rep_origin      7557..7692
                     /label=SV40 ori
                     /note="SV40 origin of replication"
     promoter        7692..7763
                     /gene="bla"
                     /label=AmpR promoter
     CDS             7764..8624
                     /codon_start=1
                     /gene="bla"
                     /product="beta-lactamase"
                     /label=AmpR
                     /note="confers resistance to ampicillin, carbenicillin, and
                     related antibiotics"
                     /translation="MSIQHFRVALIPFFAAFCLPVFAHPETLVKVKDAEDQLGARVGYI
                     ELDLNSGKILESFRPEERFPMMSTFKVLLCGAVLSRIDAGQEQLGRRIHYSQNDLVEYS
                     PVTEKHLTDGMTVRELCSAAITMSDNTAANLLLTTIGGPKELTAFLHNMGDHVTRLDRW
                     EPELNEAIPNDERDTTMPVAMATTLRKLLTGELLTLASRQQLIDWMEADKVAGPLLRSA
                     LPAGWFIADKSGAGERGSRGIIAALGPDGKPSRIVVIYTTGSQATMDERNRQIAEIGAS
                     LIKHW"
ORIGIN
        1 ttgagatcct ttttttctgc gcgtaatctg ctgcttgcaa acaaaaaaac caccgctacc
       61 agcggtggtt tgtttgccgg atcaagagct accaactctt tttccgaagg taactggctt
      121 cagcagagcg cagataccaa atactgttct tctagtgtag ccgtagttag gccaccactt
      181 caagaactct gtagcaccgc ctacatacct cgctctgcta atcctgttac cagtggctgc
      241 tgccagtggc gataagtcgt gtcttaccgg gttggactca agacgatagt taccggataa
      301 ggcgcagcgg tcgggctgaa cggggggttc gtgcacacag cccagcttgg agcgaacgac
      361 ctacaccgaa ctgagatacc tacagcgtga gctatgagaa agcgccacgc ttcccgaagg
      421 gagaaaggcg gacaggtatc cggtaagcgg cagggtcgga acaggagagc gcacgaggga
      481 gcttccaggg ggaaacgcct ggtatcttta tagtcctgtc gggtttcgcc acctctgact
      541 tgagcgtcga tttttgtgat gctcgtcagg ggggcggagc ctatggaaaa acgccagcaa
      601 cgcggccttt ttacggttcc tggccttttg ctggcctttt gctcacatgt tctttcctgc
      661 gttatcccct gattctgtgg ataaccgtat taccgccttt gagtgagctg ataccgctcg
      721 ccgcagccga acgaccgagc gcagcgagtc agtgagcgag gaagcggaag agcgcccaat
      781 acgcaaaccg cctctccccg cgcgttggcc gattcattaa tgcagctggc acgacaggtt
      841 tcccgactgg aaagcgggca gtgagcgcaa cgcaattaat gtgagttagc tcactcatta
      901 ggcaccccag gctttacact ttatgcttcc ggctcgtatg ttgtgtggaa ttgtgagcgg
      961 ataacaattt cacacaggaa acagctatga ccatgattac gccaagcgcg caattaaccc
     1021 tcactaaagg gaacaaaagc tggagctgca agcttaatgt agtcttatgc aatactcttg
     1081 tagtcttgca acatggtaac gatgagttag caacatgcct tacaaggaga gaaaaagcac
     1141 cgtgcatgcc gattggtgga agtaaggtgg tacgatcgtg ccttattagg aaggcaacag
     1201 acgggtctga catggattgg acgaaccact gaattgccgc attgcagaga tattgtattt
     1261 aagtgcctag ctcgatacat aaacgggtct ctctggttag accagatctg agcctgggag
     1321 ctctctggct aactagggaa cccactgctt aagcctcaat aaagcttgcc ttgagtgctt
     1381 caagtagtgt gtgcccgtct gttgtgtgac tctggtaact agagatccct cagacccttt
     1441 tagtcagtgt ggaaaatctc tagcagtggc gcccgaacag ggacttgaaa gcgaaaggga
     1501 aaccagagga gctctctcga cgcaggactc ggcttgctga agcgcgcacg gcaagaggcg
     1561 aggggcggcg actggtgagt acgccaaaaa ttttgactag cggaggctag aaggagagag
     1621 atgggtgcga gagcgtcagt attaagcggg ggagaattag atcgcgatgg gaaaaaattc
     1681 ggttaaggcc agggggaaag aaaaaatata aattaaaaca tatagtatgg gcaagcaggg
```

1741 agctagaacg attcgcagtt aatcctggcc tgttagaaac atcagaaggc tgtagacaaa
1801 tactgggaca gctacaacca tcccttcaga caggatcaga agaacttaga tcattatata
1861 atacagtagc aaccctctat tgtgtgcatc aaaggataga gataaaagac accaaggaag
1921 ctttagacaa gatagaggaa gagcaaaaca aaagtaagac caccgcacag caagcggccg
1981 ctgatcttca gacctggagg aggagatatg agggacaatt ggagaagtga attatataaa
2041 tataaagtag taaaaattga accattagga gtagcaccca ccaaggcaaa gagaagagtg
2101 gtgcagagag aaaaaagagc agtgggaata ggagctttgt tccttgggtt cttgggagca
2161 gcaggaagca ctatgggcgc agcgtcaatg acgctgacgg tacaggccag acaattattg
2221 tctggtatag tgcagcagca gaacaatttg ctgagggcta ttgaggcgca acagcatctg
2281 ttgcaactca cagtctgggg catcaagcag ctccaggcaa gaatcctggc tgtggaaaga
2341 tacctaaagg atcaacagct cctggggatt tggggttgct ctggaaaact catttgcacc
2401 actgctgtgc cttggaatgc tagttggagt aataaatctc tggaacagat ttggaatcac
2461 acgacctgga tggagtggga cagagaaatt aacaattaca caagcttaat acactcctta
2521 attgaagaat cgcaaaacca gcaagaaaag aatgaacaag aattattgga attagataaa
2581 tgggcaagtt tgtggaattg gtttaacata acaaattggc tgtggtatat aaaattattc
2641 ataatgatag taggaggctt ggtaggttta agaatagttt ttgctgtact ttctatagtg
2701 aatagagtta ggcagggata ttcaccatta tcgtttcaga cccacctccc aaccccgagg
2761 ggacaattct cgacctcgag acaaatggca gtattcatcc acaattttaa aagaaaaggg
2821 gggattgggg ggtacagtgc aggggaaaga atagtagaca taatagcaac agacatacaa
2881 actaaagaat tacaaaaaca aattacaaaa attcaaaatt ttcgggttta ttacagggac
2941 agcagagatc cactttggcc gcgaatcgat atgtcgagtt tactccctat cagtgataga
3001 gaacgtatgt cgagtttact ccctatcagt gatagagaac gatgtcgagt ttactcccta
3061 tcagtgatag agaacgtatg tcgagtttac tccctatcag tgatagagaa cgtatgtcga
3121 gtttactccc tatcagtgat agagaacgta tgtcgagttt atccctatca gtgatagaga
3181 acgtatgtcg agtttactcc ctatcagtga tagagaacgt atgtcgaggt aggcgtgtac
3241 ggtgggaggc ctatataagc agagctcgtt tagtgaaccg tcagatcgcc tggagaattg
3301 gctagcatca acaagtttgt acaaaaaagc aggctgcacc atggtagctg agagatcacc
3361 cgcgcgtagc ccaggctctt ggctttttcc tggtttatgg ttgctggtgc tgagtgggcc
3421 cgggggactt ctgcgagcac aggagcagcc aagttgcaga cgggcttttg atttatattt
3481 cgtattggat aaatctggat ccgttgctaa taactggatt gagatctata actttgtaca
3541 gcagctcgct gaacgtttcg tgagcccgga gatgcgattg tcctttatcg tattcagcag
3601 ccaggcaacc ataattttgc ccctgactgg agaccgcggt aagatttcta agggactgga
3661 ggatttgaaa cgcgtttccc ccgtgggtga gacctacatt cacgaggggc tcaagctggc
3721 aaatgaacag atccaaaaag ctggcggcct taagacctct tccatcatta tcgctctcac
3781 cgacgggaag ctcgacggcc tcgtgccctc ctatgccgaa aaggaagcta agatctcaag
3841 gtctcttggg gcgtcagtgt actgtgttgg agtcctcgat tttgagcaag ctcagctcga
3901 gagaatcgcc gatagtaaag agcaagtgtt tcccgtgaag ggaggatttc aggctctcaa
3961 gggtatcatc aattccatcc tggctcagtc ctgcacagaa atactggagc tgcaaccatc
4021 tagtgtatgc gttggggagg agtttcagat agttctgagt gggcgcggat tcatgctcgg
4081 ctctcggaac ggcagcgtgc tgtgcacata cacggtgaac gagacttaca ctacaagtgt
4141 taagcccgta tccgttcagc tcaacagtat gctgtgccca gcccccatcc tgaacaaagc
4201 aggggagact ctcgacgtta gcgtgtcctt caacggaggg aaatccgtga tctctgggtc
4261 cctgatcgtt acagcgacgg aatgttccaa tggaatcgca gcaatcattg taatactggt
4321 cttgcttctc ctgctgggca tcggtcttat gtggtggttc tggccactgg gtggaggtgg
4381 ctccggcaag ccaatcccta accctctgtt gggactggat agcacatagg acccagcttt
4441 cttgtacaaa gtggtttagt aatgaaccgg tccaccacca ccaccaccac taaggatccg
4501 gggttggggt tgcgcctttt ccaaggcagc cctgggtttg cgcagggacg cggctgctct
4561 gggcgtggtt ccgggaaacg cagcggcgcc gaccctgggt ctcgcacatt cttcacgtcc
4621 gttcgcagcg tcacccggat cttcgccgct acccttgtgg gccccccggc gacgcttcct
4681 gctccgcccc taagtcggga aggttccttg cggttcgcgg cgtgccggac gtgacaaacg
4741 gaagccgcac gtctcactag taccctcgca gacggacagc gccagggagc aatggcagcg
4801 cgccgaccgc gatgggctgt ggccaatagc ggctgctcag cagggcgcgc cgagagcagc
4861 ggccgggaag gggcggtgcg ggaggcgggg tgtggggcgg tagtgtgggc cctgttcctg
4921 cccgcgcggt gttccgcatt ctgcaagcct ccggagcgca cgtcggcagt cggctccctc
4981 gttgaccgaa tcaccgacct ctctccccag caattcacca tgaccgagta caagcccacg
5041 gtgcgcctcg ccacccgcga cgacgtcccc agggccgtac gcaccctcgc cgccgcgttc
5101 gccgactacc ccgccacgcg ccacaccgtc gatccggacc gccacatcga gcgggtcacc
5161 gagctgcaag aactcttcct cacgcgcgtc gggctcgaca tcggcaaggt gtgggtcgcg
5221 gacgacggcg ccgcggtggc ggtctggacc acgccggaga gcgtcgaagc gggggcggtg
5281 ttcgccgaga tcggcccgcg catggccgag ttgagcggtt cccggctggc cgcgcagcaa
5341 cagatggaag gcctcctggc gccgcaccgg cccaaggagc ccgcgtggtt cctggccacc
5401 gtcggcgtct cgcccgacca ccagggcaag ggtctgggca gcgccgtcgt gctccccgga
5461 gtggaggcgg ccgagcgcgc cggggtgccc gccttcctgg agacctccgc gccccgcaac

5521 ctccccttct acgagcggct cggcttcacc gtcaccgccg acgtcgaggt gcccgaagga
5581 ccgcgcacct ggtgcatgac ccgcaagccc ggtgccgaag gtagaggttc tctcctcact
5641 tgtggtgatg ttgaagaaaa ccctggtcca atgtctagac tggacaagag caaagtcata
5701 aacggagctc tggaattact caatggtgtc ggtatcgaag gcctgacgac aaggaaactc
5761 gctcaaaagc tgggagttga gcagcctacc ctgtactggc acgtgaagaa caagcgggcc
5821 ctgctcgatg ccctgccaat cgagatgctg gacaggcatc atacccactt ctgcccccctg
5881 gaaggcgagt catggcaaga ctttctgcgg aacaacgcca agtcataccg ctgtgctctc
5941 ctctcacatc gcgacggggc taaagtgcat ctcggcaccc gcccaacaga gaaacagtac
6001 gaaaccctgg aaaatcagct cgcgttcctg tgtcagcaag gcttctccct ggagaacgca
6061 ctgtacgctc tgtccgccgt gggccacttt acactgggct gcgtattgga ggaacaggag
6121 catcaagtag caaaagagga aagagagaca cctaccaccg attctatgcc cccacttctg
6181 agacaagcaa ttgagctgtt cgaccggcag ggagccgaac ctgccttcct tttcggcctg
6241 gaactaatca tatgtggcct ggagaaacag ctaaagtgcg aaagcggcgg gccgaccgac
6301 gcccttgacg attttgactt agacatgctc ccagccgatg cccttgacga ctttgacctt
6361 gatatgctgc ctgctgacgc tcttgacgat tttgaccttg acatgctccc cgggtaacta
6421 agtaaggatc gatccaagat atcgtattct taactatgtt gctcctttta cgctatgtgg
6481 atacgctgct ttaatgcctt tgtatcatgc tattgcttcc cgtatggctt tcattttctc
6541 ctccttgtat aaatcctggt tgctgtctct ttatgaggag ttgtggcccg ttgtcaggca
6601 acgtggcgtg gtgtgcactg tgtttgctga cgcaaccccc actggttggg gcattgccac
6661 cacctgtcag ctcctttccg ggactttcgc tttccccctc cctattgcca cggcggaact
6721 catcgccgcc tgccttgccc gctgctggac aggggctcgg ctgttgggca ctgacaattc
6781 cgtggtgttg tcggggaagc tgacgtcctt tccatggctg ctcgcctgtg ttgccacctg
6841 gattctgcgc gggacgtcct tctgctacgt cccttcggcc ctcaatccag cggaccttcc
6901 ttcccgcggc ctgctgccgg ctctgcggcc tcttccgcgt cttcgccttc gccctcagac
6961 gagtcggatc tccctttggg ccgcctcccc gcctgtttcg cctcggcgtc cggactagag
7021 gtacctttaa gaccaatgac ttacaaggca gctgtagatc ttagccactt tttaaaagaa
7081 aaggggggac tggaagggct aattcactcc caacgaagac aagatctgct ttttgcttgt
7141 actgggtctc tctggttaga ccagatctga gcctgggagc tctctggcta actagggaac
7201 ccactgctta agcctcaata aagcttgcct tgagtgcttc aagtagtgtg tgcccgtctg
7261 ttgtgtgact ctggtaacta gagatccctc agaccctttt agtcagtgtg gaaaatctct
7321 agcagtagta gttcatgtca tcttattatt cagtatttat aacttgcaaa gaaatgaata
7381 tcagagagtg agaggaactt gtttattgca gcttataatg gttacaaata aagcaatagc
7441 atcacaaatt tcacaaataa agcatttttt tcactgcatt ctagttgtgg tttgtccaaa
7501 ctcatcaatg tatcttatca tgtctggctc tagctatccc gcccctaact ccgcccatcc
7561 cgcccctaac tccgcccagt tccgcccatt ctccgcccca tggctgacta attttttttta
7621 tttatgcaga ggccgaggcc gcctcggcct ctgagctatt ccagaagtag tgaggaggct
7681 tttttggagg cctttcaaat atgtatccgc tcatgagaca ataaccctga taaatgcttc
7741 aataatattg aaaaaggaag agtatgagta ttcaacattt ccgtgtcgcc cttattccct
7801 tttttgcggc attttgcctt cctgtttttg ctcacccaga aacgctggtg aaagtaaaag
7861 atgctgaaga tcagttgggt gcacgagtgg gttacatcga actggatctc aacagcggta
7921 agatccttga gagttttcgc cccgaagaac gttttccaat gatgagcact tttaaagttc
7981 tgctatgtgg cgcggtatta tcccgtattg acgccgggca agagcaactc ggtcgccgca
8041 tacactattc tcagaatgac ttggttgagt actcaccagt cacagaaaag catcttacgg
8101 atggcatgac agtaagagaa ttatgcagtg ctgccataac catgagtgat aacactgcgg
8161 ccaacttact tctgacaacg atcggaggac cgaaggagct aaccgctttt ttgcacaaca
8221 tgggggatca tgtaactcgc cttgatcgtt gggaaccgga gctgaatgaa gccataccaa
8281 acgacgagcg tgacaccacg atgcctgtag caatggcaac aacgttgcgc aaactattaa
8341 ctggcgaact acttactcta gcttcccggc aacaattaat agactggatg gaggcggata
8401 aagttgcagg accacttctg cgctcggccc ttccggctgg ctggtttatt gctgataaat
8461 ctggagccgg tgagcgtggg tctcgcggta tcattgcagc actggggcca gatggtaagc
8521 cctcccgtat cgtagttatc tacacgacgg ggagtcaggc aactatggat gaacgaaata
8581 gacagatcgc tgagataggt gcctcactga ttaagcattg gtaactgtca gaccaagttt
8641 actcatatat actttagatt gatttaaaac ttcattttta atttaaaagg atctaggtga
8701 agatcctttt tgataatctc atgaccaaaa tcccttaacg tgagttttcg ttccactgag
8761 cgtcagaccc cgtagaaaag atcaaaggat cttc
//

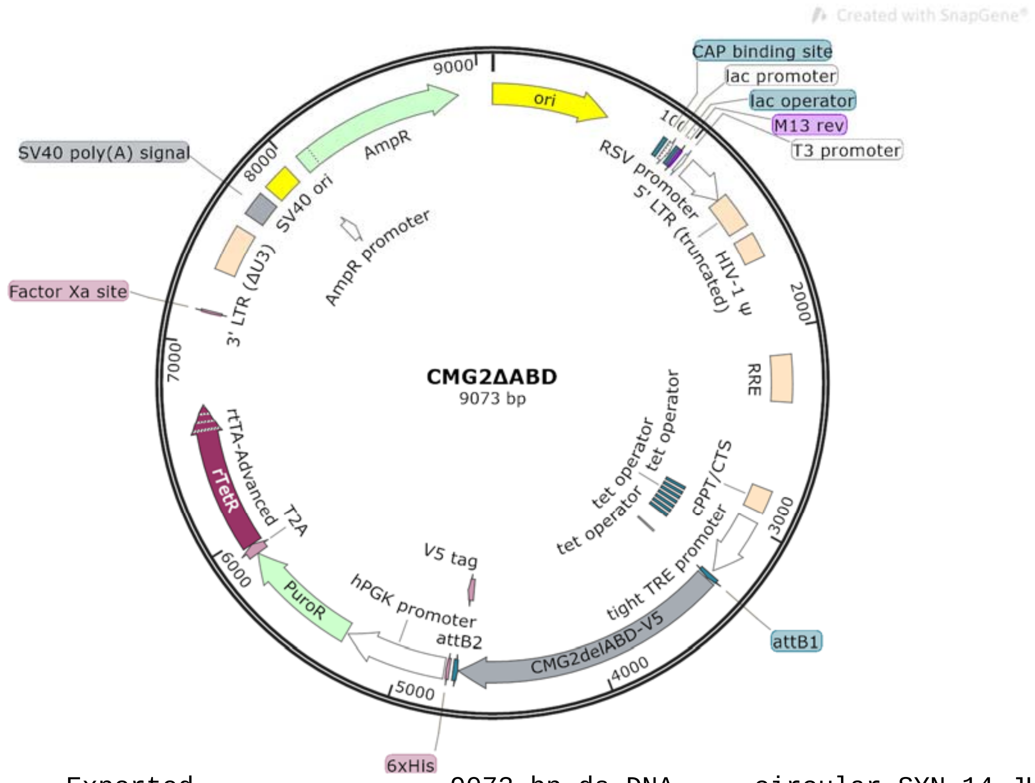


```
LOCUS       Exported                9073 bp ds-DNA     circular SYN 14-JUL-2025
DEFINITION  synthetic circular DNA
ACCESSION   .
VERSION     .
KEYWORDS    CMG2-Delta-ABD
SOURCE      synthetic DNA construct
  ORGANISM  synthetic DNA construct
REFERENCE   1  (bases 1 to 9073)
  AUTHORS   Vascular Biology Program
  TITLE     Direct Submission
  JOURNAL   Exported Jul 14, 2025 from SnapGene 4.2.11
            http://www.snapgene.com
FEATURES             Location/Qualifiers
     source          1..9073
                     /organism="synthetic DNA construct"
                     /mol_type="other DNA"
     rep_origin      1..589
                     /direction=RIGHT
                     /label=ori
                     /note="high-copy-number ColE1/pMB1/pBR322/pUC origin of
                     replication"
     protein_bind    877..898
                     /label=CAP binding site
                     /bound_moiety="E. coli catabolite activator protein"
                     /note="CAP binding activates transcription in the presence
                     of cAMP."
     promoter        913..943
                     /label=lac promoter
                     /note="promoter for the E. coli lac operon"
     protein_bind    951..967
                     /label=lac operator
                     /bound_moiety="lac repressor encoded by lacI"
                     /note="The lac repressor binds to the lac operator to
                     inhibit transcription in E. coli. This inhibition can be
                     relieved by adding lactose or
                     isopropyl-beta-D-thiogalactopyranoside (IPTG)."
     primer_bind     975..991
```

```
                     /label=M13 rev
                     /note="common sequencing primer, one of multiple similar
                     variants"
     promoter        1012..1030
                     /label=T3 promoter
                     /note="promoter for bacteriophage T3 RNA polymerase"
     promoter        1058..1284
                     /label=RSV promoter
                     /note="Rous sarcoma virus enhancer/promoter"
     LTR             1285..1465
                     /label=5' LTR (truncated)
                     /note="truncated 5' long terminal repeat (LTR) from HIV-1"
     misc_feature    1512..1637
                     /label=HIV-1 Psi
                     /note="packaging signal of human immunodeficiency virus
                     type 1"
     misc_feature    2130..2363
                     /label=RRE
                     /note="The Rev response element (RRE) of HIV-1 allows for
                     Rev-dependent mRNA export from the nucleus to the
                     cytoplasm."
     misc_feature    2805..2922
                     /label=cPPT/CTS
                     /note="central polypurine tract and central termination
                     sequence of HIV-1"
     promoter        2976..3290
                     /label=tight TRE promoter
                     /note="Tet-responsive promoter PTight, consisting of seven
                     tet operator sequences followed by the minimal CMV
                     promoter"
     protein_bind    2984..3002
                     /gene="tetO"
                     /label=tet operator
                     /bound_moiety="tetracycline repressor TetR"
                     /note="bacterial operator O2 for the tetR and tetA genes"
     protein_bind    3020..3038
                     /gene="tetO"
                     /label=tet operator
                     /bound_moiety="tetracycline repressor TetR"
                     /note="bacterial operator O2 for the tetR and tetA genes"
     protein_bind    3055..3073
                     /gene="tetO"
                     /label=tet operator
                     /bound_moiety="tetracycline repressor TetR"
                     /note="bacterial operator O2 for the tetR and tetA genes"
     protein_bind    3091..3109
                     /gene="tetO"
                     /label=tet operator
                     /bound_moiety="tetracycline repressor TetR"
                     /note="bacterial operator O2 for the tetR and tetA genes"
     protein_bind    3127..3145
                     /gene="tetO"
                     /label=tet operator
                     /bound_moiety="tetracycline repressor TetR"
                     /note="bacterial operator O2 for the tetR and tetA genes"
     protein_bind    3162..3180
                     /gene="tetO"
                     /label=tet operator
                     /bound_moiety="tetracycline repressor TetR"
                     /note="bacterial operator O2 for the tetR and tetA genes"
     protein_bind    3198..3216
                     /gene="tetO"
                     /label=tet operator
```

```
                     /bound_moiety="tetracycline repressor TetR"
                     /note="bacterial operator O2 for the tetR and tetA genes"
     protein_bind    3311..3335
                     /gene="mutant version of attB"
                     /label=attB1
                     /bound_moiety="BP Clonase(TM)"
                     /note="recombination site for the Gateway(R) BP reaction"
     regulatory      3339..3348
                     /regulatory_class="other"
                     /note="vertebrate consensus sequence for strong initiation
                     of translation (Kozak, 1987)"
     CDS             3341..4708
                     /codon_start=1
                     /label=CMG2delABD-V5
                     /translation="MVAERSPARSPGSWLFPGLWLLVLSGPGGLLRAQEQPSCRRAFDL
                     YFVLDKSGSVANNWIEIYNFVQQLAERFVSPEMRLSFIVFSSQATIILPLTGDRGKISK
                     GLEDLKRVSPVGETYIHEGLKLANEQIQKAGGLKTSSIIIALTDGKLDGLVPSYAEKEA
                     KISRSLGASVYCVGVLDFEQAQLERIADSKEQVFPVKGGFQALKGIINSILAQSCTEIL
                     ELQPSSVCVGEEFQIVLSGRGFMLGSRNGSVLCTYTVNETYTTSVKPVSVQLNSMLCPA
                     PILNKAGETLDVSVSFNGGKSVISGSLIVTATECSNGIAAIIVILVLLLLLGIGLMWWF
                     WPLCCKVVIKDPPPPPAPAPKEEEEEPEETEEPIRPRPPRPKPTHQPPQTKWYTPIKGR
                     LDALWALLRRQYDRVSLMRPQEGDEGRCINFSRVPSQGGGGSGKPIPNPLLGLDST"
     CDS             4664..4705
                     /codon_start=1
                     /product="epitope tag from simian virus 5"
                     /label=V5 tag
                     /translation="GKPIPNPLLGLDST"
     protein_bind    complement(4710..4734)
                     /gene="mutant version of attB"
                     /label=attB2
                     /bound_moiety="BP Clonase(TM)"
                     /note="recombination site for the Gateway(R) BP reaction"
     CDS             4752..4769
                     /codon_start=1
                     /product="6xHis affinity tag"
                     /label=6xHis
                     /translation="HHHHHH"
     promoter        4779..5289
                     /label=hPGK promoter
                     /note="human phosphoglycerate kinase 1 promoter"
     CDS             5299..5895
                     /codon_start=1
                     /gene="pac from Streptomyces alboniger"
                     /product="puromycin N-acetyltransferase"
                     /label=PuroR
                     /note="confers resistance to puromycin"
                     /translation="MTEYKPTVRLATRDDVPRAVRTLAAAFADYPATRHTVDPDRHIER
                     VTELQELFLTRVGLDIGKVWVADDGAAVAVWTTPESVEAGAVFAEIGPRMAELSGSRLA
                     AQQQMEGLLAPHRPKEPAWFLATVGVSPDHQGKGLGSAVVLPGVEAAERAGVPAFLETS
                     APRNLPFYERLGFTVTADVEVPEGPRTWCMTRKPGA"
     CDS             5896..5949
                     /codon_start=1
                     /product="2A peptide from Thosea asigna virus capsid
                     protein"
                     /label=T2A
                     /note="Eukaryotic ribosomes fail to insert a peptide bond
                     between the Gly and Pro residues, yielding separate
                     polypeptides."
                     /translation="EGRGSLLTCGDVEENPGP"
     CDS             5950..6696
                     /codon_start=1
                     /product="improved tetracycline-controlled transactivator"
                     /label=rtTA-Advanced
```

```
                     /note="In the Tet-On(R) system, rtTA-Advanced binds to the
                     Tet-responsive element and stimulates transcription only in
                     the presence of tetracycline or doxycycline."
                     /translation="MSRLDKSKVINGALELLNGVGIEGLTTRKLAQKLGVEQPTLYWHV
                     KNKRALLDALPIEMLDRHHTHFCPLEGESWQDFLRNNAKSYRCALLSHRDGAKVHLGTR
                     PTEKQYETLENQLAFLCQQGFSLENALYALSAVGHFTLGCVLEEQEHQVAKEERETPTT
                     DSMPPLLRQAIELFDRQGAEPAFLFGLELIICGLEKQLKCESGGPTDALDDFDLDMLPA
                     DALDDFDLDMLPADALDDFDLDMLPG"
     CDS             complement(7154..7165)
                     /codon_start=1
                     /product="Factor Xa recognition and cleavage site"
                     /label=Factor Xa site
                     /translation="IEGR"
     LTR             7370..7603
                     /label=3' LTR (Delta-U3)
                     /note="self-inactivating 3' long terminal repeat (LTR) from
                     HIV-1"
     polyA_signal    7675..7796
                     /label=SV40 poly(A) signal
                     /note="SV40 polyadenylation signal"
     rep_origin      7836..7971
                     /label=SV40 ori
                     /note="SV40 origin of replication"
     promoter        7971..8042
                     /gene="bla"
                     /label=AmpR promoter
     CDS             8043..8903
                     /codon_start=1
                     /gene="bla"
                     /product="beta-lactamase"
                     /label=AmpR
                     /note="confers resistance to ampicillin, carbenicillin, and
                     related antibiotics"
                     /translation="MSIQHFRVALIPFFAAFCLPVFAHPETLVKVKDAEDQLGARVGYI
                     ELDLNSGKILESFRPEERFPMMSTFKVLLCGAVLSRIDAGQEQLGRRIHYSQNDLVEYS
                     PVTEKHLTDGMTVRELCSAAITMSDNTAANLLLTTIGGPKELTAFLHNMGDHVTRLDRW
                     EPELNEAIPNDERDTTMPVAMATTLRKLLTGELLTLASRQQLIDWMEADKVAGPLLRSA
                     LPAGWFIADKSGAGERGSRGIIAALGPDGKPSRIVVIYTTGSQATMDERNRQIAEIGAS
                     LIKHW"
ORIGIN
        1 ttgagatcct ttttttctgc gcgtaatctg ctgcttgcaa acaaaaaaac caccgctacc
       61 agcggtggtt tgtttgccgg atcaagagct accaactctt tttccgaagg taactggctt
      121 cagcagagcg cagataccaa atactgttct tctagtgtag ccgtagttag gccaccactt
      181 caagaactct gtagcaccgc ctacatacct cgctctgcta atcctgttac cagtggctgc
      241 tgccagtggc gataagtcgt gtcttaccgg gttggactca agacgatagt taccggataa
      301 ggcgcagcgg tcgggctgaa cggggggttc gtgcacacag cccagcttgg agcgaacgac
      361 ctacaccgaa ctgagatacc tacagcgtga gctatgagaa agcgccacgc ttcccgaagg
      421 gagaaaggcg gacaggtatc cggtaagcgg cagggtcgga acaggagagc gcacgaggga
      481 gcttccaggg ggaaacgcct ggtatcttta tagtcctgtc gggtttcgcc acctctgact
      541 tgagcgtcga tttttgtgat gctcgtcagg ggggcggagc ctatggaaaa acgccagcaa
      601 cgcggccttt ttacggttcc tggccttttg ctggcctttt gctcacatgt tctttcctgc
      661 gttatcccct gattctgtgg ataaccgtat taccgccttt gagtgagctg ataccgctcg
      721 ccgcagccga acgaccgagc gcagcgagtc agtgagcgag gaagcggaag agcgcccaat
      781 acgcaaaccg cctctccccg cgcgttggcc gattcattaa tgcagctggc acgacaggtt
      841 tcccgactgg aaagcgggca gtgagcgcaa cgcaattaat gtgagttagc tcactcatta
      901 ggcaccccag gctttacact ttatgcttcc ggctcgtatg ttgtgtggaa ttgtgagcgg
      961 ataacaattt cacacaggaa acagctatga ccatgattac gccaagcgcg caattaaccc
     1021 tcactaaagg gaacaaaagc tggagctgca agcttaatgt agtcttatgc aatactcttg
     1081 tagtcttgca acatggtaac gatgagttag caacatgcct tacaaggaga gaaaaagcac
     1141 cgtgcatgcc gattggtgga agtaaggtgg tacgatcgtg ccttattagg aaggcaacag
     1201 acgggtctga catggattgg acgaaccact gaattgccgc attgcagaga tattgtattt
     1261 aagtgcctag ctcgatacat aaacgggtct ctctggttag accagatctg agcctgggag
     1321 ctctctggct aactagggaa cccactgctt aagcctcaat aaagcttgcc ttgagtgctt
```

1381 caagtagtgt gtgcccgtct gttgtgtgac tctggtaact agagatccct cagacccttt
1441 tagtcagtgt ggaaaatctc tagcagtggc gcccgaacag ggacttgaaa gcgaaaggga
1501 aaccagagga gctctctcga cgcaggactc ggcttgctga agcgcgcacg gcaagaggcg
1561 aggggcggcg actggtgagt acgccaaaaa ttttgactag cggaggctag aaggagagag
1621 atgggtgcga gagcgtcagt attaagcggg ggagaattag atcgcgatgg gaaaaaattc
1681 ggttaaggcc agggggaaag aaaaaatata aattaaaaca tatagtatgg gcaagcaggg
1741 agctagaacg attcgcagtt aatcctggcc tgttagaaac atcagaaggc tgtagacaaa
1801 tactgggaca gctacaacca tcccttcaga caggatcaga agaacttaga tcattatata
1861 atacagtagc aaccctctat tgtgtgcatc aaaggataga gataaaagac accaaggaag
1921 ctttagacaa gatagaggaa gagcaaaaca aaagtaagac caccgcacag caagcggccg
1981 ctgatcttca gacctggagg aggagatatg agggacaatt ggagaagtga attatataaa
2041 tataaagtag taaaaattga accattagga gtagcaccca ccaaggcaaa gagaagagtg
2101 gtgcagagag aaaaaagagc agtgggaata ggagctttgt tccttgggtt cttgggagca
2161 gcaggaagca ctatgggcgc agcgtcaatg acgctgacgg tacaggccag acaattattg
2221 tctggtatag tgcagcagca gaacaatttg ctgagggcta ttgaggcgca acagcatctg
2281 ttgcaactca cagtctgggg catcaagcag ctccaggcaa gaatcctggc tgtggaaaga
2341 tacctaaagg atcaacagct cctggggatt tggggttgct ctggaaaact catttgcacc
2401 actgctgtgc cttggaatgc tagttggagt aataaatctc tggaacagat ttggaatcac
2461 acgacctgga tggagtggga cagagaaatt aacaattaca caagcttaat acactcctta
2521 attgaagaat cgcaaaacca gcaagaaaag aatgaacaag aattattgga attagataaa
2581 tgggcaagtt tgtggaattg gtttaacata acaaattggc tgtggtatat aaaattattc
2641 ataatgatag taggaggctt ggtaggttta agaatagttt ttgctgtact ttctatagtg
2701 aatagagtta ggcagggata ttcaccatta tcgtttcaga cccacctccc aaccccgagg
2761 ggacaattct cgacctcgag acaaatggca gtattcatcc acaattttaa aagaaaaggg
2821 gggattgggg ggtacagtgc aggggaaaga atagtagaca taatagcaac agacatacaa
2881 actaaagaat tacaaaaaca aattacaaaa attcaaaatt ttcgggttta ttacagggac
2941 agcagagatc cactttggcc gcgaatcgat atgtcgagtt tactccctat cagtgataga
3001 gaacgtatgt cgagtttact ccctatcagt gatagagaac gatgtcgagt ttactcccta
3061 tcagtgatag agaacgtatg tcgagtttac tccctatcag tgatagagaa cgtatgtcga
3121 gtttactccc tatcagtgat agagaacgta tgtcgagttt atccctatca gtgatagaga
3181 acgtatgtcg agtttactcc ctatcagtga tagagaacgt atgtcgaggt aggcgtgtac
3241 ggtgggaggc ctatataagc agagctcgtt tagtgaaccg tcagatcgcc tggagaattg
3301 gctagcatca acaagtttgt acaaaaaagc aggctgcacc atggtggcgg aacggtcccc
3361 cgctcgcagt cccggtagct ggcttttttcc cgggttatgg cttttggtgt tgtctgggcc
3421 ggggggactc ctccgggctc aggaacagcc ctcttgtaga cgagcatttg atctgtattt
3481 cgtgctggac aaatctggct ccgtcgctaa taattggatt gagatatata actttgtgca
3541 gcagctggct gaacgcttcg tttcaccgga aatgagactg tccttcatcg tgtttagcag
3601 ccaggctact attatcctcc ctctcactgg agaccgggga aagatctcca agggtctcga
3661 agacctcaag cgggtgtccc cggtgggaga aacatacatc catgagggcc tcaaattggc
3721 caacgaacag atacagaaag cgggggggcct gaaaacctcc tccataatca ttgccctcac
3781 agatggaaaa ctggatgggc tcgttccctc ctatgccgag aaagaggcaa aaatcagtag
3841 atccctcggc gcctccgtct attgcgtggg ggtgctggac ttcgaacaag cgcagctgga
3901 gcgcattgcc gacagcaagg aacaagtgtt ccctgtgaaa ggaggatttc aagctctcaa
3961 agggatcata aattccatct tggcgcagtc ttgtactgaa atccttgaat tacaaccatc
4021 ttccgtttgt gtcggtgagg agttccagat cgtgttgagt gggaggggct tcatgctggg
4081 gagccgaaat ggcagcgttc tctgcaccta taccgtgaac gaaacatata ctactagtgt
4141 aaaaccagtg agcgtgcagc tgaacagcat gctgtgtcct gcccccatcc tgaataaagc
4201 gggggagact ctcgatgtta gcgtcagttt taatgggggg aagagcgtga ttagtggaag
4261 tctcattgtt accgctacag aatgtagcaa cggtatagcc gctatcattg ttatcctggt
4321 tctcctgttg cttctgggga taggactcat gtggtggttc tggccgctgt gttgtaaggt
4381 ggtaattaag gacccacctc caccacccgc acccgctccg aaagaggagg aggaagaacc
4441 ggaagagaca gaggagccta ttaggcctag accccccaga ccaaagccca cacatcaacc
4501 gccccaaaca aagtggtata ccccaattaa gggtagactg gatgcattat gggccctgct
4561 gcgccgccag tacgacagag tgtctttaat gcggccacag gagggcgatg aggggagatg
4621 tataaatttt tcacgagtcc catctcaggg tggaggtggc tccggcaagc caatccctaa
4681 ccctctgttg ggactggata gcacatagga cccagctttc ttgtacaaag tggtttagta
4741 atgaaccggt ccaccaccac caccaccact aaggatccgg ggttggggtt gcgccttttc
4801 caaggcagcc ctgggtttgc gcagggacgc ggctgctctg ggcgtggttc cgggaaacgc
4861 agcggcgccg accctgggtc tcgcacattc ttcacgtccg ttcgcagcgt cacccggatc
4921 ttcgccgcta cccttgtggg ccccccggcg acgcttcctg ctccgcccct aagtcgggaa
4981 ggttccttgc ggttcgcggc gtgccggacg tgacaaacgg aagccgcacg tctcactagt
5041 accctcgcag acggacagcg ccagggagca atggcagcgc gccgaccgcg atgggctgtg
5101 gccaatagcg gctgctcagc agggcgcgcc gagagcagcg gccgggaagg ggcggtgcgg

5161 gaggcggggt gtggggcggt agtgtgggcc ctgttcctgc ccgcgcggtg ttccgcattc
5221 tgcaagcctc cggagcgcac gtcggcagtc ggctccctcg ttgaccgaat caccgacctc
5281 tctccccagc aattcaccat gaccgagtac aagcccacgg tgcgcctcgc cacccgcgac
5341 gacgtcccca gggccgtacg caccctcgcc gccgcgttcg ccgactaccc cgccacgcgc
5401 cacaccgtcg atccggaccg ccacatcgag cgggtcaccg agctgcaaga actcttcctc
5461 acgcgcgtcg ggctcgacat cggcaaggtg tgggtcgcgg acgacggcgc cgcggtggcg
5521 gtctggacca cgccggagag cgtcgaagcg gggggcggtgt tcgccgagat cggcccgcgc
5581 atggccgagt tgagcggttc ccggctggcc gcgcagcaac agatggaagg cctcctggcg
5641 ccgcaccggc ccaaggagcc cgcgtggttc ctggccaccg tcggcgtctc gcccgaccac
5701 cagggcaagg gtctgggcag cgccgtcgtg ctccccggag tggaggcggc cgagcgcgcc
5761 ggggtgcccg ccttcctgga gacctccgcg ccccgcaacc tccccttcta cgagcggctc
5821 ggcttcaccg tcaccgccga cgtcgaggtg cccgaaggac cgcgcacctg gtgcatgacc
5881 cgcaagcccg gtgccgaagg tagaggttct ctcctcactt gtggtgatgt tgaagaaaac
5941 cctggtccaa tgtctagact ggacaagagc aaagtcataa acggagctct ggaattactc
6001 aatggtgtcg gtatcgaagg cctgacgaca aggaaactcg ctcaaaagct gggagttgag
6061 cagcctaccc tgtactggca cgtgaagaac aagcgggccc tgctcgatgc cctgccaatc
6121 gagatgctgg acaggcatca tacccacttc tgcccctgg aaggcgagtc atggcaagac
6181 tttctgcgga acaacgccaa gtcataccgc tgtgctctcc tctcacatcg cgacggggct
6241 aaagtgcatc tcggcacccg cccaacagag aaacagtacg aaaccctgga aaatcagctc
6301 gcgttcctgt gtcagcaagg cttctccctg gagaacgcac tgtacgctct gtccgccgtg
6361 ggccacttta cactgggctg cgtattggag gaacaggagc atcaagtagc aaaagaggaa
6421 agagagacac ctaccaccga ttctatgccc ccacttctga gacaagcaat tgagctgttc
6481 gaccggcagg gagccgaacc tgccttcctt ttcggcctgg aactaatcat atgtggcctg
6541 gagaaacagc taaagtgcga aagcggcggg ccgaccgacg cccttgacga ttttgactta
6601 gacatgctcc cagccgatgc ccttgacgac tttgaccttg atatgctgcc tgctgacgct
6661 cttgacgatt ttgaccttga catgctcccc gggtaactaa gtaaggatcg atccaagata
6721 tcgtattctt aactatgttg ctccttttac gctatgtgga tacgctgctt taatgccttt
6781 gtatcatgct attgcttccc gtatggcttt cattttctcc tccttgtata aatcctggtt
6841 gctgtctctt tatgaggagt tgtggcccgt tgtcaggcaa cgtggcgtgg tgtgcactgt
6901 gtttgctgac gcaaccccca ctggttgggg cattgccacc acctgtcagc tcctttccgg
6961 gactttcgct ttccccctcc ctattgccac ggcggaactc atcgccgcct gccttgcccg
7021 ctgctggaca ggggctcggc tgttgggcac tgacaattcc gtggtgttgt cggggaagct
7081 gacgtccttt ccatggctgc tcgcctgtgt tgccacctgg attctgcgcg ggacgtcctt
7141 ctgctacgtc ccttcggccc tcaatccagc ggaccttcct tcccgcggcc tgctgccggc
7201 tctgcggcct cttccgcgtc ttcgccttcg ccctcagacg agtcggatct ccctttgggc
7261 cgcctccccg cctgtttcgc ctcggcgtcc ggactagagg tacctttaag accaatgact
7321 tacaaggcag ctgtagatct tagccacttt ttaaaagaaa aggggggact ggaagggcta
7381 attcactccc aacgaagaca agatctgctt tttgcttgta ctgggtctct ctggttagac
7441 cagatctgag cctgggagct ctctggctaa ctagggaacc cactgcttaa gcctcaataa
7501 agcttgcctt gagtgcttca agtagtgtgt gcccgtctgt tgtgtgactc tggtaactag
7561 agatccctca gaccctttta gtcagtgtgg aaaatctcta gcagtagtag ttcatgtcat
7621 cttattattc agtatttata acttgcaaag aaatgaatat cagagagtga gaggaacttg
7681 tttattgcag cttataatgg ttacaaataa agcaatagca tcacaaattt cacaaataaa
7741 gcattttttt cactgcattc tagttgtggt ttgtccaaac tcatcaatgt atcttatcat
7801 gtctggctct agctatcccg cccctaactc cgcccatccc gcccctaact ccgcccagtt
7861 ccgcccattc tccgccccat ggctgactaa ttttttttat ttatgcagag gccgaggccg
7921 cctcggcctc tgagctattc cagaagtagt gaggaggctt ttttggaggc ctttcaaata
7981 tgtatccgct catgagacaa taaccctgat aaatgcttca ataatattga aaaaggaaga
8041 gtatgagtat tcaacatttc cgtgtcgccc ttattccctt ttttgcggca ttttgccttc
8101 ctgtttttgc tcacccagaa acgctggtga aagtaaaaga tgctgaagat cagttgggtg
8161 cacgagtggg ttacatcgaa ctggatctca acagcggtaa gatccttgag agttttcgcc
8221 ccgaagaacg ttttccaatg atgagcactt ttaaagttct gctatgtggc gcggtattat
8281 cccgtattga cgccgggcaa gagcaactcg gtcgccgcat acactattct cagaatgact
8341 tggttgagta ctcaccagtc acagaaaagc atcttacgga tggcatgaca gtaagagaat
8401 tatgcagtgc tgccataacc atgagtgata acactgcggc caacttactt ctgacaacga
8461 tcggaggacc gaaggagcta accgcttttt tgcacaacat gggggatcat gtaactcgcc
8521 ttgatcgttg ggaaccggag ctgaatgaag ccataccaaa cgacgagcgt gacaccacga
8581 tgcctgtagc aatggcaaca acgttgcgca aactattaac tggcgaacta cttactctag
8641 cttcccggca acaattaata gactggatgg aggcggataa agttgcagga ccacttctgc
8701 gctcggccct tccggctggc tggtttattg ctgataaatc tggagccggt gagcgtgggt
8761 ctcgcggtat cattgcagca ctggggccag atggtaagcc ctcccgtatc gtagttatct
8821 acacgacggg gagtcaggca actatggatg aacgaaatag acagatcgct gagataggtg
8881 cctcactgat taagcattgg taactgtcag accaagttta ctcatatata ctttagattg

```
     8941 atttaaaact tcatttttaa tttaaaagga tctaggtgaa gatccttttt gataatctca
     9001 tgaccaaaat cccttaacgt gagttttcgt tccactgagc gtcagacccc gtagaaaaga
     9061 tcaaaggatc ttc
//
```

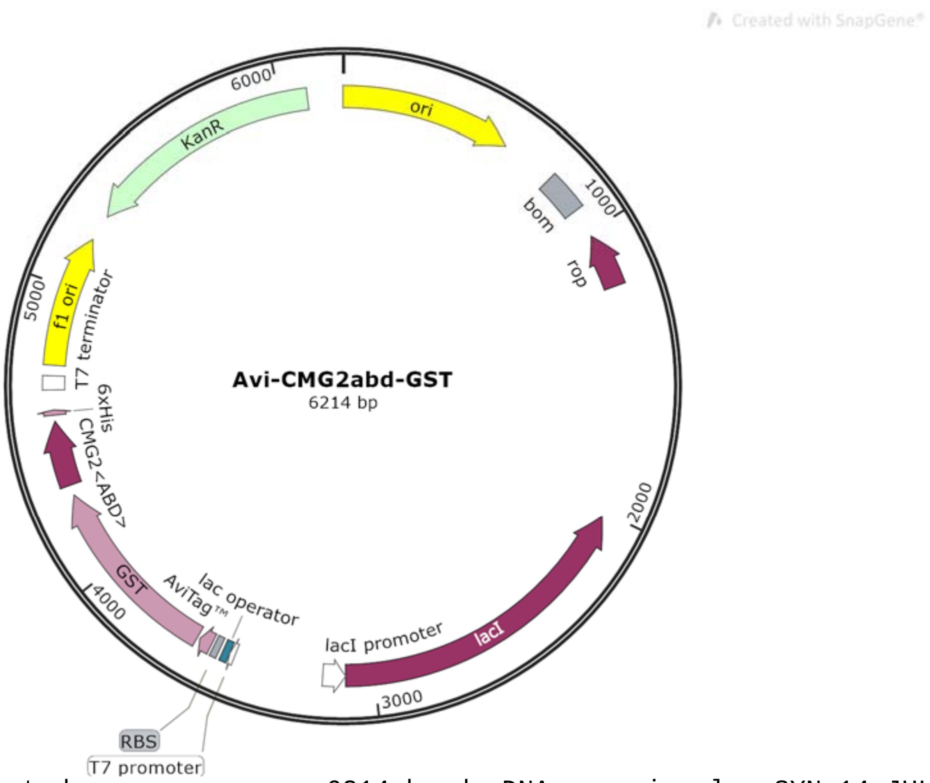


```
LOCUS       Exported                6214 bp ds-DNA     circular SYN 14-JUL-2025
DEFINITION  synthetic circular DNA
ACCESSION   .
VERSION     .
KEYWORDS    .
SOURCE      synthetic DNA construct
  ORGANISM  synthetic DNA construct
REFERENCE   1  (bases 1 to 6214)
  AUTHORS   Vascular Biology Program
  TITLE     Direct Submission
  JOURNAL   Exported Jul 14, 2025 from SnapGene 4.2.11
            http://www.snapgene.com
FEATURES             Location/Qualifiers
     source          1..6214
                     /organism="synthetic DNA construct"
                     /mol_type="other DNA"
     rep_origin      1..589
                     /direction=RIGHT
                     /label=ori
                     /note="high-copy-number ColE1/pMB1/pBR322/pUC origin of
                     replication"
     misc_feature    775..914
                     /label=bom
                     /note="basis of mobility region from pBR322"
     CDS             complement(1016..1207)
                     /codon_start=1
                     /gene="rop"
                     /product="Rop protein, which maintains plasmids at low copy
                     number"
                     /label=rop
                     /translation="MTKQEKTALNMARFIRSQTLTLLEKLNELDADEQADICESLHDHA
                     DELYRSCLARFGDDGENL"
     CDS             complement(2016..3098)
                     /codon_start=1
                     /gene="lacI"
                     /product="lac repressor"
                     /label=lacI
```

```
                     /note="The lac repressor binds to the lac operator to
                     inhibit transcription in E. coli. This inhibition can be
                     relieved by adding lactose or
                     isopropyl-beta-D-thiogalactopyranoside (IPTG)."
                     /translation="MKPVTLYDVAEYAGVSYQTVSRVVNQASHVSAKTREKVEAAMAEL
                     NYIPNRVAQQLAGKQSLLIGVATSSLALHAPSQIVAAIKSRADQLGASVVVSMVERSGV
                     EACKAAVHNLLAQRVSGLIINYPLDDQDAIAVEAACTNVPALFLDVSDQTPINSIIFSH
                     EDGTRLGVEHLVALGHQQIALLAGPLSSVSARLRLAGWHKYLTRNQIQPIAEREGDWSA
                     MSGFQQTMQMLNEGIVPTAMLVANDQMALGAMRAITESGLRVGADISVVGYDDTEDSSC
                     YIPPLTTIKQDFRLLGQTSVDRLLQLSQGQAVKGNQLLPVSLVKRKTTLAPNTQTASPR
                     ALADSLMQLARQVSRLESGQ"
     promoter        complement(3099..3176)
                     /gene="lacI"
                     /label=lacI promoter
     promoter        3485..3503
                     /label=T7 promoter
                     /note="promoter for bacteriophage T7 RNA polymerase"
     protein_bind    3504..3528
                     /label=lac operator
                     /bound_moiety="lac repressor encoded by lacI"
                     /note="The lac repressor binds to the lac operator to
                     inhibit transcription in E. coli. This inhibition can be
                     relieved by adding lactose or
                     isopropyl-beta-D-thiogalactopyranoside (IPTG)."
     RBS             3541..3563
                     /note="efficient ribosome binding site from bacteriophage
                     T7 gene 10 (Olins and Rangwala, 1989)"
     CDS             3573..3617
                     /codon_start=1
                     /product="peptide tag that allows for enzymatic
                     biotinylation"
                     /label=AviTag(TM)
                     /translation="GLNDIFEAQKIEWHE"
     CDS             3627..4280
                     /codon_start=1
                     /product="glutathione S-transferase from Schistosoma
                     japonicum"
                     /label=GST
                     /translation="MSPILGYWKIKGLVQPTRLLLEYLEEKYEEHLYERDEGDKWRNKK
                     FELGLEFPNLPYYIDGDVKLTQSMAIIRYIADKHNMLGGSPKERAEISMLEGAVLDIRY
                     GVSRIAYSKDFETLKVDFLSKLPEMLKMFEDRLSHKTYLNGDHVTHPDFMLYDALDVVL
                     YMDPMSLDAFPKLVSFKKRIEAIPQIDKYLKSSKYIAWPLQGWQATFGGGDHPPK"
     CDS             4317..4541
                     /codon_start=1
                     /label=CMG2<ABD>
                     /translation="KVVIKDPPPPPAPAPKEEEEEPLPTKKWPTVDASYYGGRGVGGIK
                     RMEVRWGDKGSTEEGARLEKAKNAVVKIP"
     CDS             4563..4580
                     /codon_start=1
                     /product="6xHis affinity tag"
                     /label=6xHis
                     /translation="HHHHHH"
     terminator      4647..4694
                     /label=T7 terminator
                     /note="transcription terminator for bacteriophage T7 RNA
                     polymerase"
     rep_origin      4731..5186
                     /direction=RIGHT
                     /label=f1 ori
                     /note="f1 bacteriophage origin of replication; arrow
                     indicates direction of (+) strand synthesis"
     CDS             complement(5278..6093)
                     /codon_start=1
```

```
                     /gene="aph(3')-Ia"
                     /product="aminoglycoside phosphotransferase"
                     /label=KanR
                     /note="confers resistance to kanamycin in bacteria or G418
                     (Geneticin(R)) in eukaryotes"
                     /translation="MSHIQRETSCSRPRLNSNMDADLYGYKWARDNVGQSGATIYRLYG
                     KPDAPELFLKHGKGSVANDVTDEMVRLNWLTEFMPLPTIKHFIRTPDDAWLLTTAIPGK
                     TAFQVLEEYPDSGENIVDALAVFLRRLHSIPVCNCPFNSDRVFRLAQAQSRMNNGLVDA
                     SDFDDERNGWPVEQVWKEMHKLLPFSPDSVVTHGDFSLDNLIFDEGKLIGCIDVGRVGI
                     ADRYQDLAILWNCLGEFSPSLQKRLFQKYGIDNPDMNKLQFHLMLDEFF"
ORIGIN
        1 ttgagatcct ttttttctgc gcgtaatctg ctgcttgcaa acaaaaaaac caccgctacc
       61 agcggtggtt tgtttgccgg atcaagagct accaactctt tttccgaagg taactggctt
      121 cagcagagcg cagataccaa atactgtcct tctagtgtag ccgtagttag gccaccactt
      181 caagaactct gtagcaccgc ctacatacct cgctctgcta atcctgttac cagtggctgc
      241 tgccagtggc gataagtcgt gtcttaccgg gttggactca agacgatagt taccggataa
      301 ggcgcagcgg tcgggctgaa cggggggttc gtgcacacag cccagcttgg agcgaacgac
      361 ctacaccgaa ctgagatacc tacagcgtga gctatgagaa agcgccacgc ttcccgaagg
      421 gagaaaggcg gacaggtatc cggtaagcgg cagggtcgga acaggagagc gcacgaggga
      481 gcttccaggg ggaaacgcct ggtatcttta tagtcctgtc gggtttcgcc acctctgact
      541 tgagcgtcga tttttgtgat gctcgtcagg ggggcggagc ctatggaaaa acgccagcaa
      601 cgcggccttt ttacggttcc tggccttttg ctggcctttt gctcacatgt tctttcctgc
      661 gttatcccct gattctgtgg ataaccgtat taccgccttt gagtgagctg ataccgctcg
      721 ccgcagccga acgaccgagc gcagcgagtc agtgagcgag gaagcggaag agcgcctgat
      781 gcggtatttt ctccttacgc atctgtgcgg tatttcacac cgcaatggtg cactctcagt
      841 acaatctgct ctgatgccgc atagttaagc cagtatacac tccgctatcg ctacgtgact
      901 gggtcatggc tgcgccccga cacccgccaa cacccgctga cgcgccctga cgggcttgtc
      961 tgctcccggc atccgcttac agacaagctg tgaccgtctc cgggagctgc atgtgtcaga
     1021 ggttttcacc gtcatcaccg aaacgcgcga ggcagctgcg gtaaagctca tcagcgtggt
     1081 cgtgaagcga ttcacagatg tctgcctgtt catccgcgtc cagctcgttg agtttctcca
     1141 gaagcgttaa tgtctggctt ctgataaagc gggccatgtt aagggcggtt ttttcctgtt
     1201 tggtcactga tgcctccgtg taagggggat ttctgttcat gggggtaatg ataccgatga
     1261 aacgagagag gatgctcacg atacgggtta ctgatgatga acatgcccgg ttactggaac
     1321 gttgtgaggg taaacaactg gcggtatgga tgcggcggga ccagagaaaa atcactcagg
     1381 gtcaatgcca gcgcttcgtt aatacagatg taggtgttcc acagggtagc cagcagcatc
     1441 ctgcgatgca gatccggaac ataatggtgc agggcgctga cttccgcgtt tccagacttt
     1501 acgaaacacg gaaaccgaag accattcatg ttgttgctca ggtcgcagac gttttgcagc
     1561 agcagtcgct tcacgttcgc tcgcgtatcg gtgattcatt ctgctaacca gtaaggcaac
     1621 cccgccagcc tagccgggtc ctcaacgaca ggagcacgat catgcgcacc cgtggggccg
     1681 ccatgccggc gataatggcc tgcttctcgc cgaaacgttt ggtggcggga ccagtgacga
     1741 aggcttgagc gagggcgtgc aagattccga ataccgcaag cgacaggccg atcatcgtcg
     1801 cgctccagcg aaagcggtcc tcgccgaaaa tgacccagag cgctgccggc acctgtccta
     1861 cgagttgcat gataaagaag acagtcataa gtgcggcgac gatagtcatg ccccgcgccc
     1921 accggaagga gctgactggg ttgaaggctc tcaagggcat cggtcgagat cccggtgcct
     1981 aatgagtgag ctaacttaca ttaattgcgt tgcgctcact gcccgctttc cagtcgggaa
     2041 acctgtcgtg ccagctgcat taatgaatcg gccaacgcgc ggggagaggc ggtttgcgta
     2101 ttgggcgcca gggtggtttt tcttttcacc agtgagacgg gcaacagctg attgcccttc
     2161 accgcctggc cctgagagag ttgcagcaag cggtccacgc tggtttgccc cagcaggcga
     2221 aaatcctgtt tgatggtggt taacggcggg atataacatg agctgtcttc ggtatcgtcg
     2281 tatcccacta ccgagatatc cgcaccaacg cgcagcccgg actcggtaat ggcgcgcatt
     2341 gcgcccagcg ccatctgatc gttggcaacc agcatcgcag tgggaacgat gccctcattc
     2401 agcatttgca tggtttgttg aaaaccggac atggcactcc agtcgccttc ccgttccgct
     2461 atcggctgaa tttgattgcg agtgagatat ttatgccagc cagccagacg cagacgcgcc
     2521 gagacagaac ttaatgggcc cgctaacagc gcgatttgct ggtgacccaa tgcgaccaga
     2581 tgctccacgc ccagtcgcgt accgtcttca tgggagaaaa taatactgtt gatgggtgtc
     2641 tggtcagaga catcaagaaa taacgccgga acattagtgc aggcagcttc cacagcaatg
     2701 gcatcctggt catccagcgg atagttaatg atcagcccac tgacgcgttg cgcgagaaga
     2761 ttgtgcaccg ccgctttaca ggcttcgacg ccgcttcgtt ctaccatcga caccaccacg
     2821 ctggcaccca gttgatcggc gcgagattta atcgccgcga caatttgcga cggcgcgtgc
     2881 agggccagac tggaggtggc aacgccaatc agcaacgact gtttgcccgc cagttgttgt
     2941 gccacgcggt tgggaatgta attcagctcc gccatcgccg cttccacttt ttcccgcgtt
     3001 ttcgcagaaa cgtggctggc ctggttcacc acgcgggaaa cggtctgata agagacaccg
     3061 gcatactctg cgacatcgta taacgttact ggtttcacat tcaccaccct gaattgactc
```

3121 tcttccgggc gctatcatgc cataccgcga aaggttttgc gccattcgat ggtgtccggg
3181 atctcgacgc tctcccttat gcgactcctg cattaggaag cagcccagta gtaggttgag
3241 gccgttgagc accgccgccg caaggaatgg tgcatgcaag gagatggcgc ccaacagtcc
3301 cccggccacg gggcctgcca ccatacccac gccgaaacaa gcgctcatga gcccgaagtg
3361 gcgagcccga tcttccccat cggtgatgtc ggcgatatag gcgccagcaa ccgcacctgt
3421 ggcgccggtg atgccggcca cgatgcgtcc ggcgtagagg atcgagatct cgatcccgcg
3481 aaattaatac gactcactat aggggaattg tgagcggata acaattcccc tctaggatcc
3541 tttgtttaac tttaagaagg agatatacaa tgggcctgaa cgacatcttc gaggctcaga
3601 aaatcgaatg gcacgaacgc ggatccatgt cccctatact aggttattgg aaaattaagg
3661 gccttgtgca acccactcga cttcttttgg aatatcttga agaaaaatat gaagagcatt
3721 tgtatgagcg cgatgaaggt gataaatggc gaaacaaaaa gtttgaattg ggtttggagt
3781 ttcccaatct tccttattat attgatggtg atgttaaatt aacacagtct atggccatca
3841 tacgttatat agctgacaag cacaacatgt tgggtggttc tccaaaagag cgtgcagaga
3901 tttcaatgct tgaaggagcg gttttggata ttagatacgg tgtttcgaga attgcatata
3961 gtaaagactt tgaaactctc aaagttgatt ttcttagcaa gctacctgaa atgctgaaaa
4021 tgttcgaaga tcgtttatct cataaaacat atttaaatgg tgatcatgta acccatcctg
4081 acttcatgtt gtatgacgct cttgatgttg ttttatacat ggacccaatg tccctggatg
4141 cgttcccaaa attagtttct tttaaaaaac gtattgaagc tatcccacaa attgataagt
4201 acttgaaatc cagcaagtat atagcatggc ctttgcaggg ctggcaagcc acgtttggtg
4261 gtggcgacca tcctccaaaa ggtggcggtt ccggtggcgg ttccggtggc ggttccaagg
4321 tggtaattaa ggacccacct ccaccacccg cacccgctcc gaaagaggag gaggaagaac
4381 cgctgcctac taagaaatgg cccaccgtgg acgcttcata ctatggcggg cgtggcgtgg
4441 gcggcattaa aggatggag gtaaggtggg gagataaggg atcaactgaa gagggtgcac
4501 gactggagaa ggctaagaat gctgtcgtaa agattcctta gaagcttgcg gccgcactcg
4561 agcaccacca ccaccaccac tgagatccgg ctgctaacaa agcccgaaag gaagctgagt
4621 tggctgctgc caccgctgag caataactag cataacccct tggggcctct aaacgggtct
4681 tgaggggttt tttgctgaaa ggaggaacta tatccggatt ggcgaatggg acgcgccctg
4741 tagcggcgca ttaagcgcgg cgggtgtggt ggttacgcgc agcgtgaccg ctacacttgc
4801 cagcgcccta gcgcccgctc ctttcgcttt cttcccttcc tttctcgcca cgttcgccgg
4861 ctttccccgt caagctctaa atcgggggct ccctttaggg ttccgattta gtgctttacg
4921 gcacctcgac cccaaaaaac ttgattaggg tgatggttca cgtagtgggc catcgccctg
4981 atagacggtt tttcgccctt tgacgttgga gtccacgttc tttaatagtg gactcttgtt
5041 ccaaactgga acaacactca accctatctc ggtctattct tttgatttat aagggatttt
5101 gccgatttcg gcctattggt taaaaaatga gctgatttaa caaaaattta acgcgaattt
5161 taacaaaata ttaacgctta caatttaggt ggcacttttc ggggaaatgt gcgcggaacc
5221 cctatttgtt tatttttcta aatacattca aatatgtatc cgctcatgaa ttaattctta
5281 gaaaaactca tcgagcatca aatgaaactg caatttattc atatcaggat tatcaatacc
5341 atatttttga aaaagccgtt tctgtaatga aggagaaaac tcaccgaggc agttccatag
5401 gatggcaaga tcctggtatc ggtctgcgat tccgactcgt ccaacatcaa tacaacctat
5461 taatttcccc tcgtcaaaaa taaggttatc aagtgagaaa tcaccatgag tgacgactga
5521 atccggtgag aatggcaaaa gtttatgcat ttctttccag acttgttcaa caggccagcc
5581 attacgctcg tcatcaaaat cactcgcatc aaccaaaccg ttattcattc gtgattgcgc
5641 ctgagcgaga cgaaatacgc gatcgctgtt aaaaggacaa ttacaaacag gaatcgaatg
5701 caaccggcgc aggaacactg ccagcgcatc aacaatattt tcacctgaat caggatattc
5761 ttctaatacc tggaatgctg ttttcccggg gatcgcagtg gtgagtaacc atgcatcatc
5821 aggagtacgg ataaaatgct tgatggtcgg aagaggcata aattccgtca gccagtttag
5881 tctgaccatc tcatctgtaa catcattggc aacgctacct ttgccatgtt tcagaaacaa
5941 ctctggcgca tcgggcttcc catacaatcg atagattgtc gcacctgatt gcccgacatt
6001 atcgcgagcc catttatacc catataaatc agcatccatg ttggaattta atcgcggcct
6061 agagcaagac gtttcccgtt gaatatggct cataacaccc cttgtattac tgtttatgta
6121 agcagacagt tttattgttc atgaccaaaa tcccttaacg tgagttttcg ttccactgag
6181 cgtcagaccc cgtagaaaag atcaaaggat cttc
//

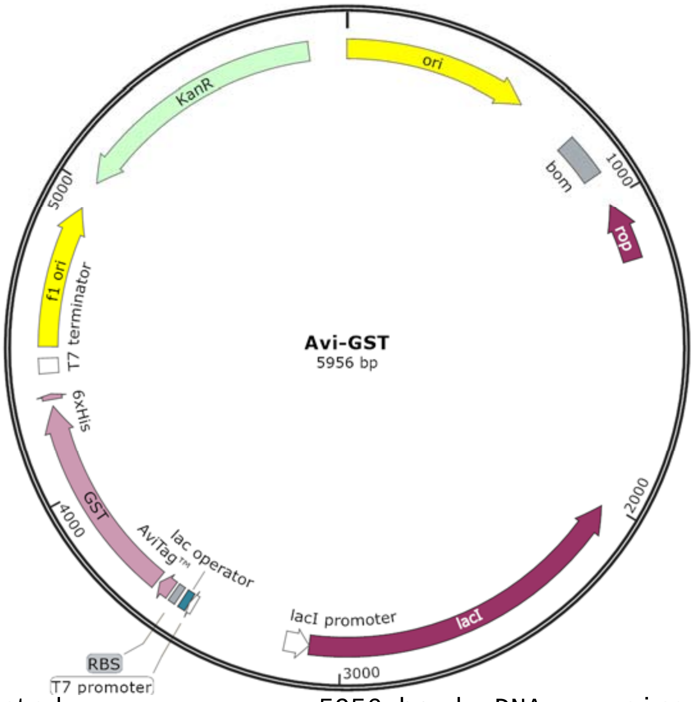


```
LOCUS       Exported                5956 bp ds-DNA     circular SYN 14-JUL-2025
DEFINITION  synthetic circular DNA
ACCESSION   .
VERSION     .
KEYWORDS    .
SOURCE      synthetic DNA construct
  ORGANISM  synthetic DNA construct
REFERENCE   1  (bases 1 to 5956)
  AUTHORS   Vascular Biology Program
  TITLE     Direct Submission
  JOURNAL   Exported Jul 14, 2025 from SnapGene 4.2.11
            http://www.snapgene.com
FEATURES             Location/Qualifiers
     source          1..5956
                     /organism="synthetic DNA construct"
                     /mol_type="other DNA"
     rep_origin      1..589
                     /direction=RIGHT
                     /label=ori
                     /note="high-copy-number ColE1/pMB1/pBR322/pUC origin of
                     replication"
     misc_feature    775..914
                     /label=bom
                     /note="basis of mobility region from pBR322"
     CDS             complement(1016..1207)
                     /codon_start=1
                     /gene="rop"
                     /product="Rop protein, which maintains plasmids at low copy
                     number"
                     /label=rop
                     /translation="MTKQEKTALNMARFIRSQTLTLLEKLNELDADEQADICESLHDHA
                     DELYRSCLARFGDDGENL"
     CDS             complement(2016..3098)
                     /codon_start=1
                     /gene="lacI"
                     /product="lac repressor"
                     /label=lacI
                     /note="The lac repressor binds to the lac operator to
                     inhibit transcription in E. coli. This inhibition can be
```

```
                     relieved by adding lactose or
                     isopropyl-beta-D-thiogalactopyranoside (IPTG)."
                     /translation="MKPVTLYDVAEYAGVSYQTVSRVVNQASHVSAKTREKVEAAMAEL
                     NYIPNRVAQQLAGKQSLLIGVATSSLALHAPSQIVAAIKSRADQLGASVVVSMVERSGV
                     EACKAAVHNLLAQRVSGLIINYPLDDQDAIAVEAACTNVPALFLDVSDQTPINSIIFSH
                     EDGTRLGVEHLVALGHQQIALLAGPLSSVSARLRLAGWHKYLTRNQIQPIAEREGDWSA
                     MSGFQQTMQMLNEGIVPTAMLVANDQMALGAMRAITESGLRVGADISVVGYDDTEDSSC
                     YIPPLTTIKQDFRLLGQTSVDRLLQLSQGQAVKGNQLLPVSLVKRKTTLAPNTQTASPR
                     ALADSLMQLARQVSRLESGQ"
     promoter        complement(3099..3176)
                     /gene="lacI"
                     /label=lacI promoter
     promoter        3485..3503
                     /label=T7 promoter
                     /note="promoter for bacteriophage T7 RNA polymerase"
     protein_bind    3504..3528
                     /label=lac operator
                     /bound_moiety="lac repressor encoded by lacI"
                     /note="The lac repressor binds to the lac operator to
                     inhibit transcription in E. coli. This inhibition can be
                     relieved by adding lactose or
                     isopropyl-beta-D-thiogalactopyranoside (IPTG)."
     RBS             3541..3563
                     /note="efficient ribosome binding site from bacteriophage
                     T7 gene 10 (Olins and Rangwala, 1989)"
     CDS             3573..3617
                     /codon_start=1
                     /product="peptide tag that allows for enzymatic
                     biotinylation"
                     /label=AviTag(TM)
                     /translation="GLNDIFEAQKIEWHE"
     CDS             3627..4283
                     /codon_start=1
                     /product="glutathione S-transferase from Schistosoma
                     japonicum"
                     /label=GST
                     /translation="MSPILGYWKIKGLVQPTRLLLEYLEEKYEEHLYERDEGDKWRNKK
                     FELGLEFPNLPYYIDGDVKLTQSMAIIRYIADKHNMLGGSPKERAEISMLEGAVLDIRY
                     GVSRIAYSKDFETLKVDFLSKLPEMLKMFEDRLSHKTYLNGDHVTHPDFMLYDALDVVL
                     YMDPMSLDAFPKLVSFKKRIEAIPQIDKYLKSSKYIAWPLQGWQATFGGGDHPPK"
     CDS             4305..4322
                     /codon_start=1
                     /product="6xHis affinity tag"
                     /label=6xHis
                     /translation="HHHHHH"
     terminator      4389..4436
                     /label=T7 terminator
                     /note="transcription terminator for bacteriophage T7 RNA
                     polymerase"
     rep_origin      4473..4928
                     /direction=RIGHT
                     /label=f1 ori
                     /note="f1 bacteriophage origin of replication; arrow
                     indicates direction of (+) strand synthesis"
     CDS             complement(5020..5835)
                     /codon_start=1
                     /gene="aph(3')-Ia"
                     /product="aminoglycoside phosphotransferase"
                     /label=KanR
                     /note="confers resistance to kanamycin in bacteria or G418
                     (Geneticin(R)) in eukaryotes"
                     /translation="MSHIQRETSCSRPRLNSNMDADLYGYKWARDNVGQSGATIYRLYG
                     KPDAPELFLKHGKGSVANDVTDEMVRLNWLTEFMPLPTIKHFIRTPDDAWLLTTAIPGK
```

```
                     TAFQVLEEYPDSGENIVDALAVFLRRLHSIPVCNCPFNSDRVFRLAQAQSRMNNGLVDA
                     SDFDDERNGWPVEQVWKEMHKLLPFSPDSVVTHGDFSLDNLIFDEGKLIGCIDVGRVGI
                     ADRYQDLAILWNCLGEFSPSLQKRLFQKYGIDNPDMNKLQFHLMLDEFF"
ORIGIN
        1 ttgagatcct ttttttctgc gcgtaatctg ctgcttgcaa acaaaaaaac caccgctacc
       61 agcggtggtt tgtttgccgg atcaagagct accaactctt tttccgaagg taactggctt
      121 cagcagagcg cagataccaa atactgtcct tctagtgtag ccgtagttag gccaccactt
      181 caagaactct gtagcaccgc ctacatacct cgctctgcta atcctgttac cagtggctgc
      241 tgccagtggc gataagtcgt gtcttaccgg gttggactca agacgatagt taccggataa
      301 ggcgcagcgg tcgggctgaa cggggggttc gtgcacacag cccagcttgg agcgaacgac
      361 ctacaccgaa ctgagatacc tacagcgtga gctatgagaa agcgccacgc ttcccgaagg
      421 gagaaaggcg gacaggtatc cggtaagcgg cagggtcgga acaggagagc gcacgaggga
      481 gcttccaggg ggaaacgcct ggtatcttta tagtcctgtc gggtttcgcc acctctgact
      541 tgagcgtcga tttttgtgat gctcgtcagg ggggcggagc ctatggaaaa acgccagcaa
      601 cgcggccttt ttacggttcc tggccttttg ctggcctttt gctcacatgt tctttcctgc
      661 gttatcccct gattctgtgg ataaccgtat taccgccttt gagtgagctg ataccgctcg
      721 ccgcagccga acgaccgagc gcagcgagtc agtgagcgag gaagcggaag agcgcctgat
      781 gcggtatttt ctccttacgc atctgtgcgg tatttcacac cgcaatggtg cactctcagt
      841 acaatctgct ctgatgccgc atagttaagc cagtatacac tccgctatcg ctacgtgact
      901 gggtcatggc tgcgccccga cacccgccaa cacccgctga cgcgccctga cgggcttgtc
      961 tgctcccggc atccgcttac agacaagctg tgaccgtctc cgggagctgc atgtgtcaga
     1021 ggttttcacc gtcatcaccg aaacgcgcga ggcagctgcg gtaaagctca tcagcgtggt
     1081 cgtgaagcga ttcacagatg tctgcctgtt catccgcgtc cagctcgttg agtttctcca
     1141 gaagcgttaa tgtctggctt ctgataaagc gggccatgtt aagggcggtt ttttcctgtt
     1201 tggtcactga tgcctccgtg taagggggat ttctgttcat gggggtaatg ataccgatga
     1261 aacgagagag gatgctcacg atacgggtta ctgatgatga acatgcccgg ttactggaac
     1321 gttgtgaggg taaacaactg gcggtatgga tgcggcggga ccagagaaaa atcactcagg
     1381 gtcaatgcca gcgcttcgtt aatacagatg taggtgttcc acagggtagc cagcagcatc
     1441 ctgcgatgca gatccggaac ataatggtgc agggcgctga cttccgcgtt tccagacttt
     1501 acgaaacacg gaaaccgaag accattcatg ttgttgctca ggtcgcagac gttttgcagc
     1561 agcagtcgct tcacgttcgc tcgcgtatcg gtgattcatt ctgctaacca gtaaggcaac
     1621 cccgccagcc tagccgggtc ctcaacgaca ggagcacgat catgcgcacc cgtggggccg
     1681 ccatgccggc gataatggcc tgcttctcgc cgaaacgttt ggtggcggga ccagtgacga
     1741 aggcttgagc gagggcgtgc aagattccga ataccgcaag cgacaggccg atcatcgtcg
     1801 cgctccagcg aaagcggtcc tcgccgaaaa tgacccagag cgctgccggc acctgtccta
     1861 cgagttgcat gataaagaag acagtcataa gtgcggcgac gatagtcatg ccccgcgccc
     1921 accggaagga gctgactggg ttgaaggctc tcaagggcat cggtcgagat cccggtgcct
     1981 aatgagtgag ctaacttaca ttaattgcgt tgcgctcact gcccgctttc cagtcgggaa
     2041 acctgtcgtg ccagctgcat taatgaatcg gccaacgcgc ggggagaggc ggtttgcgta
     2101 ttgggcgcca gggtggtttt tcttttcacc agtgagacgg gcaacagctg attgcccttc
     2161 accgcctggc cctgagagag ttgcagcaag cggtccacgc tggtttgccc cagcaggcga
     2221 aaatcctgtt tgatggtggt taacggcggg atataacatg agctgtcttc ggtatcgtcg
     2281 tatcccacta ccgagatatc cgcaccaacg cgcagcccgg actcggtaat ggcgcgcatt
     2341 gcgcccagcg ccatctgatc gttggcaacc agcatcgcag tgggaacgat gccctcattc
     2401 agcatttgca tggtttgttg aaaaccggac atggcactcc agtcgccttc ccgttccgct
     2461 atcggctgaa tttgattgcg agtgagatat ttatgccagc cagccagacg cagacgcgcc
     2521 gagacagaac ttaatgggcc cgctaacagc gcgatttgct ggtgacccaa tgcgaccaga
     2581 tgctccacgc ccagtcgcgt accgtcttca tgggagaaaa taatactgtt gatgggtgtc
     2641 tggtcagaga catcaagaaa taacgccgga acattagtgc aggcagcttc cacagcaatg
     2701 gcatcctggt catccagcgg atagttaatg atcagcccac tgacgcgttg cgcgagaaga
     2761 ttgtgcaccg ccgctttaca ggcttcgacg ccgcttcgtt ctaccatcga caccaccacg
     2821 ctggcaccca gttgatcggc gcgagattta atcgccgcga caatttgcga cggcgcgtgc
     2881 agggccagac tggaggtggc aacgccaatc agcaacgact gtttgcccgc cagttgttgt
     2941 gccacgcggt tgggaatgta attcagctcc gccatcgccg cttccacttt ttcccgcgtt
     3001 ttcgcagaaa cgtggctggc ctggttcacc acgcgggaaa cggtctgata agagacaccg
     3061 gcatactctg cgacatcgta taacgttact ggtttcacat tcaccaccct gaattgactc
     3121 tcttccgggc gctatcatgc cataccgcga aaggttttgc gccattcgat ggtgtccggg
     3181 atctcgacgc tctcccttat gcgactcctg cattaggaag cagcccagta gtaggttgag
     3241 gccgttgagc accgccgccg caaggaatgg tgcatgcaag gagatggcgc ccaacagtcc
     3301 cccggccacg gggcctgcca ccatacccac gccgaaacaa gcgctcatga gcccgaagtg
     3361 gcgagcccga tcttccccat cggtgatgtc ggcgatatag gcgccagcaa ccgcacctgt
     3421 ggcgccggtg atgccggcca cgatgcgtcc ggcgtagagg atcgagatct cgatcccgcg
     3481 aaattaatac gactcactat aggggaattg tgagcggata acaattcccc tctaggatcc
```

```
3541 tttgtttaac tttaagaagg agatatacaa tgggcctgaa cgacatcttc gaggctcaga
3601 aaatcgaatg gcacgaacgc ggatccatgt cccctatact aggttattgg aaaattaagg
3661 gccttgtgca acccactcga cttcttttgg aatatcttga agaaaaatat gaagagcatt
3721 tgtatgagcg cgatgaaggt gataaatggc gaaacaaaaa gtttgaattg ggtttggagt
3781 ttcccaatct tccttattat attgatggtg atgttaaatt aacacagtct atggccatca
3841 tacgttatat agctgacaag cacaacatgt tgggtggttc tccaaaagag cgtgcagaga
3901 tttcaatgct tgaaggagcg gttttggata ttagatacgg tgtttcgaga attgcatata
3961 gtaaagactt tgaaactctc aaagttgatt ttcttagcaa gctacctgaa atgctgaaaa
4021 tgttcgaaga tcgtttatct cataaaacat atttaaatgg tgatcatgta acccatcctg
4081 acttcatgtt gtatgacgct cttgatgttg ttttatacat ggacccaatg tccctggatg
4141 cgttcccaaa attagtttct tttaaaaaac gtattgaagc tatcccacaa attgataagt
4201 acttgaaatc cagcaagtat atagcatggc ctttgcaggg ctggcaagcc acgtttggtg
4261 gtggcgacca tcctccaaaa tagaagcttg cggccgcact cgagcaccac caccaccacc
4321 actgagatcc ggctgctaac aaagcccgaa aggaagctga gttggctgct gccaccgctg
4381 agcaataact agcataaccc cttggggcct ctaaacgggt cttgaggggt tttttgctga
4441 aaggaggaac tatatccgga ttggcgaatg ggacgcgccc tgtagcggcg cattaagcgc
4501 ggcgggtgtg gtggttacgc gcagcgtgac cgctacactt gccagcgccc tagcgcccgc
4561 tcctttcgct ttcttccctt cctttctcgc cacgttcgcc ggctttcccc gtcaagctct
4621 aaatcggggg ctccctttag ggttccgatt tagtgcttta cggcacctcg accccaaaaa
4681 acttgattag ggtgatggtt cacgtagtgg gccatcgccc tgatagacgg tttttcgccc
4741 tttgacgttg gagtccacgt tctttaatag tggactcttg ttccaaactg gaacaacact
4801 caaccctatc tcggtctatt cttttgattt ataagggatt ttgccgattt cggcctattg
4861 gttaaaaaat gagctgattt aacaaaaatt taacgcgaat tttaacaaaa tattaacgct
4921 tacaatttag gtggcacttt tcggggaaat gtgcgcggaa cccctatttg tttatttttc
4981 taaatacatt caaatatgta tccgctcatg aattaattct tagaaaaact catcgagcat
5041 caaatgaaac tgcaatttat tcatatcagg attatcaata ccatattttt gaaaaagccg
5101 tttctgtaat gaaggagaaa actcaccgag gcagttccat aggatggcaa gatcctggta
5161 tcggtctgcg attccgactc gtccaacatc aatacaacct attaatttcc cctcgtcaaa
5221 aataaggtta tcaagtgaga aatcaccatg agtgacgact gaatccggtg agaatggcaa
5281 aagtttatgc atttctttcc agacttgttc aacaggccag ccattacgct cgtcatcaaa
5341 atcactcgca tcaaccaaac cgttattcat tcgtgattgc gcctgagcga gacgaaatac
5401 gcgatcgctg ttaaaaggac aattacaaac aggaatcgaa tgcaaccggc gcaggaacac
5461 tgccagcgca tcaacaatat tttcacctga atcaggatat tcttctaata cctggaatgc
5521 tgttttcccg gggatcgcag tggtgagtaa ccatgcatca tcaggagtac ggataaaatg
5581 cttgatggtc ggaagaggca taaattccgt cagccagttt agtctgacca tctcatctgt
5641 aacatcattg gcaacgctac ctttgccatg tttcagaaac aactctggcg catcgggctt
5701 cccatacaat cgatagattg tcgcacctga ttgcccgaca ttatcgcgag cccatttata
5761 cccatataaa tcagcatcca tgttggaatt taatcgcggc ctagagcaag acgtttcccg
5821 ttgaatatgg ctcataacac cccttgtatt actgtttatg taagcagaca gttttattgt
5881 tcatgaccaa aatcccttaa cgtgagtttt cgttccactg agcgtcagac cccgtagaaa
5941 agatcaaagg atcttc
//
```
