## Supplemental Video for "CMG2 interaction with actin is required for growth factor-induced chemotaxis in endothelial cells"

### Slide 1
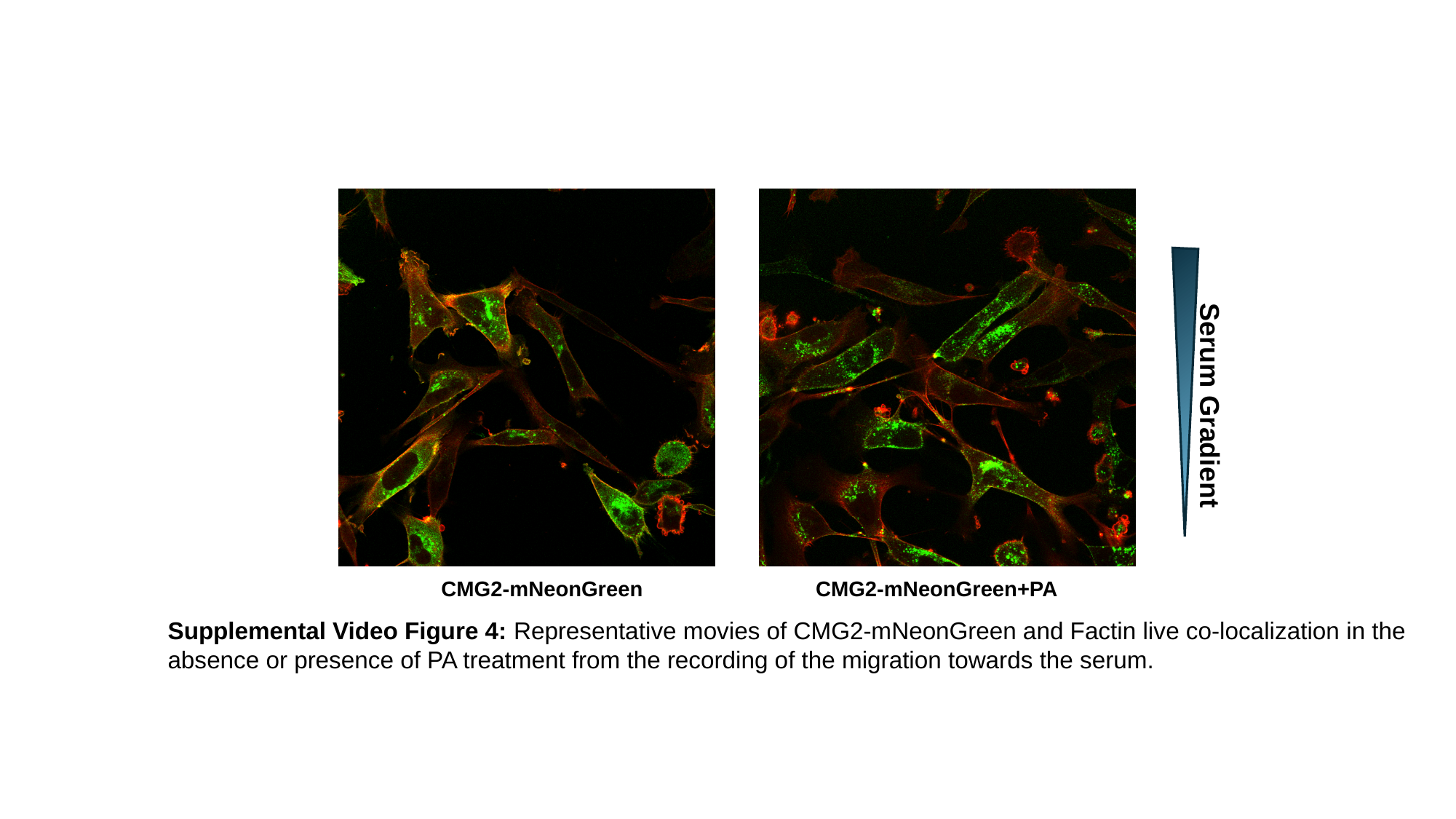

Serum Gradient
CMG2-mNeonGreen
CMG2-mNeonGreen+PA
Supplemental Video Figure 4: Representative movies of CMG2-mNeonGreen and Factin live co-localization in the
absence or presence of PA treatment from the recording of the migration towards the serum.
